# Paths and Pathways that Generate Cell-Type Heterogeneity and Developmental Progression in Hematopoiesis

**DOI:** 10.1101/2021.02.11.430681

**Authors:** Juliet R. Girard, Lauren M. Goins, Dung M. Vuu, Mark S. Sharpley, Carrie M. Spratford, Shreya R. Mantri, Utpal Banerjee

## Abstract

Mechanistic studies of *Drosophila* lymph gland hematopoiesis are limited by the availability of cell-type specific markers. Using a combination of bulk RNA-Seq of FACS-sorted cells, single cell RNA-Seq and genetic dissection, we identify new blood cell subpopulations along a developmental trajectory with multiple paths to mature cell types. This provides functional insights into key developmental processes and signaling pathways. We highlight metabolism as a driver of development, show that graded Pointed expression allows distinct roles in successive developmental steps, and that mature crystal cells specifically express an alternate isoform of Hypoxia-inducible factor (Hif/Sima). Mechanistically, the Musashi-regulated protein Numb facilitates Sima-dependent non-canonical, while inhibiting canonical, Notch signaling. Broadly, we find that prior to making a fate choice, a progenitor selects between alternative, biologically relevant, transitory states allowing smooth transitions reflective of combinatorial expressions rather than stepwise binary decisions. Increasingly, this view is gaining support in mammalian hematopoiesis.

## Introduction

The *Drosophila* lymph gland is the major hematopoietic organ that develops during the larval stages for the purpose of providing blood cells during later pupal/adult periods (reviewed in Banerjee et al. 2019). Hematopoietic function for the larva itself is largely provided by a separate set of sessile or circulating blood cells outside of the lymph gland (reviewed in Letourneau et al. 2016). The only time the lymph gland provides blood cells to the circulating larval hemolymph is if the larva faces a stress or immune challenge.

The first appearance of the lymph gland tissue as a separate entity from the dorsal mesoderm is during embryonic stages 11-13. Three distinct segments (T1-T3) contribute to the formation of the primary lobes of the lymph gland. Cells from the anterior T1 and T2 segments coalesce to form the lymph gland proper, while cells from T3 combine with the rest, but maintain their separate identity as the PSC (Posterior Signaling Center). The latter cells are not hematopoietic in function but provide a supporting role for the blood progenitors much as the stromal cells do in mammalian systems (reviewed in Crane et al. 2017). The PSC is marked by its specific expression of *Antp* (Mandal et al. 2007) and by high expression of *knot* (*kn*)/*collier* (*col)*, which encodes an orthologue of EBF (Early B-Cell factor) (Crozatier et al. 2004). The above description is for the primary (or anterior) lymph gland lobes. Further posterior to them are the secondary (or posterior) lobes and pericardial cells. The primary/anterior lobes of the lymph gland display the highest hematopoietic activity during normal larval development. For that reason, we concentrate exclusively on the hematopoietic events of the primary lobes in this paper.

Invertebrates did not evolve a lymphoid system for acquired immunity. Accordingly, *Drosophila* blood cells are all similar in function to cells of the vertebrate myeloid lineage. Of these, the most predominant class called plasmatocytes share a monophyletic connection with vertebrate macrophages, and the two serve similar functions. Classic studies identified three morphologically distinct mature hemocyte types based on their ultrastructure, cytochemistry, and function (Shrestha and Gateff 1982). In addition to the highly abundant plasmatocytes (95% of all hemocytes), a minor (4-5%) but incredibly important class is represented by crystal cells. The third class, lamellocytes (<1%), are not usually present in healthy animals but are induced upon specific stresses, particularly during wasp parasitization (reviewed in Letourneau et al. 2016). Later work developed cell type-specific antibodies against surface antigens (Kurucz et al. 2007), enhancer traps/GAL4 lines (Rodriguez et al. 1996; Braun et al. 1997), and specific transcription factors that specify hemocyte fate.

Plasmatocytes, marked by their expression of *Hemolectin* (*Hml*), *Peroxidasin* (*Pxn*) and *NimC1*, participate in engulfment of microbes and apoptotic cells, and they produce extracellular matrix proteins (Fessler and Fessler 1989; Tepass et al. 1994; Franc et al. 1996). Crystal cells (CCs), named for their crystalline inclusions of pro-phenoloxidase enzymes (PPO1 and PPO2), are necessary for melanization, blood clot formation, immunity against bacterial infections, and to help mitigate hypoxic stress caused by CO_2_/O_2_ imbalance (Rämet et al. 2002; Galko and Krasnow 2004; Binggeli et al. 2014; Dudzic et al. 2015; Cho et al. 2018). The transcription factor Lozenge (Lz) cooperates with Notch signaling to express a number of target genes (such as *hindsight*/*pebbled*) to specify CCs (Lebestky et al. 2000; Duvic et al. 2002) while the Sima (vertebrate HIF-1alpha) protein is required for their maintenance (Mukherjee et al. 2011). The orthologue of Lz in mammals is RUNX1, with broad hematopoietic function at many developmental stages, and RUNX1 is often dysregulated in acute myeloid leukemias (de Bruijn and Speck 2004; Ito 2004).

The primary lobe cells are physically arranged into zones defined by the cell types that occupy them. Blood progenitors, marked by *dome* and *Tep4*, reside in the medullary zone (MZ) while differentiating hemocytes (expressing plasmatocyte and CC markers) occupy the cortical zone (CZ) (Jung et al. 2005). A third zone named the intermediate zone (IZ) is the site of the intermediate progenitors (IPs) (Krzemien et al. 2010) that is explored in detail in this, and in a companion paper (Spratford et al. 2020). Prior work describing the zonation of the lymph gland also noted a small population of cells near the dorsal vessel with characteristics of a pre-progenitor (Jung et al. 2005; Dey et al. 2016; Tiwari et al. 2020), but the rest of the MZ was considered to be fairly homogeneous. More recent studies from multiple laboratories, point to considerable heterogeneity and complexity within the progenitor population (reviewed in Banerjee et al. 2019). Particularly noteworthy is the functional distinction into a Hh-sensitive and a Hh-resistant group of progenitors within the MZ (Baldeosingh et al. 2018). Many different signaling modalities coalesce to control differentiation and maintenance of progenitors during hematopoiesis including direct cell to cell interactions (e.g. Serrate/Notch), interzonal communication (e.g. Hedgehog), and systemic signals (e.g. olfactory and nutritional) (Lebestky et al. 2003; Crozatier et al. 2004; Mandal et al. 2007; Shim et al. 2012; Shim et al. 2013; Ferguson and Martinez-Agosto 2014). Manipulation of such pathways using mutations of their key components impacts blood cell development. For example, loss of nutritional signals by starvation or blocking olfaction greatly impacts the balance between progenitors and differentiated cell types.

A rather important type of interzonal signaling mechanism relevant to this paper involves multiple cell types across the zones. In brief, progenitors are maintained not only through niche-derived signals but also through a signaling relay mediated by the differentiated cells. This backward signal is named the Equilibrium Signal (Mondal et al. 2011; Mondal et al. 2014). In this process, Pvf1 (PDGF- and VEGF-related factor 1) produced by the PSC trans-cytoses through the MZ to bind its receptor Pvr (PDGF/VEGF receptor), expressed in the CZ. This initiates a STAT-dependent but JAK-independent signaling cascade that ultimately leads to the secretion of the extracellular enzyme ADGF-A (adenosine deaminase-related growth factor A). This enzyme breaks down adenosine, preventing its mitogenic signal and proliferation of MZ progenitors. Together with the niche signal, this backward signal allows a balance between progenitor and differentiated cell types (Mondal et al. 2011; Mondal et al. 2014). The genetic studies broadly implicated the CZ cells as originators of this backward signal. Finer analysis, afforded by cell-separated bulk and single cell RNA-Seq in this study, allows us to attribute this role to a smaller and more specific subset of cells.

RNA-Seq has been used recently as a technique to study *Drosophila* blood cells (Cattenoz et al. 2020; Cho et al. 2020; Fu et al. 2020; Ramond et al. 2020; Tattikota et al. 2020). However, several important distinctions make this current work broadly relevant to developmental biology. The first four of the cited studies, analyze circulating blood cells that have a completely different developmental profile than the lymph gland. Cho et al. (2020) utilize the lymph gland and some of the similarities and many differences between the conclusions are directly compared and contrasted later. For our current study it proved critical to utilize a complex strategy that combines several techniques in order to arrive at any of the mechanistic aspects of our conclusions including a full analysis of intermediate zone cells, identification of two transitional populations, role of metabolism, functional aspects of transcriptional regulation, as well as the unique mechanism of crystal cell maturation by the combined functions of a novel and specific isoform of Sima, Notch, Numb and Musashi. Bulk RNA-Seq on FACS sorted populations of dissociated cells bearing cell-specific markers provides enough depth of sequencing to identify alternate isoforms. When combined with single cell RNA-Seq, these results produce a well-articulated developmental trajectory that we verify and establish by genetic means.

This combination of molecular genetics and whole genome approaches makes it clear that hematopoietic cells are far more heterogeneous and diverse than previously realized by genetics alone, and helps shift our view of hematopoiesis from being a series of discrete steps to a more continuous journey along multiple paths. The multiplicity in layers of decision points creates these new routes, which can each lead to a distinct differentiated endpoint, or, alternatively, follow their parallel trajectories to a single final outcome.

## Results

### Bulk RNA-Seq analysis of zonal patterning within the lymph gland

To better understand the distribution of gene-expression patterns in different lymph gland zones, we utilize a combination of established, directly driven, reporter constructs that mark the medullary zone, MZ (*dome^MESO^* enhancer driven nuclear EGFP) as well as the cortical zone, CZ (*Hml* enhancer driven nuclear DsRed). The absence of any GAL4 driver allows simultaneous visualization and manipulation of different cell types. Lymph glands from these marked third instar larvae are dissected and the primary lobes are separated from the rest of the lymph gland. The primary lobe cells are then dissociated and sorted by FACS (fluorescence-activated cell sorting). We identify two marked cell types that are singly positive for either marker, representing the cells of the MZ and the CZ (Figure 1A-B). Additionally, we identify a distinct cell population that is positive for both markers (Figure 1A-B). These represent intermediate progenitors (IPs) belonging to an Intermediate Zone (IZ) between the MZ and the CZ (Krzemien et al. 2010). The IPs were initially identified as transitioning cells at the distal edge of the MZ that simultaneously express a *dome* reporter and Peroxidasin (Pxn), an alternative to *Hml* as a CZ marker (Sinenko et al. 2009; Krzemien et al. 2010). Here, using direct drivers and nuclear fluorescent proteins, we readily identify this double positive population both in the intact lymph gland (Figure 1A-A’) and in dissociated cells (Figure 1B). We find that IPs express lower levels of the markers EGFP and DsRed than in MZ and CZ cells, respectively (Figure 1B). The single and double positive cells are separable by FACS (Figure 1-figure supplement 1A) and are then used in bulk RNA-Seq experiments. A fourth population that is double negative for both markers is also detected in the FACS sorted populations. We have not characterized these cells in detail, as they seem to arise for a multitude of reasons, some interesting, such as belonging to the PSC, which is not marked in these tissues, but is explored in the single cell RNA-Seq (scRNA-Seq) experiments, and some that are uninteresting such as unintended loss of fluorescence due to bleaching or contamination with non-hematopoietic tissue such as the heart and some pericardial cells.

**Figure 1:**
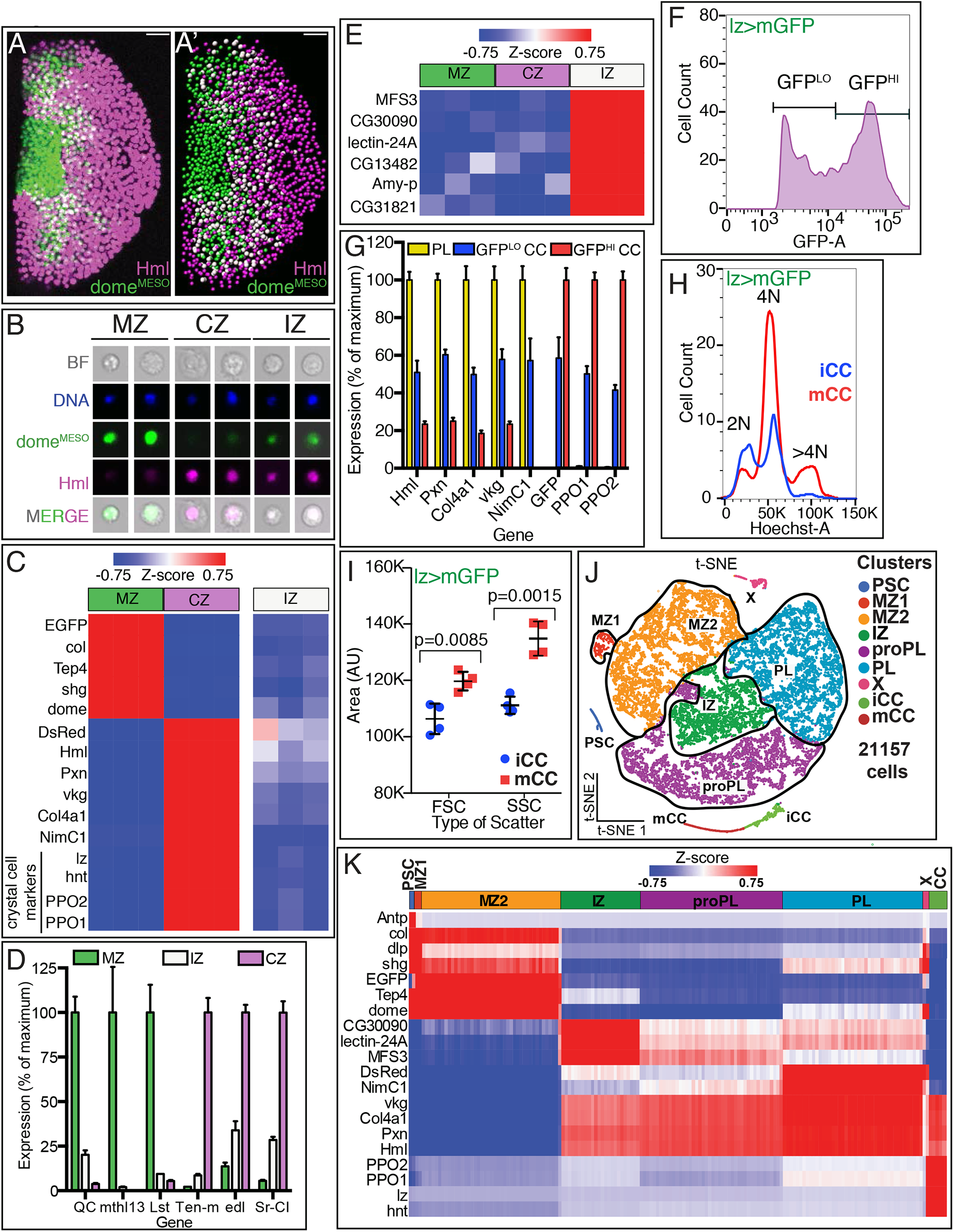
Analysis of subzonal patterning by RNA-Seq of the lymph gland. (**A-E**) Genotype of glands used is *dome^MESO^-GFP.nls, Hml^Δ^-DsRed.nls*. GFP (*dome*) is in green and DsRed (*Hml*) is in magenta. **(A)** Confocal image shows the zonal pattern of an early third instar lymph gland. Progenitors of the medullary zone (MZ, green), differentiated cells of the cortical zone (CZ, magenta), and cells in the intermediate zone (IZ, white due to colocalization of green and magenta) are seen as distinct cell types. **(A’)** Computer rendering of the confocal image shown in **(A).** Nuclei are pseudo-colored based on their fluorescence: MZ (green), CZ (magenta) and IZ double positive cells (white). **(A-A’)** Scale bars, 20 µm. **(B)** Images of individual dissociated lymph gland cells. Brightfield (BF), DAPI/DNA (blue), MZ cells (green), CZ cells (magenta), and IZ cells (green (weak) and magenta). **(C-E)** Gene expression from bulk RNA-Seq analysis of dissociated and sorted cells in three biological replicates. **(C)** High levels of hallmark genes representative of the MZ progenitors: (*dome^MESO^-)EGFP*, *col* (*collier*), *Tep4*, *shg* (*shotgun*; *E-Cad*), *dome* (*domeless*); CZ plasmatocytes: (*Hml^Δ^-)DsRed, Hml* (*Hemolectin*), *Pxn* (*Peroxidasin*), *vkg* (*viking*), *Col4a1*, and *NimC1*; and CZ crystal cells: *lz* (*lozenge*), *hnt* (*hindsight*/*pebbled*), *PPO2*, and *PPO1* are seen in the corresponding populations. IZ cells show low to moderate levels of most MZ and CZ hallmark genes. **(D)** Newly identified zone-enriched genes for MZ include *QC*, *mthl13*, and *Lst*. For CZ, these include *Ten-m*, *edl*, and *Sr-Cl*. In general, IZ cells show low to moderate levels of these MZ and CZ-specific markers. **(E)** IZ cells show high expression of the newly identified IZ-enriched marker genes *MFS3*, *CG30090*, *lectin-24A, CG13482*, *Amy-p*, and *CG31821*. MZ and CZ cells show low levels of these genes. **(F-I)** Flow cytometric analysis and bulk RNA-Seq of dissociated crystal cell-marked lymph gland cells. Genotype: *lz-GAL4*, *UAS-mGFP*; *Hml^Δ^-DsRed.nls*. Crystal cells expressing *lz-GAL4* are marked by GFP and are distinguishable from plasmatocytes expressing DsRed (*lz>mGFP* negative, *Hml^Δ^-DsRed.nls* positive). **(F)** Analysis of *lz>mGFP* fluorescence shows two distinct populations of crystal cells (CC), those with low GFP (GFP^LO^) and those with high GFP (GFP^HI^). GFP^LO^ CCs are referred to as iCCs and GFP^HI^ CCs are referred to as mCCs (see text). **(G)** Gene expression analysis based on bulk RNA-Seq. Plasmatocytes (PL) show high expression of *Hml, Pxn, Col4a1, vkg, and NimC1* and no expression of *(lz>)mGFP*, *PPO1* and *PPO2*. GFP^LO^ CCs (iCCs) show moderate levels of both plasmatocyte and crystal cell specific genes. GFP^HI^ CCs (mCCs) show high *PPO1* and *PPO2*, no expression of *NimC1* and low expression of other PL markers. **(H-I)** Flow cytometric analysis of iCCs and mCCs. **(H)** DNA content (Hoechst-A) analysis shows that iCCs have 2N or 4N DNA content, while mCCs have a significant number of cells with >4N DNA content indicative of endoreplication. **(I)** Quantitation of data from four individual experiments. mCCs have higher mean FSC-A (cell size) and mean SSC-A (cellular complexity) values than iCCs. (p values shown are from t-test) **(J-K)** Single cell RNA-Seq analysis of dissociated cells from *dome^MESO^-GFP.nls, Hml^Δ^-DsRed.nls* lymph glands. **(J)** t-SNE visualization of graph-based clustering identifies 9 clusters: PSC (dark blue), MZ1 (red), MZ2 (orange), IZ (green), proPL (purple), PL (light blue), X (pink), iCC (light green), and mCC (dark red). **(K)** Expression analysis of hallmark genes shows enrichment in appropriate clusters. PSC (*Antp*, *col*, and *dlp*), MZ (*shg*, *EGFP*, *Tep4*, and *dome*), IZ (*CG30090*, *lectin-24A*, and *MFS3*), CZ (*DsRed*, *vkg*, *Col4a1*, *Pxn*, and *Hml*), mature plasmatocytes (*NimC1*), and crystal cells (*PPO2*, *PPO1*, *lz* and *hnt*).

For each bulk RNA-Seq sample, we dissect approximately 100 lymph glands from mid-third instar larvae (90-96 hours after egg lay (AEL) at 25°C), dissociate them, and sort the resulting cells using 4 gates (Figure 1-figure supplement 1A). Three biological replicates are analyzed for each sample and in the results reported here, approximately 11,000 genes meet our threshold criteria for transcript expression across the populations. Previously established markers, which we refer to as “hallmark genes”, such as *Tep4*, *dome*, *shg/E-Cad*, and *kn/collier,* (and *EGFP),* are detected in the MZ (Figure 1C). Similarly, *vkg*, *Col4a1*, *Hml*, *Pxn*, and *NimC1* (and *DsRed*) transcripts are enriched in the CZ (Figure 1C). In addition, we identify several novel markers that uniquely characterize the MZ (*QC*, *mthl13*, *Lst*) or the CZ (*Ten-m*, *edl*, *Sr-CI*) population (Figure 1D). Future genetic analysis will determine how these genes function differentially in their specified zones. The *Hml^Δ^-DsRed* population very clearly also contains crystal cells (CC), as the markers *lz*, *hnt* (*pebbled; peb*), *PPO1*, and *PPO2* are seen in this population (Figure 1C). *Hml-GAL4* expression in crystal cells has been previously reported (Goto et al. 2003), and indeed, in whole-mount stained lymph glands we find that nearly all crystal cells express low levels of *Hml^Δ^-DsRed* (Figure 1-figure supplement 1B). This low *Hml* is lost in crystal cells expressing very high Hnt.

IZ cells express low to moderate levels of many MZ and CZ markers, but importantly, we detect no expression of late differentiation markers such as *NimC1* or *PPO1/2*, which are characteristic of mature plasmatocytes and crystal cells, respectively (Figure 1C). Nor do they express significant levels of very early progenitor markers such as *Tep4* and *kn/collier* (Figure 1C). The fact that the IPs do not express either the earliest progenitor or the most mature differentiation markers provides further support to the idea that they are a transitional population between the MZ and the CZ. Importantly, transcriptomic analysis reveals that the IPs are, in themselves, a *bona fide* cell type as they are uniquely enriched in transcripts such as *MSF3*, *CG30090*, *lectin-24A, CG13482*, *Amy-p*, and *CG31821* as compared with the expression of these transcripts in either MZ or CZ (Figure 1E). The collective expression of these bulk RNA-Seq derived novel IZ markers proved crucial in specifying a cell as an IP in our subsequent single cell-Seq analysis.

The described bulk RNA-Seq contains a mixed CZ population of plasmatocytes and crystal cells (CCs). In order to sort crystal cells away from other *Hml*-positive cells, we use a genotype which combines *Hml^Δ^-DsRed.nls* with *lz>mGFP* (*lz-GAL4, UAS-mGFP*), in which crystal cells are identified by their expression of *lz* (Lebestky et al. 2000; Siddall et al. 2009). For this second bulk RNA-Seq, we use late wandering third instar larvae (93-117 hours AEL) at which stage crystal cells are more abundant than in the mid-third instar. All other conditions remain the same. When we gate and sort for *Hml^Δ^-DsRed.nls* and *lz>mGFP*, two clearly separable GFP populations become evident. One displays high GFP (GFP^HI^) and the other is found to be lower (GFP^LO^) (Figure 1F). A third population of DsRed-positive cells is GFP-negative and represents the plasmatocytes (Figure 1-figure supplement 1C). The identification of the *lz-*negative and *Hml-*positive population as plasmatocytes is further confirmed by their expression of the hallmark genes: *Hml*, *Pxn*, *Col4a1*, *vkg*, and *NimC1,* and because they do not express *PPO1* or *PPO2* (Figure 1G). Both GFP^HI^ and GFP^LO^ cells express *PPO1/2* and they are both crystal cell populations (Figure 1G).

GFP^HI^ cells express much higher levels of the known maturity markers *PPO1/2* than their GFP^LO^ CC counterparts (Figure 1G). This suggests that GFP^HI^ cells are a more mature CC (mCC) population and comparatively speaking, GFP^LO^ CCs are immature (iCC). Plasmatocyte-related genes are expressed at a higher level in iCCs than in mCCs (Figure 1G; Figure 1-figure supplement 1D). mCCs and iCCs express comparable levels of the pan-crystal cell marker *hnt* (Figure 1-figure supplement 1D). The expression of *lz* RNA is also only marginally different between the two populations (Figure 1-figure supplement 1D), although its surrogate, *lz>mGFP*, is readily distinguishable (Figure 1G), and is the basis for separating the two populations by FACS. Flow cytometric analysis reveals further interesting differences in the biological characteristics of the immature (iCC; GFP^LO^) and more mature (mCC; GFP^HI^) crystal cells. As expected, both mCC and iCC populations are represented by a mixture of cells with 2N and 4N DNA content (Figure 1H). Interestingly, however, a subset of mCCs, but not iCCs, exhibits >4N DNA content, indicative of endocycling (Figure 1H). Previous studies have also found evidence for endocycling in crystal cells (Krzemien et al. 2010; Terriente-Felix et al. 2013). Combined with DNA content analysis, these data suggest that endocycling is confined to the more mature, mCC subpopulation. We also find that the average forward scatter (FSC-A), a measure of cell size, and average side scatter (SSC-A), a measure of internal complexity, are higher in mCCs compared to iCCs (Figure 1I; Figure 1-figure supplement 1E-F). This suggests that mCCs are larger and more granular than iCCs. Taken together, these results show that mCCs and iCCs differ in their gene expression profiles and cellular properties, and that they represent two separate maturity states of crystal cells.

### Single cell RNA-Seq defines subzonal patterns within the lymph gland

Bulk RNA-Seq is a useful tool for identifying gene expression in a relatively large group of cells with previously established canonical biomarkers. Its usefulness is in that it provides a broad landscape for gene expression. However, in the case of the lymph gland, a finer analysis at a cellular level is warranted because of the considerable heterogeneity reported amongst the cells belonging to individual zones (Jung et al. 2005; Kurucz et al. 2007; Oyallon et al. 2016; Baldeosingh et al. 2018; Blanco-Obregon et al. 2020). We therefore use single cell RNA-Seq to complement and enhance the bulk RNA-Seq data and to characterize subpopulations of cells that reside within each zone. The same genetic background and similar developmental timing (90-93 hours AEL at 25° C) is used as in the bulk RNA-Seq experiments to facilitate comparison between the two approaches. Each single cell RNA-Seq (scRNA-Seq) sample utilizes 11 lymph glands to yield a concentrated cell suspension with high (85-90%) cell viability. Three identical biological replicates are processed in parallel and in the results presented here, the transcriptome of about 21,200 individual cells is determined using the 10x Genomics platform and analyzed using Partek Flow software (see Methods). Graph-based clustering analysis and t-distributed stochastic neighbor embedding (t-SNE) visualization in 2D and 3D are then performed.

This analysis predicts nine individual cell clusters within the lymph gland (Figure 1J). Each of the graph-based clusters is assigned its unique identity, in part, based on the presence of known zone-specific markers within the differentially expressed genes. The assigned names for these clusters are justified in later sections. The PSC and IZ are each represented by single clusters. We identify two clusters (MZ1 and MZ2) with progenitor characteristics. In addition to IZ, the data suggest a second transitional population, proPL, that straddles MZ2 and a plasmatocyte cluster, PL (Figure 1J; Video 1). Similar to the two CC clusters seen in bulk RNA-Seq analysis, the single major crystal cell cluster splits into two (iCC and mCC) upon subclustering (Figure 1J). All of the above clusters express their respective previously identified zone-specific genes (Figure 1K; see Supplementary file 1).

The cluster designated as “X” on the t-SNE exhibits high levels of mitosis and replication stress related genes. The PSC, CC, and X clusters are distinct enough from the rest to remain as islands distant from each other and the core group of the other cell populations. The similarities and gradual transitions between the rest of the cells (belonging to MZ1, MZ2, IZ, proPL, and PL) causes their clusters to be closely associated as a core group of neighbors on the t-SNE map (Figure 1J).

Trajectory and pseudotime analysis are used to map the time-line of progression of the identified heterogeneous population of cells through their multiple phases of maturity (Figure 2A-J). In this analysis, changes in gene expression are calculated as the distance of each cell from the beginning of the trajectory (the “pseudotime”). A “branched’’ trajectory is produced if multiple cellular outcomes or fates are possible. The branch points represent cellular decision points and “states” correspond to stretches bounded by the branch points. This trajectory analysis can be used to create further groupings within the major clusters. PSC is separate in developmental origin from the rest of the lymph gland (Mandal et al. 2007) and cluster X likely represents mitotic states of several distinct cell types and therefore, although these two populations are represented in the t-SNE, they are not included for the purpose of constructing the trajectory. We find that the lymph gland cells form a branched trajectory with a total of three branch points and seven states (Figure 2A-J). Mapping the states back onto the t-SNE allows visualization of individual paths between related clusters (Figure 2K-S).

**Figure 2:**
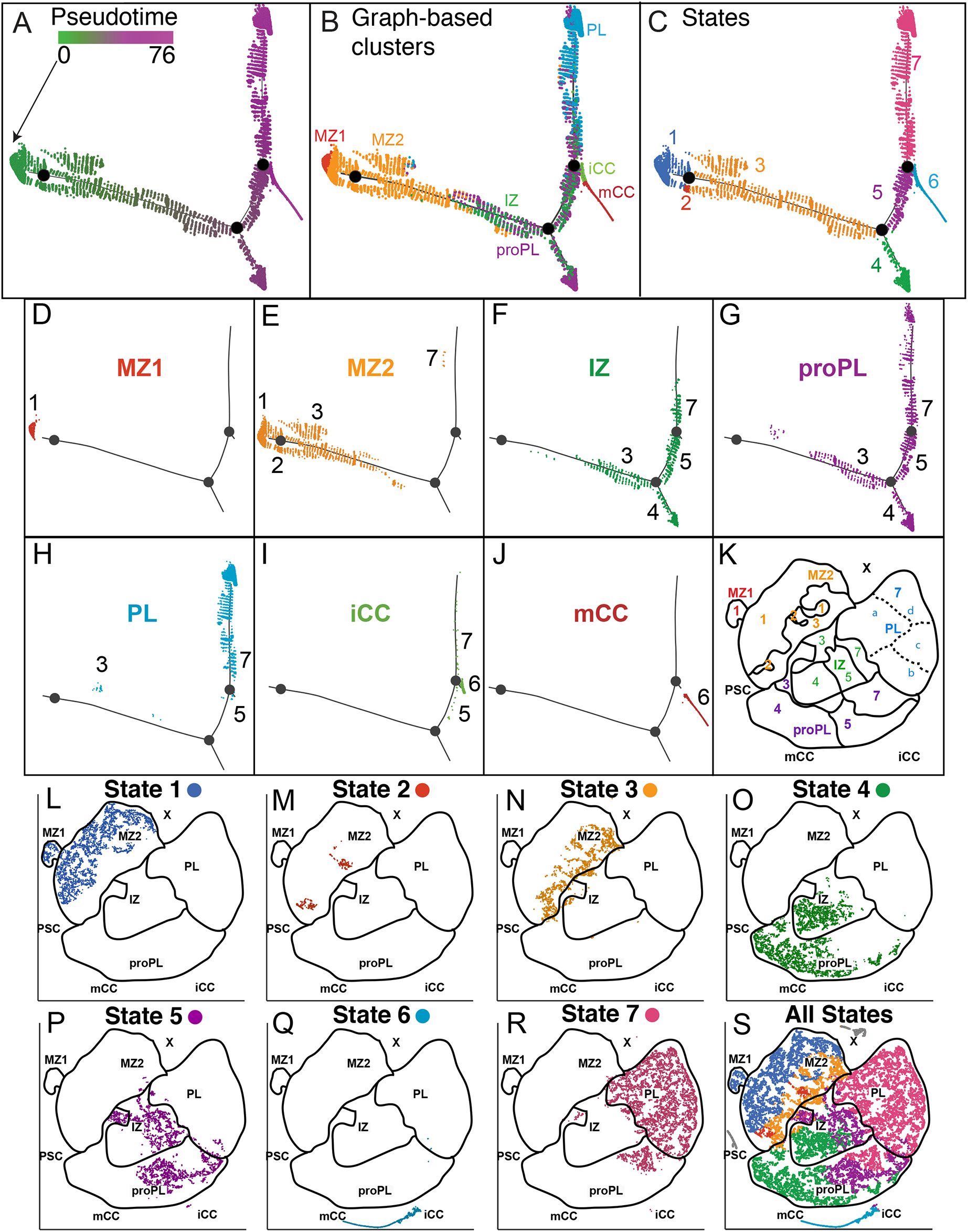
Developmental trajectory analysis of single cell RNA-Seq results using Monocle 2. **(A-J)** Trajectory diagram shows the progression of lymph gland cells from the earliest progenitors to the most mature cell types. **(A)** Pseudotime initiates (green) progressing in its temporal order to later (magenta) developmental stages. **(B)** Superposition of graph-based clusters onto the trajectory shows the initial appearance of MZ1 at the beginning of the trajectory, proceeding through MZ2, IZ/proPL and ultimately onto the differentiated cell types, iCC, mCC and PL. **(C)** Trajectory diagram showing 7 states, each separated between branch points of the trajectory. State 1 (early progenitors), State 2 (later progenitors), State 3 (transitioning cells belonging to progenitor, intermediate progenitor, and proplasmatocyte clusters), States 4 and 5 (differentiating cells belonging to intermediate progenitor and proplasmatocyte clusters), State 6 (terminal crystal cells), and State 7 (terminal plasmatocytes). **(D-J)** Trajectory diagrams split by individual graph-based clusters. **(D)** MZ1 progenitors are found at the beginning of the trajectory exclusively in state 1. **(E)** MZ2 progenitors are found at the beginning and middle of the trajectory and span states 1, 2, and 3, with a small number in state 7. **(F)** IZ cells are found in the middle of the trajectory in states 3, 4, 5, and 7. **(G)** proPL cells are found in the middle of the trajectory in states 3, 4, 5, and 7. **(H)** PL cells are found toward the end of the trajectory almost entirely in state 7, with very small numbers of cells in states 3 and 5. **(I)** iCCs are found towards the end of the trajectory, mostly on a unique branch (state 6), with a few cells found in states 5 and 7. **(J)** mCCs are located exclusively at the end of the trajectory in state 6. **(K-S)** Visualization of the trajectory “states” overlaid on the t-SNE plot. **(K)** A blank template of the t-SNE plot that outlines the graph-based clusters (indicated by their respective color) and the trajectory states (indicated by their respective number). **(L-S)** Cells are pseudo-colored by the state in which they are found on the trajectory. **(L)** State 1 contains all MZ1 cells (MZ1-1) as well as a portion of the MZ2 cells (MZ2-1) that cluster closest to MZ1 on the t-SNE. **(M)** State 2 contains MZ2 cells (MZ2-2) that border MZ2 cells in states 1 and 3. **(N)** State 3 contains multiple cell clusters (MZ-3, IZ-3, proPL-3, and PL-3) that are found at the border between these clusters on the t-SNE. This is consistent with their status as transitioning cells. **(O)** State 4 contains IZ and proPL cells primarily clustered on the left side of the t-SNE. **(P)** State 5 is made up of IZ and proPL cells clustered in the center of the t-SNE. **(Q)** State 6 is found exclusively on the crystal cell island in the t-SNE. **(R)** State 7 is primarily made up of IZ, proPL, and PL cells mapping to the right side of the t-SNE. **(S)** All states shown together on the t-SNE. PSC and X cells are colored grey as they were not used for trajectory analysis.

#### MZ clusters

The MZ is represented by the two major clusters MZ1 and MZ2 (Figure 1J) that express known hallmark genes as well as a number of genes that are newly identified as MZ-specific in the bulk RNA-Seq experiments (Figure 1K; Figure 2-figure supplement 1A-D). MZ1 cells are found entirely at the beginning of the trajectory at the earliest pseudotime (Figure 2A-C) and all MZ1 cells are contained within state 1 of the trajectory (thus named, MZ1-1; Figure 2D, L). MZ1-1 represents a small number of cells (1.3% of total cells; 4.8% of total MZ cells) with higher expression levels of many progenitor markers (e.g. *shg*, *col/kn*, *dome*, *Tep4*) than MZ2 (Figure 2-figure supplement 1E). On the t-SNE map, MZ1 is exclusively adjacent to MZ2 and is not near the differentiating CZ clusters (Figure 1J). As described later, gene expression in MZ1 has certain similarities to the PSC. The MZ1 cells are clearly the earliest progenitors at this stage of development and it is conceivable that they also have some signaling function.

MZ2 is placed in between MZ1 and IZ, proPL, and PL clusters (Figure 1J) and can be subdivided into three subclusters, MZ2-1, MZ2-2 and MZ2-3 based on their three trajectory states (Figure 2E, L-N). The majority of MZ2 progenitors are found in State 1 (MZ2-1, 68% of all MZ2 cells). A very small number of cells are found in State 2, (MZ2-2; 2% of MZ2). The MZ2-3 subcluster, in state 3, is of significant size (29% of MZ2 cells). The levels of the progenitor markers *Tep4* and *dome* are highest in MZ1-1 and decrease in MZ2-1, and decline even further in MZ2-3 (Figure 2-figure supplement 1F-G). Importantly, however, the *Tep4* and *dome* levels in MZ2-3 are still higher than those seen in the intermediate zone (Figure 2-figure supplement 1F-G).

To calculate an enrichment score for the collective expression level of a set of genes (e.g. those belonging to a single pathway or those expressed in subpopulations), we use AUCell analysis (Aibar et al. 2017). Using the expression set of a number of IZ-specific markers determined from the bulk RNA-Seq data (Figure 1E) in this manner, we find that the expression of the IZ enriched gene set is negligible in MZ1-1, increases stepwise from MZ2-1 to MZ2-2 and MZ2-3 and is further higher in IZ, but then decreases in plasmatocyte populations (Figure 2-figure supplement 1H and Figure 2-figure supplement 3A). This contrasts with the opposite trend seen for *dome* and *Tep4.* Based on gene expression patterns, we propose that MZ2-2 and MZ2-3 are similar and represent a more mature population within MZ2. Together they represent a transitional group of cells that mature to IZ, proPL, and PL.

There are several commonalities between MZ1 and MZ2, justifying the classification of both as MZs. However, there are also very significant differences between the two. As for similarities, these two subpopulations share 43 of the 241 differentially expressed MZ genes (Supplementary file 1). In addition to the canonical markers, both MZ1 and MZ2 are enriched in genes that are involved in ribosome biogenesis (more details in Figure 3) and glutathione metabolism/glutathione transferase activity (e.g. *Gsts D3*, *E6*, *E9*, and *O3*).

**Figure 3.**
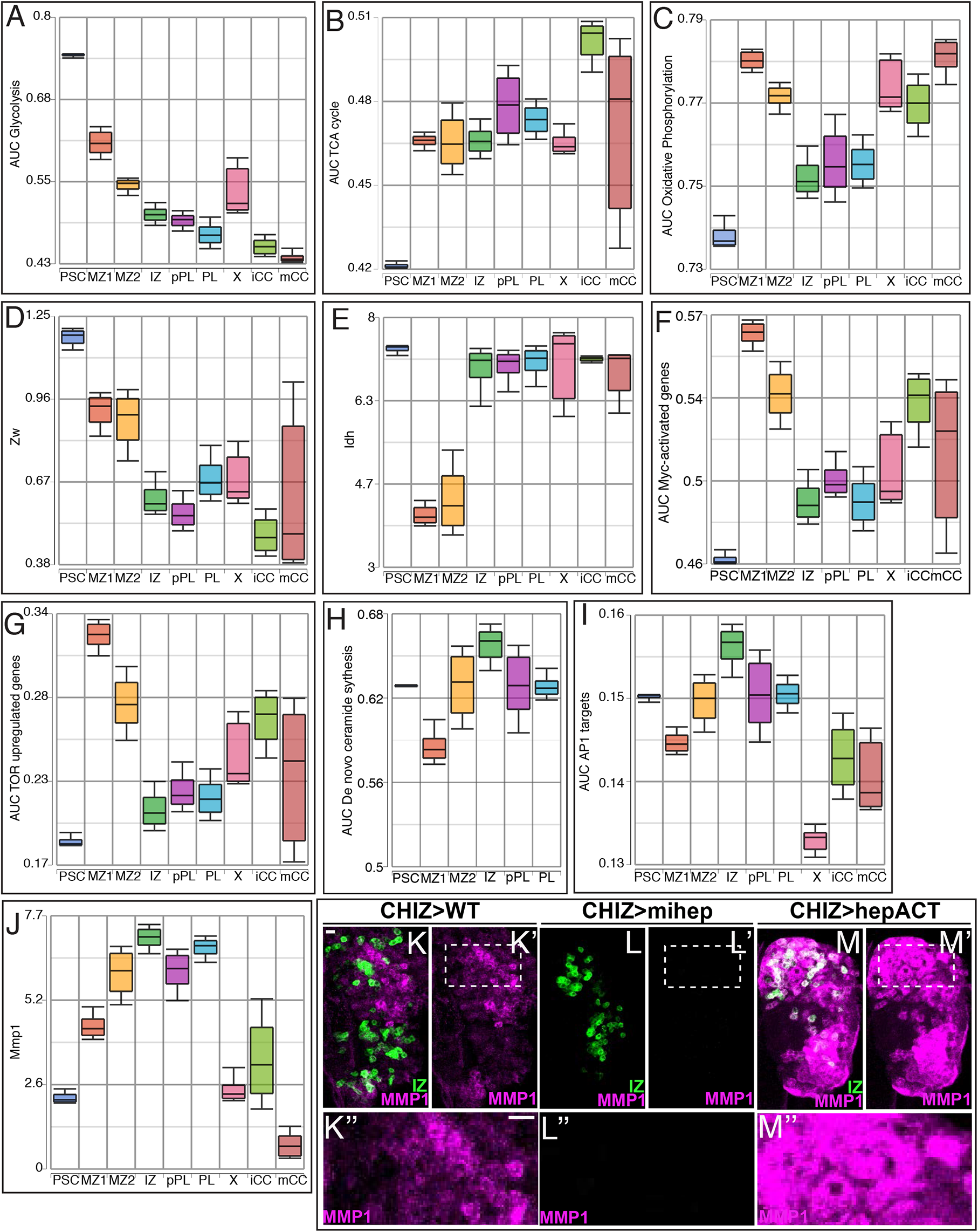
Developmental metabolism of the lymph gland by single cell analysis. Expression analysis of either single genes **(D-E, J)** or groups of genes scored as AUCell **(A-C, F-I)** are analyzed for their expression levels. **(A)** The activity of the Glycolysis pathway genes is highest in the PSC, declining progressively through MZ1, MZ2, X, IZ, proPL, PL and CCs. **(B)** TCA cycle enzymes are expressed at exceptionally low levels in the PSC compared with the cells of other clusters. **(C)** Expression of oxidative phosphorylation pathway enzymes is low in the PSC, high in MZ1 and MZ2 and moderate in IZ, proPL, and PL clusters. **(D-E)** Cytosolic NADPH producing dehydrogenases *Zw* **(D)** and *Idh* **(E)** are expressed at high levels in the PSC but dramatically lower in the MZ. *Zw* declines, while *Idh* levels are high, in IZ, proPL, PL, and CC clusters. **(F-J)** Expression trends of growth and stress related genes. **(F)** The expression of Myc targets is high in MZ and drops dramatically in IZ, proPL, and PL clusters. **(G)** TOR pathway upregulated genes follow a similar pattern. In contrast, the IZ is most prominent in the activities of *de novo* ceramide synthesis **(H),** AP1 targets **(I)** and *MMP1* expression **(J)**. **(K-M’’)** Manipulation of JNK pathway using *CHIZ-GAL4*, *UAS-mGFP* (green) that marks IZ cells (Spratford et al. 2020). Immunolocalization of MMP1 is shown in magenta. Images are maximum intensity projections of the middle third of a confocal z-stack of lymph glands from wandering 3rd instar larvae. Below the main panels are magnifications of regions boxed in K’, L’ and M’ respectively. Scale bars, 10 µm. **(K-K’’)** Moderate levels of MMP1 protein expressed in close proximity to IZ cells in wild type (WT). **(L-L’’)** A microRNA-based depletion of *JNKK/hep* in the IZ cells results in loss of MMP1 throughout the lymph gland. **(M-M’’)** Overactivation of JNK by a constitutively active isoform of JNKK/hep causes a large increase in MMP1 staining throughout the lymph gland.

As for differences, 144 of the MZ enriched genes are specifically enriched in MZ1 and 54 are specifically enriched in MZ2. More importantly, by multiple criteria, MZ1 and MZ2 show evidence of distinct biological functions. For instance, MZ1, but not MZ2, expresses some genes that are also found in the PSC, such as *dlp* and *kn/col* (Figure 2-figure supplement 1E). But MZ1 does not express the established PSC marker *Antp* (Figure 1K). Instead, high *Ubx* expression seems to be a hallmark of MZ1 (Figure 2-figure supplement 1I). Several glycolytic genes are expressed at much higher levels in MZ1 than in MZ2 suggesting distinct metabolic requirements (explored later in Figure 3). MZ1 progenitors are also specifically enriched for the metabolism related member of the steroid hormone receptor superfamily, Hnf4 (Figure 2-figure supplement 1J), which is bound by a fatty acid ligand and activates enzymes for beta-oxidation (Palanker et al. 2009). The MZ1 progenitors are also enriched for components involved in Notch signaling (Figure 2-figure supplement 1K). It has been reported that at an earlier time point in larval life, a transient activity for Notch is seen in progenitors physically adjacent to the dorsal vessel (Dey et al. 2016). At the specific stage of development in this study, MZ1 cells are the most immature of the progenitors reminiscent of the cell type described as “pre-prohemocytes” (Jung et al. 2005). Whether MZ1 is derived from the first instar transient progenitors (Dey et al. 2016; Ferguson and Martinez-Agosto 2017; Tiwari et al. 2020) is not yet clear. The cells retained in the third instar that are active for Notch has been studied in more detail by Cho et al., (2020) whose analysis extends to different developmental stages. In our direct comparison of cell size by forward scatter (FSC) in flow cytometry, a subset of MZ cells with very high *dome*^MESO^ (GFP), and virtually no *Hml* (DsRed), are especially large even when compared with other MZ cells (Figure 2-figure supplement 1L). This population of larger MZ cells might include MZ1 since large cell size is reported as a characteristic of very early hematopoietic progenitors in *Drosophila* (Dey et al. 2016).

Other distinct biological differences between MZ1 and MZ2 involve expression of specific immunity-related genes. For example, MZ2 (but not MZ1) progenitors are specifically enriched for all four Cecropin genes (*CecA1*, *CecA2*, *CecB*, *CecC*), which are involved in antibacterial humoral response downstream of the Toll and Imd pathways (Supplementary file 1). Other Imd-related genes such as *Jra* and *PGRP-SC2*, are also enriched in MZ2 compared to MZ1. This is also true for genes in the Toll pathway such as *grass*. AUCell analysis shows a similar trend for Toll and Imd pathways, both represented by higher AUC scores in MZ2 than MZ1 (Figure 2-figure supplement 1M-N). Overall, these results indicate that the slightly more mature MZ2 cells might be better poised to respond to immune challenge than MZ1 cells, while MZ1 cells are the least mature and are more likely to respond to different metabolic and local signaling cues.

#### Cluster X

X is a very small cluster of cells (∼1% of total) with a rather unique genomic composition. While most zones, which are much larger than X, are enriched for approximately 100-200 genes, ANOVA analysis suggests that for X this number is over 2000 (Supplementary file 1). We concentrate here on a limited number of defining aspects of X. It is very clear from the data that X represents a mitotic component of the lymph gland and includes cells from multiple zones. Transcripts for 206 of the 565 total cell cycle related proteins (GO:0007049, FDR=1.51E-27) are enriched in cluster X, with the five most highly enriched being *cdc25/stg* (30 fold), *cdt1*/*dup* (11 fold), *Mcm5* (10 fold), *Claspin* (9 fold), and *dap/p21* (9 fold). These are all known regulators of the cell cycle (reviewed in Lee and Orr-Weaver 2003; Berridge 2014). Of these, the first four are best known for their function in S and G2 and although p21 functions with Cyclin E in G1, its expression initiates in late G2 (de Nooij et al. 1996; Lane et al. 1996). Functional analysis of the entire group of cell-cycle related genes provides insights about cluster X with additional granularity. 64 genes are involved in the process of cell division (of 249 total mitosis-related genes, GO:0051301, FDR=2.15E-7) and 55 genes are involved in DNA replication (of 87 genes, GO:0006260, FDR=9.00E-14). Highly enriched genes include *Cdk1, Cdk2*, *aurA, aurB*, *polo*, *CycA*, *CycB*, *Chk1/grp*, spindle checkpoint proteins such as *mad2*, *Bub3*, and *BubR1*, general regulators of the cell cycle checkpoints, and DNA repair enzymes. Genes encoding 47 components of the mitotic spindle and kinetochores, including *Incenp*, *Spindly*, and members of the kinesin superfamily (*pav, Klp10A, 61F,* and *67A*) are also enriched in Cluster X (29 of 51 kinetochores, GO:0000776, FDR=5.34E-7; 23 of 45 mitotic spindle, GO:0072686, FDR=4.10E-4). AUCell analysis further suggests that Cluster X shows high levels of “mitotic G2/M transition” activity (Figure 2-figure supplement 2A). Many genes involved in DNA replication including *PCNA*, nearly all members of the MCM complex (*dpa*, *Mcm2*, *3*, *5*, *6*, and *7*), and many components of the nuclear replication fork, such as *Rfc3* and *Rfc4*, are highly enriched in this cluster. Interestingly, single cell transcriptomic study of the human HSC/HSPC cells from the bone marrow (Velten et al. 2017) also found small high cell cycle activity clusters with characteristics similar to X.

Particularly noteworthy, and perhaps surprising for this cluster, is the high representation of genes involved in intra-S DNA damage checkpoint, without any externally induced stress to cause DNA damage (Figure 2-figure supplement 2B). Several studies suggest that the regulation of replication forks in transcriptionally active cells causes replication stress and this constitutes a crucial function of the intra-S checkpoint under normal conditions (Lee et al. 2012; Blythe and Wieschaus 2015; Iyer and Rhind 2017). The expression profile related to this mechanism is striking in its specificity for cluster X, as all other clusters are virtually lacking in this unusual process (Figure 2-figure supplement 2B). The intra-S checkpoint slows down DNA synthesis and progression of replication forks by suppressing the firing of new origins of replication (Iyer and Rhind 2017). This process likely stabilizes existing replication forks and therefore “replication stress” is a means to control progression of the cell-cycle (Berti and Vindigni 2016; Zou and Nguyen 2018). In the lymph gland, the MZ cells are prolonged in their G2 state (Sharma et al. 2019) and it is attractive to speculate that replication stress related S-phase events are, at least in part, responsible for the slow-down of the subsequent G2.

Finally, 11 of the 12 members of the Myb complex (Myeloblastosis oncoprotein family), including the *Myb* transcription factor itself, are specifically and highly enriched in Cluster X (Figure 2-figure supplement 2C-D). Myb helps regulate DNA damage checkpoints and homology assisted DNA repair in breast cancer cells and also performs a similar role in the DNA repair process in *Drosophila* (Yang et al. 2019). Myb plays crucial roles in G2/M transition, transcriptional activation, and epigenetic regulation of gene expression in *Drosophila* blood cells (Davidson et al. 2005) and is known to play roles in mammalian hematopoietic cells (Greig et al. 2008). Future studies of the Myb related pathway may clarify the role of cluster X in the context of lymph gland hematopoiesis.

Cluster X is physically separated from the larger core group of clusters on the t-SNE and although small, the X cluster itself is separated into two islands (Figure 1J). One of the X islands (∼30%) is physically closer to the MZ, while the larger one (∼70%) maps in the direction of the PL cluster (Figure 1J). The smaller island is more MZ-like (X^MZ^) and the larger is more CZ-like (X^CZ^) in gene expression (Figure 2-figure supplement 2E). Subclustering of X leads to 3 separate groups, one which corresponds directly to X^MZ^ and the other two represent a split of X^CZ^ into two subclusters, X^TR^ (transitional) and X^PL^ (plasmatocyte-like) (Figure 2-figure supplement 2E-F). These subclusters show different patterns when profiled for hallmark zone-specific genes and markers such as *Tep4, collier, shg, dome, Hml, Pxn, NimC1, Col4a1, vkg*, *EGFP* or *DsRed* (Figure 2-figure supplement 2E). The clear variation in expression of MZ and CZ genes within X provides the basis for the designation X^MZ^, X^TR^, and X^PL^ for the three subclusters. Based on trajectory analysis of cluster X alone, X^MZ^ is earliest in pseudotime and X^PL^ is the latest (Figure 2-figure supplement 2G-H). X^TR^ represents a mid-point transition into the PL state (Figure 2-figure supplement 2G-H). As expected, the trajectory diagrams also reveal similar transitions for individual genes that mark the MZ and PL populations, such as *shg* and *NimC1*, respectively (Figure 2-figure supplement 2I-J). In contrast, the mitosis and DNA damage-related genes are fully dispersed over pseudotime and do not follow the subclustered distribution over X (Figure 2-figure supplement 2K-L). The X subclusters do not express any of the IZ-specific genes (Figure 2-figure supplement 3A) and its non-uniform but high level of expression of *shg*/*Ecad* (Figure 1K and Figure 2-figure supplement 2I) is particularly intriguing. When these facts are evaluated together with the physical location of X, in the region between MZ and PL, with the gene expression profile, and with the data from the developmental trajectory, it seems that cluster X likely represents a direct transitional path between MZ2 and PL with a minor presence during normal homeostatic development.

#### IZ cluster

The scRNA-Seq and bulk RNA-Seq results concur in their placement of the top IZ-specific markers such as *CG30090*, *lectin-24A, MFS3,* and *CG13482* (Figure 1E, K; Figure 2-figure supplement 3A). *CG30090* and *lectin-24A*, in particular, are useful new IZ markers since their expression is low in all other clusters (Figure 2-figure supplement 3B-E). *MFS3* and *CG13482* are also differentially expressed in IZ, but their expression is fairly high in some cells of other clusters as well (Figure 2-figure supplement 3F-G). IZ and a subset of the proPL cells share a significant proportion of the 35 genes that are enriched in the IZ cluster (Figure 2-figure supplement 3H).

Single cell analysis also confirms that the IZ cluster expresses lower levels of canonical CZ markers compared to the plasmatocyte clusters and does not express mature blood cell markers such as *NimC1*, *PPO1*, *lz*, and *hnt* (Figure 1K). On the t-SNE, the IZ cluster lies between MZ2 and PL (Figure 1J), consistent with its intermediate nature between progenitors and differentiated cells. Although IZ cells form a single cluster, they are found in multiple trajectory states 3, 4, 5, and 7 (Figure 2F), which is consistent with their transitional status. IZ cells first appear on the developmental trajectory in State 3 (IZ-3), then they progress into states 4 (IZ-4) and 5 (IZ-5), and terminate in state 7 (IZ-7) (Figure 2F, N-P, R). IZ-3 borders MZ2-3 in the t-SNE and, as the earliest IZ subzone in pseudotime, it represents the step immediately after the MZ2-3 state (Figure 2A-C, F, N).

A small number of IZ-5 cells lie near CCs on the t-SNE, while all other IZ-5 cells place between IZ-4 and IZ-7 (Figure 2-figure supplement 3I). IZ-7 lies at the border with PL and therefore represents a transition to committed plasmatocytes (Figure 2-figure supplement 3I). Together these results suggest that IZ-5 cells have the capacity to directly become crystal cells (CCs), or alternatively, via IZ-7, they take on a plasmatocyte fate. Finally, IZ-independent paths to mature blood cells are also observed that are described later.

#### CC clusters

The cells of the CC cluster are identified by the high expression of their canonical markers including *lozenge (lz)* and its target genes (Figure 1K; Figure 2-figure supplement 4A). They are also characterized by the complete absence of *NimC1* (Figure 1K). In agreement with the bulk RNA-Seq results, single cell experiments also show two distinct subpopulations, iCC and mCC (Figure 1J), which are readily distinguished by the maturation markers *PPO1* and *PPO2* (Figure 2-figure supplement 4B-C). mCC displays higher levels of *PPO1*, *PPO2*, *lz*, and *hnt* and lower levels of *Hml* when compared with iCC (Figure 2-figure supplement 4B-F). Both *lz* and *hnt* correlate positively iCCs and mCCs, while *Hml* correlates in a very strong negative manner, especially in mCCs, with *PPO2* (Figure 2-figure supplement 4G-I). Thus, based on multiple criteria, the two subclusters of CC represent an immature (iCC) and a mature (mCC) population.

Trajectory and pseudotime analysis shows that a vast majority (95%) of CCs map to the terminal state 6 (Figure 2A-C, I-J, P-R). The mCC-6 cells map at the very tip of the terminal branch on the trajectory, later in pseudotime than iCC-6 (Figure 2A-C, I-J, P-R). The remaining (5%) of CCs are seen in states 5 or 7 and belong to the immature CC class (i.e. iCC-5 and iCC-7). Although small in number, they represent distinct developmental paths in the formation of crystal cells. These paths are based on adjacencies found on the separate crystal cell island whereby the broader tip of this island bifurcates and one arm contains a small number of IZ-5s while a set of PL-7 cells occupy the second arm with no overlap between the arms (Figure 2-figure supplement 4J-K). The iCC-5 cells express higher levels of the IZ-specific gene *CG30090* than in other CCs and they map nearest to IZ-5 on the t-SNE (Figure 2-figure supplement 4K-L). This suggests that iCC-5 is derived from IZ-5. The iCC-7 cells are near PL-7s and express slightly higher levels of *NimC1* than other CCs suggesting that iCC-7 is derived from PL-7 representing a plasmatocyte to crystal cell path (Figure 2-figure supplement 4K, M). This PL to CC route is consistent with the observed expression of *Hml* reporters in the iCCs (Figure 1G; Figure 1-figure supplement 1B-B’). These transitions are also supported by the fact that both populations of PL-7 and IZ-5 cells that are detected on the crystal cell island express higher *lz* than in other PL/IZ populations (Figure 2-figure supplement 4N), even as by expression-based clustering, neither population represents a CC cell type. To summarize the conclusions from the above findings, iCC-5 and iCC-7 are derived from IZ-5 and PL-7, respectively, and these iCC subclasses transition to iCC-6. Finally, the iCC-6 cells (low *PPO1/2*) mature to the terminal mCC-6 state (high *PPO1/2*).

Transcripts for heat shock and other chaperone-like factors are enriched in CCs. These include the heat-shock transcription factor *Hsf (HSF1)* and several heat shock proteins (9 Hsps, 2 Hscs, and Hsp40/DnaJ-1). Of these, previous work has reported a role for DnaJ-1 in CC formation (Miller et al. 2017). Additionally, stress response and unfolded protein binding genes such as *foxo, mrj*, *nudC*, *Tpr2*, and *Droj2* are also enriched in CCs (Figure 2-figure supplement 4O). These results are consistent with genetic analyses that place CCs as important mediators of stress response (Sorrentino et al. 2002; Cho et al. 2018), in addition to their well-known activity in blood clotting, wound healing and immunity (Binggeli et al. 2014; Letourneau et al. 2016).

#### proPL and PL clusters

The initial identification of CZ cells was based on their expression of genes generally considered to be markers of plasmatocytes, such as *Hml*, *Pxn*, *Col4a1*, and *vkg* (Figure 1K). However, scRNA-Seq based expression profiling shows that the hallmark and newly identified CZ-enriched genes are not entirely specific to plasmatocytes. Varying but significant amounts of their transcripts are found in CCs and in the IZ (Figure 1K; Figure 2-figure supplement 5A-D). The single cell analysis reveals additional genes that are more specific to plasmatocytes and are not expressed in CCs. These include the laminins *LanA, LanB1*, and *LanB2*, as well as *Pvr*, and *NimC1* marker (Figure 2-figure supplement 5A) (Sears et al. 2003; Urbano et al. 2009; Grigorian et al. 2013). Pvr is highly enriched in the CZ although genetic analysis suggests a very low level of activity in the MZ (Mondal et al. 2011). Generally, the plasmatocyte-enriched genes are expressed at very low levels in the IZ, with *NimC1* virtually absent in these cells (Figure 2-figure supplement 5A).

Two separate clusters (proPL and PL) identified in scRNA-Seq both express the plasmatocyte-specific genes described above and neither cluster expresses CC specific genes (Figure 1K; Figure 2-figure supplement 5A). One of these clusters (pro-plasmatocyte or proPL) generally expresses lower levels of plasmatocyte markers (Figure 1K; Figure 2-figure supplement 5A) and appears earlier in pseudotime (Figure 2A-C, G). The second cluster (plasmatocytes or PL) initiates later in pseudotime (Figure 2A-C, H) and has higher levels of plasmatocyte markers than proPL (Figure 1K; Figure 2-figure supplement 5A). The distinction between PL and proPL clusters is highlighted by the differential expression of at least 750 genes that are not similarly represented in proPL.

#### proPL cluster

The proPL population arises in multiple states on the developmental trajectory. Its first appearance is in state 3 (proPL-3). Later in the trajectory, pro-plasmatocytes are seen as proPL-4, proPL-5 and proPL-7 (Figure 2A-C, G). This contrasts with the fact that virtually all cells of the PL cluster (99%) are seen exclusively in State 7 (PL-7) at the terminal arm of the trajectory indicating a path to full differentiation (Figure 2A-C, H). The proPL subclasses (proPL-3, proPL-4, proPL-5, proPL-7) differ from each other in their placement within the cluster (Figure 2N-P, R), and in the expression levels of multiple mitosis- and maturation-related genes (Figure 2-figure supplement 5E-H), likely representing different cell cycle stages as well as points of maturation. Some of the characteristic features of these subclusters include a particularly low expression level of *Hml* in proPL-3 that is higher than in MZ2-3 and lower than in other proPLs. Interestingly, proPL-3 and IZ-3 have similar levels of *Hml*, while proPL-4 shows higher levels of IZ-specific genes than other proPL subsets. Also, the proPL-4 subcluster shows a high AUC score for genes that have been identified as participating in the backward or equilibrium signal (*Pvr*, *STAT92E*, *ADGF-A*, *bip1*, *RPS8*, *Nup98-96*) (Figure 2-figure supplement 5F). Both proPL-5 and proPL-7 show high *NimC1* (Figure 2-figure supplement 5G) and are virtually identical except for their pseudotime origin. A small increase in mitotic genes is detected in proPL-7 over its proPL-5 counterpart (Figure 2-figure supplement 5H). The expression analysis that separates the subclusters does so in a manner that is functionally relevant and distinguishable by markers.

As described earlier for IZ, placement in multiple trajectory states is a characteristic of transitional cells that can represent multiple developmental paths. In this sense, the IZ and proPL cells play similar roles, both representing transitional states. There is no direct overlap or adjacency between the cells of IZ and proPL, a fact easier to discern in a 3D representation of the t-SNE (Figure 2-figure supplement 5I; Video 1). In contrast, direct adjacencies are observed between MZ2-3 and proPL cells (Figure 2S). We conclude that proPL represents a developmental route between a subset of MZ progenitors that form plasmatocytes without going through the IZ state. Instead, these cells use proPL as their transitional population. There is no evidence from the current data that suggests a transitional path between IZ and proPL. They are parallel inputs that generate differentiated cells of the CZ.

#### PL cluster

*NimC1* encodes a member of the Nimrod receptor family and as a phagocytosis-related receptor, it is a well-characterized marker of mature plasmatocytes. As expected, the highest levels of *NimC1* transcripts are seen in the PL cluster (Figure 2-figure supplement 5J), the majority of which belong to the PL-7 state in pseudotime, with a very small number (and fraction) of PL cells defined by pseudotime as PL-3 (Figure 2-figure supplement 5K). These cells form a very thin but distinctive border separating MZ2-3 and PL-7 (Figure 2-figure supplement 5K). PL-3 and some of their adjacent PL-7 cells express unexpectedly high amounts of the progenitor marker *E-Cad*/*shg* (Figure 2-figure supplement 5N). Normally, *E-Cadherin* expression is a characteristic of MZ, with its expression declining dramatically in IZ/proPL and PL clusters. High levels of *E-Cad* in a small subset of PL cells suggests that they came directly from MZ cells, not via IZ/proPL transitional populations.

*NimC1* RNA is not expressed at a significant level in the medullary zone and very low levels are seen in IZ, proPL-3, proPL-4 and PL-3 (Figure 2-figure supplement 5L). Overall its expression is highest in PL-7. Within the bounds of PL-7, however, finer analysis reveals a more complex narrative. It is possible to further subcluster PL cells, which breaks PL-7 into four smaller groups that we named PL-7a, PL-7b, PL-7c and PL-7d (Figure 2K; Figure 2-figure supplement 5K). Surprisingly, both proPL-5 and proPL-7 are significantly higher in their *NimC1* expression than in any other proPL cluster-state and they are similar, instead, to PL-7a and PL-7b (PL-7a/b for convenience; Figure 2-figure supplement 5L). By far, the highest *NimC1* levels are reserved for PL-7c and PL-7d (PL-7c/d) located along the edge of the t-SNE (Figure 2-figure supplement 5K-L). In summary, amongst the PL-7 plasmatocytes, two very distinct levels of *NimC1* transcript are detected. Pseudotime analysis shows that low (PL-7a/b) and high (PL-7c/d) *NimC1*-expressing cells arise early and late in development, respectively (Figure 2-figure supplement 5M). Based on genome-wide transcript enrichment data, we believe that all PL cells are a single cell type (plasmatocytes). However, inasmuch as differences in *NimC1* levels and temporal distinctions in pseudotime together can be used as measures of maturity, we conclude that these plasmatocytes follow a progressive maturation path from the proPL-5/7 and PL-7a/b to the more mature PL-7c/d cells. As alluded to earlier, and further discussed in the section on developmental paths, a fraction of CCs are derived from the PL (rather than proPL) population. The PL-7 cells found on the CC island primarily belong to the less mature PL-7a/b clusters rather than the more mature and differentiated PL-7c/d.

### Comparative gene enrichment

To better understand major genetic components that control similarities and differences between differentiating cells of IZ, proPL, PL, and CC clusters, we performed gene enrichment analysis to identify GO terms that characterize a cluster and then used the gene lists generated by this process to analyze their AUCell values in cells of each cluster. An important finding from this study is the enrichment of basement membrane organization components in IZ, proPL, and PL, relative to the MZ and CCs (Figure 2-figure supplement 6A). Known ECM/basement membrane protein components identified include *vkg*, *Col4a1*, *SPARC*, *Laminins* (*A*, *B1*, and *B2*), and *Tiggrin* (Pastor-Pareja 2020). Collagen secretion related genes are progressively high in IZ, proPL, and PL, with PL cells exhibiting the highest levels (Figure 2-figure supplement 6B). This pattern is seen, for example, for the hydroxyproline production related genes *PH4alphaEFB*, *Pdi*, and *Plod* that are all involved in the formation of functional collagen trimers prior to their secretion (Lamande and Bateman 1999; Gorres and Raines 2010; Rappu et al. 2019) (Figure 2-figure supplement 6C-D).

The fact that the MZ cells lack basement membrane proteins can be attributed to their non-secretory nature. Their defining feature is the tight interactions between the cells by E-Cadherin. It was surprising, however, to find high levels of collagen and other basement membrane (BM) organization components in the transitory populations, proPL and IZ. We investigated whether this process is true for the secretion of these components as well. The last step in the secretion process involves genes such as *Crag* (Calmodulin binding protein), *Sec23* and *Sec24CD* (vesicular transport important for secretion), *Pis* (Phosphatidylinositol synthase), and *sktl* (PI4P5K) (Norum et al. 2010; Pastor-Pareja and Xu 2011; Devergne et al. 2014; Isabella and Horne-Badovinac 2015). This BM secretion gene set is significantly more enriched in PL over either IZ or proPL (Figure 2-figure supplement 6E). We conclude that basement membrane proteins initiate their expression in the transitory IZ/proPL cells and even undergo maturation but the entire secretory mechanism becomes fully functional only at the PL stage.

Not all pathways are neatly confined in transcription to single zones. For example, AUCell analysis shows that components of the Ras/MAPK signal transduction pathway are expressed broadly, enriched in IZ, proPL, and PL cells, and substantially de-enriched in crystal cells (Figure 2-figure supplement 6F). Such Ras/MAPK signal related transcripts include several guanyl-nucleotide exchange factors (e.g. *CG42674*, *mbc*, *Rgl*, and *Rabex-5* among others) as well as components of MAPK signaling (such as *p38a*, *RhoL*, *Rac2*, *edl*, and *Drk*). Furthermore, *pointed* (*pnt*; discussed in some detail later), an important player in Ras/MAPK signaling, is also highly expressed in IZ, proPL, and PL but is much lower in crystal cells (Figure 4A). In stark contrast, the Notch pathway components show the opposite pattern with enrichment in CCs and with lower levels in IZ, proPL, and PL (Figure 2-figure supplement 6G).

**Figure 4.**
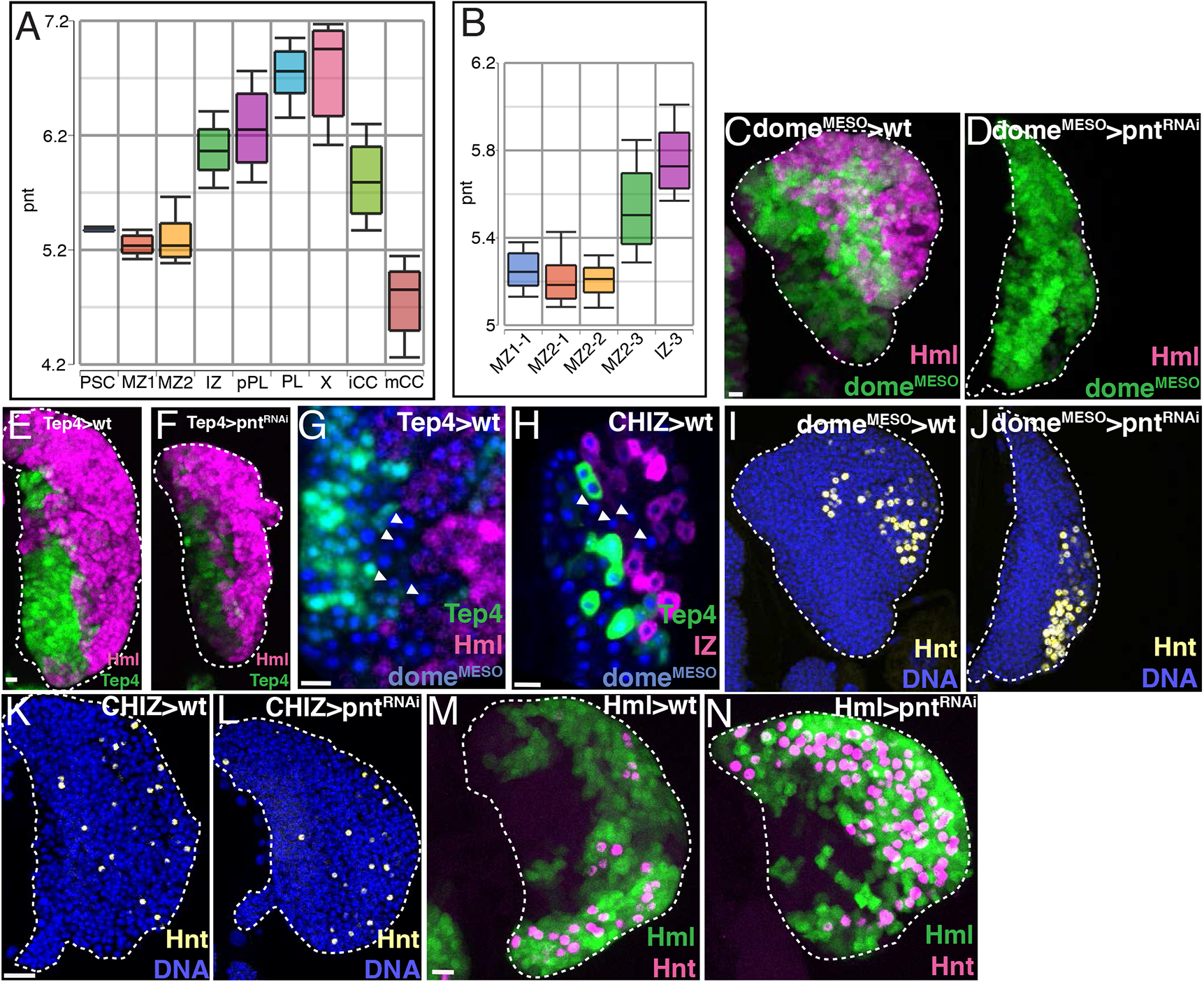
Role of Pnt in lymph gland development. **(A)** *pnt* expression is low in the PSC, MZ1, and MZ2, rises in IZ, and then further in proPL, PL, and X, decreasing significantly in iCC and mCC. **(B)** Significant *pnt* expression is first observed in the MZ2-3 subcluster and it further increases in IZ-3. **(C-D)** Genetic analysis of *dome^MESO^-GAL4, UAS-GFP, Hml^Δ^-DsRed* lymph glands.**(C)** Control lymph gland. *dome^MESO^* marks MZ (reported by GFP, green) and *Hml* marks CZ (reported by DsRed, magenta). IZ expresses both (white). **(D)** Expression of *pnt^RNAi^* in the MZ cells that are *dome^MESO^* positive (green) prevents *Hml* expression (magenta) and the formation of IZ and CZ cells. Contrast with **(C)**. **(E-F)** Genotype, *Tep4-GAL4, UAS-GFP, Hml^Δ^-DsRed*. **(E)** Control lymph gland. *Tep4* (green) is expressed in a subset of MZ precursors. *Hml* (magenta) marks IZ/CZ. **(F)** *pnt^RNAi^* expressed in *Tep4*+ MZ cells (green) has no effect on *Hml* (magenta) expression and formation of CZ cells. Contrast with **(E)**. **(G)** Genotype, *Tep4-GAL4, UAS-GFP, dome^MESO^-EBFP, Hml^Δ^-DsRed.* Late progenitors marked by arrowheads are *dome^MESO^* (blue) positive but *Tep4* (green) negative and *Hml* (red) negative. **(H)** Genotype, *CHIZ-GAL4, UAS-mGFP, dome^MESO^-EBFP, Tep4-QF2, QUAS-mCherry*. Image confirms a population (marked by arrowheads) of *dome^MESO^* (blue) positive, Tep4 (green) negative, and IZ (red) negative progenitors that are post-*Tep4* and pre-IZ. **(I-J)** Genotype is the same as in **(C**-**D)**. **(I)** Control, showing all nuclei (DNA, blue) and crystal cells (Hnt, yellow). **(J)** Depletion of *pnt* in *dome^MESO^* positive MZ cells does not prevent formation of Hnt+ crystal cells. Contrast with strong *Hml* phenotype seen in **(D)**. **(K-L)** Genotype, *CHIZ-GAL4, UAS-mGFP* (GFP not shown). **(K)** Control shows DNA (blue) and Hnt (yellow). **(L)** Depletion of *pnt* in IZ cells does not prevent formation of Hnt+ (yellow) crystal cells. **(M-N)** Genotype, *Hml-GAL4, UAS-2xEGFP*. *Hml* (green) and Hnt (magenta). **(M)** Control lymph gland. **(N)** *pnt^RNAi^* expressed in *Hml+* cells increases the number of Hnt+ crystal cells. Maximum intensity projections of the middle third of a confocal z-stack **(C-F, I-N).** Single confocal slice **(G, H)**. Lymph glands are from late third instar larvae for all images except **(G-I, M-N)** that are from early third instar. All scale bars, 10 µm.

#### PSC Cluster

Known canonical PSC markers, such as *Antp*, *col*, and *dlp* are all co-expressed at high levels in the PSC cluster (Figure 1K). Additional highly expressed genes include: *Pvf1*, *Dad*, *Dif*, and *EGFR*, several of which are known to play roles in PSC function (Mondal et al. 2011; Pennetier et al. 2012; Sinenko et al. 2012; Louradour et al. 2017). The nature of the PSC has been extensively investigated prior to this study and many of the biological pathways important for PSC maintenance and function have been identified (Krzemień et al. 2007; Mandal et al. 2007; Sinenko et al. 2009; Morin-Poulard et al. 2016; reviewed in Luo et al. 2020). When detectable, their corresponding genes are enriched in the PSC including: TGF beta signaling pathway (*Dad*, *Smox*, *Ubi-p5E*, *Ubi-p63E*), signaling by Robo receptors (*Dg*, *Ubi-p5E*, *Ubi-p63E*, *dlp*), positive regulators of Wnt gradient formation (*Flo2*, *Ssdp*, *Swim*, *dally*, *dlp*, *flw*, *hipk*, *skd*, *spen*), genes that assist in cytoneme production (*Act5C*, *AnxB11*, *Egfr*, *Flo1*, *Flo2*, *dlp*, *skd*, *sqh*) (Bischoff et al. 2013), and plasma membrane bounded cell projection morphogenesis (40 different genes).

Some of the more surprising examples of enrichment in the PSC are in genes that belong to prominent metabolic pathways. That the PSC might have a unique metabolic profile is unexpected and this prompted us to investigate specificities in gene expression across many of the key metabolic pathways in the lymph gland. This analysis revealed several interesting trends.

### Metabolic pathways and hematopoietic development

Metabolic pathways are essential for the survival and function of all cells. Contrary to the viewpoint that these pathways are limited to passive participation in “housekeeping” roles during development, data on both cancer and developmental metabolism show that selective use of such pathways can be drivers of critical developmental decisions (Pavlova and Thompson 2016; Nagaraj et al. 2017; Miyazawa and Aulehla 2018; Chi et al. 2020; Li and Simon 2020; Nakamura-Ishizu et al. 2020; Tiwari et al. 2020). When participating in such a mode, metabolic pathway genes are expected to be coordinately controlled in a cell-specific manner. Complete functional insight on developmental metabolism requires analysis of the metabolome of the kind that is outside the scope of this current investigation. What we can glean from transcriptomic data is valuable only when multiple components of a single metabolic pathway are co-regulated in a zone-restricted manner. AUCell values are particularly reflective of co-regulation and are well suited for metabolic pathway analysis.

Following such a method, we find that glycolysis related genes are expressed at much higher levels and are strongly enriched in the PSC in comparison with all other clusters (Figure 3A; Figure 3-figure supplement 1A-C). On the other hand, TCA cycle and oxidative phosphorylation-related enzymes are particularly low in their relative expression in the PSC (Figure 3B-C). At a first glance this may seem to imply that the bioenergetic requirement of the PSC is solely maintained through aerobic glycolysis, similar to that seen in cancer cells that utilize glucose uptake as a means for ATP generation (reviewed in Liberti and Locasale 2016). However, this is very unlikely to be the case since we also find that the transcript for lactate dehydrogenase (*Ldh*), the enzyme involved in the last step of pyruvate/lactate interconversion is not expressed at all (below cutoff criteria), anywhere in the lymph gland, while the rest of the glycolysis-related genes are high (Figure 3-figure supplement 1B). The striking contrast between *Ldh* and the rest of the glycolysis-related genes combined with relatively low expression of TCA and Ox-Phos genes (Figure 3A-C; Figure 3-figure supplement 1A-B) implies that the PSC cells have exceptionally low bioenergetic requirements, similar to other non-proliferative, quiet populations of cells. The combined amounts of ATP generated in the mitochondrion and in the cytoplasm is expected to be at a basal level required for cell survival. Yet, in the PSC, all glycolytic enzymes genes except *Ldh* are very high. This is likely due to the fact that the gene s*ugarbabe*, which encodes a transcription factor that regulates multiple glycolysis and gluconeogenesis-related genes, is highly expressed in the PSC (Figure 3-figure supplement 1D). Its putative target genes are also similarly enriched and they exhibit a strong positive correlation with glycolytic genes in the PSC (Figure 3-figure supplement 1E-F). But if not for energy generation, what could be the need for the high glycolysis gene expression in the PSC? The data point to the importance of biosynthetic arms of glucose metabolism. For example, we find that genes of the pentose phosphate pathway (PPP), best known for its role in nucleotide synthesis (Stincone et al. 2015), are enriched in the PSC, and within the main hematopoietic compartments, the expression of PPP and glycolytic genes declines from MZ1 (high) through MZ2 to further lower in IZ/proPL/PL (Figure 3A; Figure 3-figure supplement 1G). PPP gene expression rises but glycolytic genes further decline in the CCs (Figure 3A; Figure 3-figure supplement 1G). Further contrasting with the PSC, the TCA cycle and oxidative phosphorylation genes are relatively enriched in the MZ clusters (Figure 3B-C), suggesting a higher bioenergetic status for the progenitors than that of the PSC. However, although this increased mitochondrial function facilitates ATP generation, this also raises reactive oxygen species (ROS) levels.

The PPP component enzyme *Zw* (*G6PD*) is expressed at an especially high level in the PSC (Figure 3D). G6PD is a dehydrogenase that catalyzes the first PPP pathway reaction and produces NADPH. NADPH is produced by only a handful of enzymes within the cell but is crucial for biosynthetic reactions and in maintaining oxidative balance, keeping free radical concentrations low (Ying 2007; Fan et al. 2014; Lewis et al. 2014; Kuehne et al. 2015). Importantly, NADPH is essential for maintaining glutathione in its reduced form (GSH) that functions as a scavenger of intracellular ROS. In the cytoplasm, a prominent dehydrogenase involved in NADPH production is Isocitrate dehydrogenase (*Idh*) (Geer et al. 1979a; Geer et al. 1979b), which is also highly expressed in the PSC (Figure 3E). *G6PD* and *Idh* function as dehydrogenases that utilize very different substrates, but both generate NADPH in the course of their respective enzymatic functions. Both enzymes are highly expressed in the PSC and at a significantly lower level in the MZ (Figure 3D-E). This will lead to a different capacity to generate NADPH, suggesting important and dosage-dependent roles for this metabolite in the two cell types. The predicted high NADPH-generating capacity (low ROS) in the PSC and low NADPH (high ROS) in the MZ have interesting biological correlates in past genetic studies of the lymph gland. Under conditions of homeostasis, the PSC is known to have low ROS levels, which is maintained unless the larva is parasitized by wasp eggs. If ROS is increased, either artificially (Sinenko et al. 2012) or by wasp parasitization (Crozatier et al. 2004; Louradour et al. 2017), the PSC elicits a strong immune response that causes extensive differentiation of new lamellocytes. To prevent such a stress response from occurring in normal development will require that any or all high ROS is scavenged away from the PSC. The MZ cells, on the other hand, have been shown to contain physiologically relevant but moderately high ROS during normal, wild-type development (Owusu-Ansah and Banerjee 2009). This amount of ROS is needed for signaling purposes essential for the progenitors to differentiate. The low NADPH in the MZ predicted by the low expression of dehydrogenases would cause inefficient scavenging of ROS. Added to this, the higher mitochondrial (TCA cycle and Ox-phos) activity in these cells will raise their ROS. These could be contributing factors in keeping ROS high in the MZ, although in reality, additional control mechanisms are likely involved in the steady maintenance of ROS in the MZ as well as for keeping its level undetectable in the PSC.

Additional important enzymes involved in NADPH generation are malic enzyme (*Men*) and phosphogluconate dehydrogenase (*Pgd*) (Geer et al. 1979a; Geer et al. 1979b; reviewed in Stanton 2012). The crystal cells, by far, show the highest expression for both *Men* and *Pgd*, compared to all other cell types, and both transcripts are significantly elevated with maturation of iCCs to mCCs (Figure 3-figure supplement 2A-B). Thus, the two CC subclusters have different metabolic profiles with mature crystal cells especially enriched in genes that affect redox state. High dehydrogenase levels would keep ROS low in crystal cells, and our transcriptomic data establish that this ROS suppression is further aided by antioxidants such as *Sod1*, *Catalase* (*Cat*), *Jafrac1*, and *Trx-2*, which are also all expressed at high levels in iCC and higher still in mCC (Figure 3-figure supplement 2C-F). Mutations in at least one of these antioxidants, *Trx-2*, have been reported to cause crystal cell defects (Jin et al. 2008). Importantly, high ROS triggers JNK activity, and this would result in bursting of crystal cells and release of their contents (Bidla et al. 2007). Therefore, keeping ROS levels low is critical in avoiding premature bursting and the consequent loss of crystal cells in the absence of injury or infection.

The iCCs and mCCs are also different in additional aspects of their metabolic profiles. For instance, both peroxisomal and mitochondrial fatty acid beta oxidation, as well as fatty acid synthesis genes decrease in relative expression in mCCs compared to iCCs (Figure 3-figure supplement 2G-I). On the other hand, while lipid synthesis enzymes in general are low in CCs, genes involved in the glycerolipid remodeling process (e.g., *Bbc*, *Pld*, *Lpin*, *laza*, *Plc21C*, *GK2*, *sws*, and *CG10602*) are highly enriched in mCC (Figure 3-figure supplement 2J).

Autophagy related genes are also strongly enriched in mCCs that are fairly specific in their higher expression of *Atg1*, *Atg13*, *Atg17*, *Atg18b*, *Atg4a*, and *Atg8a* relative to iCCs (Figure 3-figure supplement 2K). Glycerolipids have been shown to regulate autophagy (reviewed in Soto-Avellaneda and Morrison 2020), and we find that autophagy genes strongly correlate with genes involved in such lipid signaling pathways in crystal cells (r=0.99; Figure 3-figure supplement 2L). Additionally, chaperone proteins *Hsc70-4*, *Hsp67Bc*, and *Hsp70Bb,* which have previously been linked to autophagy in *Drosophila* (Carra et al. 2010; Kaushik and Cuervo 2012; Uytterhoeven et al. 2015) are differentially expressed in CCs (Supplementary file 1). We also find that an AUCell analysis comprising 42 genes involved in the general process of chaperone-mediated protein folding show high differential expression in mCCS (Figure 3-figure supplement 2M) and AUCell scores for this list correlate very well with autophagy-related genes (r=0.95; Figure 3-figure supplement 2N) in CCs. Future genetic explorations will likely unravel the precise link between lipid signaling, chaperone-mediated autophagy, and the maturation of crystal cells.

Among transcription factors that control metabolism-related genes, Spargel (*srl*; PGC1-alpha), is highly expressed in MZ1 (Figure 3-figure supplement 3A), and Srl targets are enriched in both MZ1 and MZ2 progenitors relative to other clusters (Figure 3-figure supplement 3B). *Srl,* a homologue of mammalian PGC1-alpha, functions downstream of the Insulin receptor/TOR signaling pathways and mediates ribosome biogenesis, mitochondrial activity, and cell growth (Tiefenböck et al. 2010; Mukherjee and Duttaroy 2013; Mukherjee et al. 2014). Srl is a transcriptional target of Myc, which is also an important mediator of Insulin/TOR-dependent regulation of ribosome biogenesis (Teleman et al. 2008). We find that Myc, its transcriptional targets, and TOR pathway regulated genes, as well as additional ribosome biogenesis genes (including RNPs and structural proteins), are all highly enriched in MZ1 and MZ2 when compared with the other major clusters (Figure 3F-G; Figure 3-figure supplement 3C-E). By direct comparison of cell size by forward scatter (FSC) measurements in flow cytometry, we find that MZ progenitors are on average larger in size than the cells of the CZ (Figure 3-figure supplement 3F), which is consistent with the higher growth-promoting pathway activity within the MZ.

The cells of the IZ frequently express intermediate levels (between MZ and CZ) of most metabolic pathway genes we have analyzed thus far, and none are the most prominent within the IZ population. An exception to this general rule is a set of genes involved in sphingolipid metabolism, in particular, the five genes required for *de novo* ceramide synthesis. These are highly enriched in the IZ cluster (Figure 3H). The enzymes encoded by these genes convert palmitoyl-CoA and Serine to Ceramide. Specifically, the rate-limiting enzyme is serine-palmitoyl transferase (*spt1*/*lace*) that combines serine and palmitoyl-CoA to form 3-keto sphingosine. *CG10425* is predicted to encode *Drosophila* 3-keto sphinganine reductase that then makes sphinganine. Dihydroceramide synthase (*schlank*) converts it to dihydroceramide, which is then converted to ceramide by dihydroceramide desaturase (*des-1*/*ifc*). When mapped individually, these genes are expressed in subsets of cells from several zones, but for the AUCell construct representing the entire set, peak expression is observed in the IZ (Figure 3H). Like other IZ-enriched genes described here, the expression is not exclusively IZ specific, but represents a rare group of genes that are expressed highest in this zone. Small sections of proPL that are IZ-like also express significant amounts of these genes (e.g. proPL-4).

Excess accumulation of ceramide in a cell is toxic and can cause cell death. The level of ceramide in the IZ cells is kept in check by at least two different strategies. The first is a mechanism that converts ceramide to more complex forms of lipids that are not harmful to the cell. This process utilizes the glycosphingolipid pathway and several of the genes involved, *GlcT*, *Ect3*/*Beta-Gal* and *CG7997*/*alpha-Gal* are enriched in their expression in the IZ cells (Figure 3-figure supplement 4A-C). High levels of glucosylceramide synthase (*GlcT*) in the IZ, is particularly noteworthy as it is the rate-limiting enzyme of the glycosphingolipid pathway and plays a controlling role in limiting ceramide accumulation in a cell (Kohyama-Koganeya et al. 2004). The second strategy employed to reduce ceramide levels is by either converting ceramide (indirectly) or sphinganine (directly) into a phosphorylated sphingolipid. The gene encoding the corresponding kinase, *Sphk1*/*Sk1* is differentially expressed in the IZ (Figure 3-figure supplement 4D).

Importantly, ceramide production has been linked to JNK activation in *Drosophila* and in other organisms (Adachi-Yamada et al. 1999; reviewed in Ruvolo 2003; Kraut 2011) and AUCell analysis of JNK/AP-1 targets shows high activity in the IZ relative to other clusters (Figure 3I). AP-1 targets also show a high degree of correlation with *de novo* ceramide synthesis in IZ cells (r=0.94; Figure 3-figure supplement 4E). Prominent amongst these highly expressed and correlated targets is *MMP1* (Figure 3J; Figure 3-figure supplement 4F) (Uhlirova and Bohmann 2006; Stevens and Page-McCaw 2012). The enzyme *GlcT* mentioned earlier that removes ceramide from the system is also a target of the JNK pathway and its expression correlates with MMP1 (Figure 3-figure supplement 4G). This would provide an opportunity for feedback inhibition where ceramide activates the JNK pathway but its target is involved in limiting free ceramide levels to prevent uncontrolled JNK activation and cell death (Kohyama-Koganeya et al. 2004).

The possibility of a link between ceramide biosynthesis, JNK pathway, and MMP1 within the transitional IZ population is intriguing from a functional standpoint, and we therefore probed this further using molecular-genetic tools. Immunolocalization using an antibody against MMP1 reveals that the expression of the protein is limited to the region of the IZ (Figure 3K-K’’). Based on the diffuse staining, we surmise that MMP1, a secreted protein, is locally distributed at the edge of the IZ and is likely to act as a metalloprotease in reorganizing the ECM around the newly forming hemocytes. The RNA for *MMP1* is expected to be transcribed in the IZ cells since the diffuse protein expression is dramatically reduced when the JNK pathway is inhibited in the IZ cells (Figure 3L-L’’) and MMP1 protein increases in level when an activated form of JNKK (*hep^act^*) is expressed in the IZ (Figure 3M-M’’). Interestingly, activation of JNK in this manner does not cause extensive cell death suggesting the possible concurrent presence of a cell death inhibition mechanism (Uhlirova et al. 2005).

### Genetic analysis, context dependent gene function, and zonal expression

Graph-based clustering is extremely valuable in identifying genes with cell-type specific expression (such as *lz* or *NimC1*). However, decades of genetic and developmental analyses suggest that such functionally specific genes are rare, because most gene function is pleiotropic and context dependent. In the next two sections, we demonstrate that genome-wide transcriptomic data greatly aids and enhances the powerful genetic dissection strategies that are available in *Drosophila*.

### Case study 1. Pointed and plasmatocyte formation

As a first example, we investigate the ETS family transcription factor Pointed (Pnt), a known regulator of differentiation and proliferation in multiple fly tissues with implications in hemocyte development (Zettervall et al. 2004; Dragojlovic-Munther and Martinez-Agosto 2013; Shwartz et al. 2013; reviewed in Vivekanand 2018). *Pnt* transcript is expressed at very low levels in the PSC, MZ1-1, MZ2-1, and MZ2-2 (Figure 4A-B) and its level rises significantly in MZ2-3 (Figure 4B). Therefore, within the confines of the medullary zone, Pnt function is expected to be highest in this later subcluster. However, *pnt* levels are then even higher than in the MZ in the IZ, proPL, and PL populations, which suggests multiple functions in these diverse cell types (Figure 4A). *Pnt* levels decline in CCs suggesting low RTK-related activity in these cells (Figure 4A).

In search of possible Pnt function in the different zones where it is differentially expressed at high levels, we first knocked down Pnt function specifically in the MZ (using *dome^MESO^-GAL4, UAS-pnt^RNAi^*). This, we find, blocks differentiation of the progenitor population (Figure 4C-D). No *Hml* positive cells, in the form of either IZ, proPL, or PL are detected (Figure 4C-D). In stark contrast, the same *pnt^RNAi^* expressed in the high *Tep4* positive early MZ progenitors (*Tep4-GAL4, UAS-pnt^RNAi^*), has no observable phenotypic consequences (Figure 4E-F). Based on *Tep4* expression levels, the *Tep4*-based enhancer will drive high amounts of the RNAi in MZ1-1 and MZ2-1 but not in MZ2-3 (Figure 2-figure supplement 1F). Thus, removing Pnt from the early MZ cells has no phenotypic effect, and within the MZ, Pnt must function in a *dome^MESO^* positive and *Tep4* negative population of cells. Indeed, using a genetic background that allows simultaneous visualization of three different reporters, we detect by direct observation, a significant number of *dome^MESO^* positive, but *Tep4* and *Hml* double negative, cells in wild-type 3rd instar larval lymph glands (Figure 4G). These are not IZ cells as they do not express an Intermediate zone specific-Gal4 driver (*CHIZ-GAL4*, Spratford et al. 2020; Figure 4H). Taken together, these data identify a post-*Tep4* and pre-IZ population of *dome*-expressing cells within the MZ, and show that Pnt functions in this group. This group of cells includes, or is likely identical to, the MZ2-3 subcluster. The loss of function phenotype of Pnt in these specific cells suggests that Pnt is critical for entry from the late MZ into the IZ or proPL states. An interesting additional result from this experiment is that while no *Hml* positive cells develop upon Pnt loss in MZ2-3, crystal cells still form in this genetic background (Figure 4I-J). This suggests that CC formation does not require Pnt activity and that there is a direct route (perhaps made more prominent under these mutant conditions) for an MZ cell to become a CC without first going through a *Hml* positive transitional (IZ/proPL) or differentiated (PL) state.

Since the loss of Pnt in the MZ blocks entry into IZ, its even higher expression in the IZ suggests a different and additional role in this zone. This IZ function is explored in some detail in a companion paper, where we demonstrate that loss of Pnt in the IZ prevents these cells from exiting their transitional state (Spratford et al. 2020). Together with the data presented here, we conclude that Pnt is required for both entry into and exit from the IZ. In a manner similar to that seen for MZ (Figure 4I-J), crystal cell formation is not affected upon loss of Pnt in IZ (Figure 4K-L), consistent with the possibility of a direct path between MZ and CC without an intervening *Hml-*positive state.

Finally, since we find that the proPL and PL clusters express the highest *pnt* levels of all cell types (Figure 4A), we eliminated *pnt* function in *Hml* expressing cells (*Hml-GAL4, UAS-pnt^RNAi^*). This causes a large number of plasmatocytes to be converted into crystal cells (Figure 4M-N). Thus, loss of *pnt* prevents a *Hml*+ precursor from becoming a plasmatocyte, forcing it instead into a CC fate. Keeping in mind that Pnt is activated by RTK/MAPK pathways and that the Serrate/Notch pathway is important for crystal cell formation, we surmise that Notch activation is the default pathway for early *Hml*-expressing cells to become CCs, and that the activation of Pnt acts antagonistically to prevent this process thus favoring instead, the plasmatocyte fate. Similar antagonism between Notch and RTK pathways have been seen in multiple developing tissues (reviewed in Sundaram 2005).

### Case study 2. Numb/Musashi assisted non-canonical Notch signaling in crystal cells

A canonical, Serrate-dependent Notch signal is required for crystal cell formation from a *Hml*+ precursor, while a separate, non-canonical, ligand-independent and Sima (Hif)-dependent Notch signal is important for CC maintenance (Mukherjee et al. 2011). These pioneering findings, particularly the non-canonical signaling aspect, were intriguing at the time, and only by combining transcriptomic and further genetic analysis, are we now able to provide a more complete mechanistic description of this complex signaling network.

Based on the RNA-Seq data, well established Notch target genes (Krejčí et al. 2009; Terriente-Felix et al. 2013) can be classified into two groups, which for simplicity, we call type I and type II. Type I targets (such as *E(spl)m3-HLH*, *cv-c*, and *CG3847*) are expressed at a higher level in iCCs than in mCCs (Figure 5A), while type II targets (such as *CG32369*, *bnl*, and *IP3K2*), are more highly expressed in mCCs than in iCCs (Figure 5C). Using *PPO2* as a maturity marker, we find that type I targets correlate positively in iCCs, but negatively in mCCs with *PPO2* expression (Figure 5B). In contrast, type II targets positively correlate in both iCCs and mCCs with *PPO2* (Figure 5D). An example of a type II target is *bnl*, which shows a very steep rise in expression in mCC. This is reflected by staining for a *bnl* reporter in the tissue that decorates only a subset of the crystal cells, presumably the ones that are more mature (Figure 5-figure supplement 1A-A’; Tattikota et al. 2020). Genes including *bnl* and *CG32369*, that we classify as type II Notch targets, have been previously established as both Notch and Sima/hypoxia-responsive in independent studies (Li et al. 2013; Terriente-Felix et al. 2013; Kamps-Hughes et al. 2015; Du et al. 2017). Indeed, we find that enhancer sequences for these two genes contain combinations of both Su(H) and Sima binding sites (Figure 5-figure supplement 1B-C).

**Figure 5.**
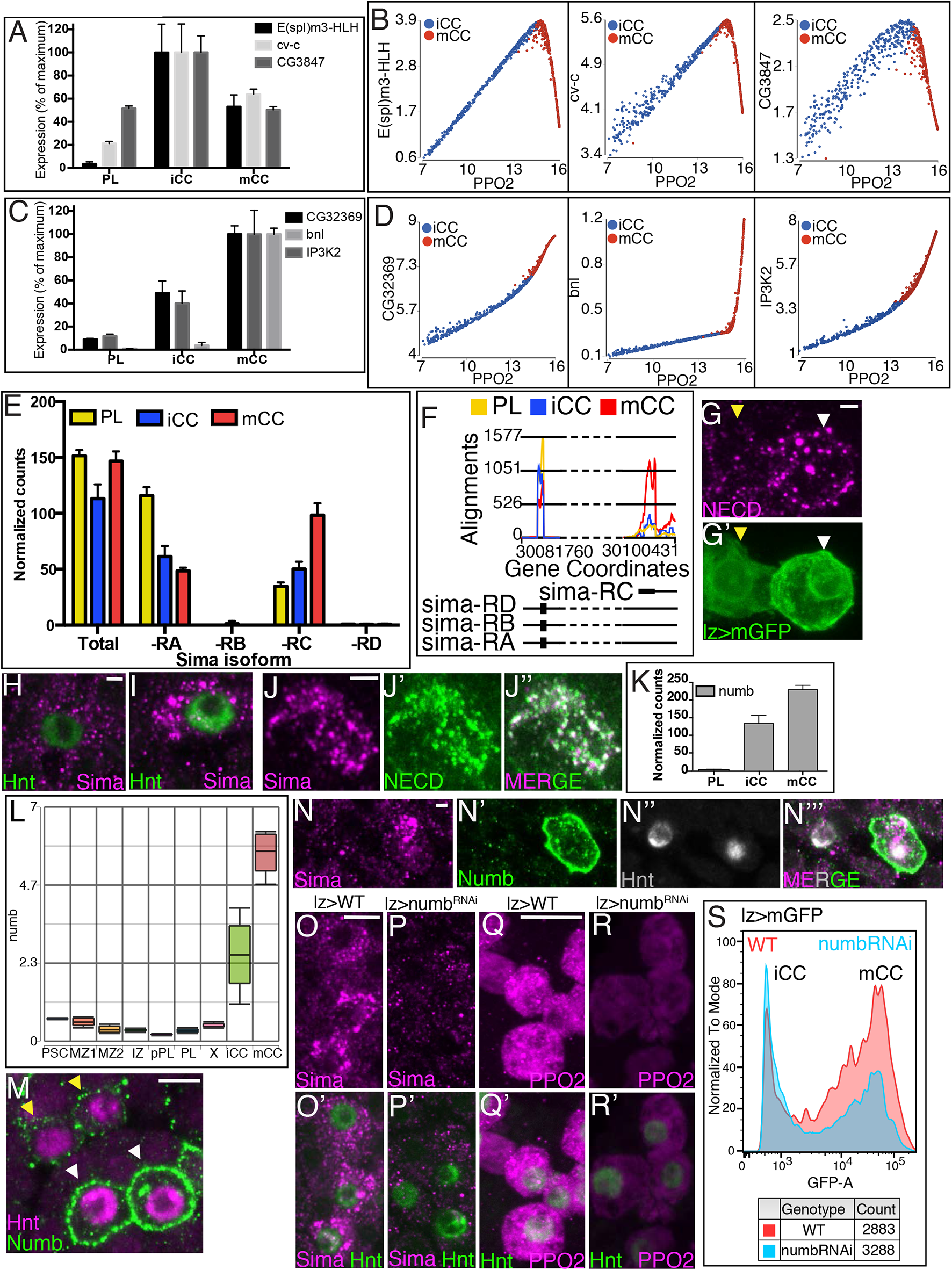
Numb promotes non-canonical Notch/Sima signaling. Data in **(A, C, E-F, K)** are based on bulk RNA-Seq. Data in **(B, D, L)** are based on single cell RNA-Seq. **(A)** Examples of “Type I” Notch targets (*E(spl)m3-HLH*, *cv-c*, and *CG3847*) with highest expression in iCC and lower in mCC, usually lower still in PL. **(B)** Type I Notch targets correlate positively with the CC maturity marker *PPO2* in iCC and negatively in mCC. **(C)** Examples of “Type II” Notch targets (*CG32369*, *bnl*, and *IP3K2*) that have their lowest expression in PLs, increase in iCCs, and are expressed even higher still in mCCs. **(D)** Type II Notch targets correlate positively with *PPO2* in iCCs and positively correlate with an even higher slope in mCCs. **(E)** Total *sima* transcript levels are similar in PL, iCC and mCC. The usually major splice variant, *sima-RA* decreases with CC maturity. The normally minor *sima-RC* isoform increases with maturity from PL to iCC to higher still in mCC. **(F)** Alignment counts specific to *sima-RA*/*RB*/*RD* are highest in PL (yellow). The alternate start exon specific to the *sima-RC* is highest in its expression in mCC (red). The exact flyBase coordinates are in **Figure 5—figure supplement 1D**. **(G-G’)** Live internalization assay in *lz-GAL4, UAS-mGFP* lymph glands with an antibody against the extracellular domain (N^ECD^) of Notch (magenta) to visualize uptake and stabilization of full-length Notch protein. Large Notch punctae are specifically located in mCC (GFP^HI^; white arrowhead) but not in iCC (GFP^LO^; yellow arrowhead). **(H-I)** Hnt (green) and Sima protein staining (magenta). Numerous large Sima punctae are detected in mCC **(I**, high Hnt) but not in iCC **(H,** low Hnt). **(J-J’’)** Full-length endocytosed Notch protein is visualized in a live internalization assay with an antibody against N^ECD^ (green) and then fixed and stained for Sima protein (magenta). Numerous large N^ECD^ and Sima punctae colocalize and therefore appear white in the merged image **(J’’)**. **(K)** *numb* transcript levels are minimal in PL, increase in iCC and are further increased in mCC. These data are based on bulk RNA-Seq analysis. **(L)** Compared to all cells identified by scRNA-Seq, *numb* transcript levels specifically increase in iCC and are even higher in mCC. **(M)** Strong Numb protein staining (green) is restricted to mCCs (white arrowheads), with stronger Hnt staining (magenta) and not detected in low Hnt-expressing iCCs (yellow arrowheads). **(N-N’’’)** Large Sima punctae (magenta) are only seen in Hnt (grey) positive crystal cells with high Numb staining (green). **(O-P’)** Genotype, *lz-GAL4.* Sima protein (magenta). Hnt protein (green) in **(O’, P’)**. **(O-O’)** Wild-type lymph glands display large Sima punctae (magenta) in Hnt+ (green) crystal cells. **(P-P’)** The large Sima punctae are eliminated when *numb* is depleted in crystal cells using *lz-GAL4 UAS-numb^RNAi^*. **(Q-R’)** Genotype, *lz-GAL4.* PPO2 protein (magenta). Hnt protein (green) in **(Q’, R’)**. **(Q-Q’)** PPO2 is high in most Hnt+ crystal cells in wild-type lymph glands. **(R-R’)** PPO2 levels decrease in Hnt+ crystal cells when *numb* is depleted using *lz-GAL4 UAS-numb^RNAi^*. **(S)** Flow cytometric analysis of GFP levels (GFP-Area) shows that when *numb* is knocked down (Genotype: *lz-GAL4 UAS-numb^RNAi^*), a large proportion of the mCCs (GFP^HI^) are lost while the total number of crystal cells (Count) does not change significantly (2883 in WT *vs* 3288 in *numb^RNAi^*). **(G, J, N, O-P)** Scale bars, 2 µm. **(M, Q-R)** Scale bars, 5 µm.

Despite past work showing Sima is required in CC maintenance, we detect no significant differences in *sima* transcript levels between PL, iCC, and mCC (Figure 5E). This is not entirely surprising because Sima is known to be primarily controlled at the protein level, but since our bulk RNA-Seq experiments have sufficient depth of sequencing, we are able to individually analyze the expression of the putatively proposed RNA isoforms, *sima-RA, -RB, -RC and -RD* (Figure 5E-F) for their expression. The isoforms *-RB* and *-RD* are not expressed in the lymph gland, and the most widely studied, full-length *sima-RA* isoform is higher in its expression in PLs than in CCs (Figure 5E-F). Surprisingly, and most importantly for this study, the smaller isoform *sima-RC* is expressed at its highest level in mCCs (Figure 5E-F; Figure 5-figure supplement 1D). Using isoform-specific primer sets for qRT-PCR, we confirmed that sequences corresponding to the unique 5’ regions of *sima-RC* are highly represented in mCCs (Figure 5-figure supplement 1E-F). Previous studies suggest an auto-regulation of *sima-RC* by a stabilized Sima protein (Kamps-Hughes et al. 2015). In the lymph gland, this would result in a mCC specific expression of *sima-RC*, akin to a type II Notch target (Figure 5-figure supplement 1G). Notably, the predicted protein encoded by *sima-RC* (Sima-PC) lacks the N-terminal motif for binding of the Sima partner (Tango; Hif-1Beta/ARNT), which is important for the hypoxic response elicited by full length Sima-PA (Gorr et al. 2004; Romero et al. 2008) (Figure 5-figure supplement 1H-I). However, Sima-PC retains the oxygen dependent degradation (ODD) domain that is responsible for hypoxia-related protein stabilization (Figure 5-figure supplement 1H-I; Gorr et al. 2004; Romero et al. 2008). The revelation that a very specific isoform of Sima is an important determinant of mCCs explains previous work from our laboratory showing that crystal cell maturation is independent of Tango, and does not elicit a characteristic Sima/Tango related hypoxia response and yet upon hypoxia, Sima protein levels and CC numbers increase (Mukherjee et al. 2011). These and past results suggest a role for Notch/Sima signaling specifically in the mCC population. We find that mCCs, but not iCCs, contain large punctae of endocytosed and stabilized full-length Notch receptor (Figure 5G) and large punctae of Sima protein (Figure 5H-I). Moreover, these large Notch and Sima punctae strongly co-localize in these cells (Figure 5J-J”).

### Numb and Musashi in CC determination

The punctae containing stabilized N/Sima-PC proteins suggest that endocytic mechanisms are important for this non-canonical Notch pathway. This motivated us to focus on the gene *numb*, which encodes a component of the endocytic pathway that promotes internalization and regulates trafficking of the Notch receptor thereby blocking canonical Notch signaling (Couturier et al. 2013; Yap and Winckler 2015; Johnson et al. 2016; Shao et al. 2016). Given that CC induction requires ligand-dependent canonical Notch signaling (Lebestky et al. 2000), and that Numb blocks such a signal (Frise et al. 1996; Spana and Doe 1996), we found it surprising that *numb* RNA levels are by far the highest in both iCCs and mCCs compared to all other lymph gland cell types (Figure 5L). *numb* mRNA also increases significantly during the transition from iCC to mCC (Figure 5K-L), correlating strongly (r = 0.99) with *PPO2* (Figure 5-figure supplement 2A). The steep increase in *numb* mRNA is indicative of a CC-specific transcriptional regulation, likely controlled by Notch as seen in other developmental systems (Rebeiz et al. 2011). This is consistent with our findings that a constitutively active form of Notch (Notch^ACT^) expressed in CCs raises Numb levels while *Notch^RNAi^* has the opposite effect (Figure 5-figure supplement 2B-D’’’).

By far, the most spectacular control of Numb is at a post-transcriptional level. In spite of the large expression of *numb* RNA in iCCs, Numb protein expression is exclusively detected in mCCs and not in iCCs (Figure 5M; also Figure 6A-C’). In fact, in our hands, immunohistochemically detected Numb protein is the best marker for the mCC population. We also find that large Sima punctae are exclusively seen in the Numb-expressing mCCs (Figure 5N-N’’’). These full-length Notch/Sima punctae co-localize with Numb punctae and are seen in Hrs8-2 positive early endosomes (Figure 5-figure supplement 3A-G).

**Figure 6:**
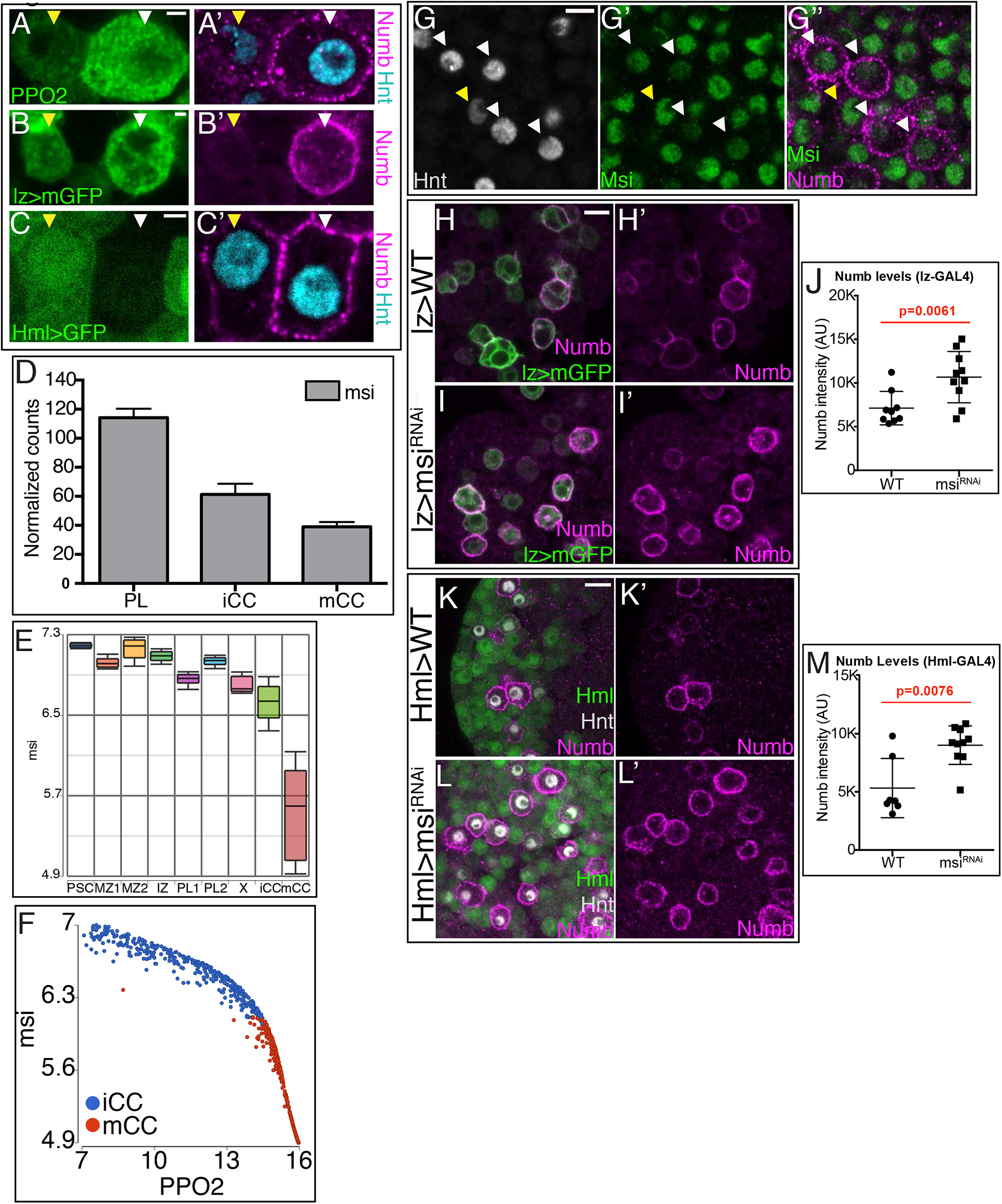
Musashi (Msi) regulates Numb protein translation. **(A-C’)** Distinction between iCC and mCC by three different criteria. A pair of cells is shown in each panel, and for each, the one on the left (yellow arrowhead) is iCC and the one on the right (white arrowhead) is mCC. **(A-A’)** iCC is low Hnt and low PPO2, while mCC is high Hnt and high PPO2. Hnt (cyan), Numb (magenta), PPO2 (green). **(B-B’)** iCC is low *lz>mGFP*, while mCC is high *lz>mGFP*. Numb (magenta), *lz>mGFP* (green). **(C-C’)** iCC is high *Hml>GFP* and low Hnt, while mCC is low *Hml>GFP* and high Hnt. Hnt (cyan), Numb (magenta), *Hml>GFP* (green). **(A’, B’, C’)** Numb staining (magenta) in each panel demonstrates that by all three criteria, Numb protein is specifically seen in mCCs (white arrowhead) and not iCCs (yellow arrowhead). **(D)** *msi* transcript levels decrease with CC maturity from PL to iCC to mCC. Data from bulk RNA-Seq. **(E)** single cell RNA-Seq shows reasonably uniform *msi* transcript level in all lymph gland cells with the very clear exception of mCCs in which *msi* transcript is dramatically suppressed. **(F)** *msi* expression negatively correlates with *PPO2* levels in iCC and mCC populations. **(G-G”)** CCs (Hnt+; grey) with low levels of Msi-GFP (green; white arrowheads) have high Numb staining (magenta). iCCs with low Hnt staining (yellow arrowheads) show higher levels of Msi-GFP and no Numb protein. **(H-J)** Genotype, *lz-GAL4, UAS-mGFP.* Numb (magenta) is expressed at moderate levels in wild-type *lz>mGFP* CCs (**H-H’)** and increases significantly upon loss of *msi* with *lz-GAL4* driving *UAS-msi^RNAi^* **(I-I’)**. Quantitation is shown in **(J)**. **(K-M)** CCs are visualized with Hnt (grey) and *Hml-GAL4, UAS-2xEGFP* is used as a driver. In wild-type **(K-K’)** *Hml>GFP* positive (green) cells do not show high levels of Numb protein staining (magenta). Whereas upon expression of *UAS-msi^RNAi^* **(L)**, Numb protein greatly increases. Data are quantitated in **(M)**. All images are single confocal slices. See **Figure 6—figure supplement 1** for lower magnification views of lymph glands shown in **(G-L)**. **(A-C)** Scale bars, 2 µm. **(G)** Scale bar, 5 µm. **(H-I, K-L)** Scale bars, 10 µm. (p values in **(J)** and **(M)** are from t-test)

This combination of data is consistent with earlier findings that trapping full-length Notch in early endosomes promotes ligand independent Notch function (Vaccari et al. 2008; reviewed in Fortini and Bilder 2009). In the mCCs, we conclude that the exclusive and specific expression of the Numb protein allows such a trapping of the full length-Notch/Sima complex in early endosomes, enhancing rather than inhibiting signaling by this unusual and non-canonical pathway. As a direct genetic test of this model, we downregulate *numb* RNA in crystal cells (*lz-GAL4; UAS-numb^RNAi^)* and find a clear reduction in the large Sima punctae, in PPO2 expression, and in the number of mCCs, without affecting the total number of crystal cells (Figure 5O-R’; Figure 5-figure supplement 3H-N). In summary, loss of Numb has no effect on initial crystal cell specification and iCC induction; rather depletion of Numb largely prevents the maturation of iCCs to the mCC state. These results are further confirmed by flow cytometric analysis, which shows that *numb^RNAi^* expressed in all crystal cells causes a decrease in the proportion of mCCs with a concomitant increase in the iCC population (Figure 5S). Flow cytometry also allows us to estimate effects on cellular size and complexity attributed to loss of *numb*. We find that *numb^RNAi^* expressed in CCs causes smaller size (FSC) and lower complexity (SSC) in mCCs but not in iCCs when compared to wild type (Figure 5-figure supplement 4A-D). In Figure 1, we showed evidence that a subset of the most mature crystal cell population exhibits >4N DNA content attributed to endocycling (Figure 1H). We find that *numb* knockdown in CCs also results in a very clear reduction in the number of cells with >4N DNA content (Figure 5-figure supplement 4E). Loss of *numb* dramatically reduces mCCs (Figure 5S), and the ones that remain, likely due to incomplete knockdown by RNAi, show loss of endoreplication, a process that is a sign of full CC maturity.

The above results lead us to investigate how the expression of the Numb protein is restricted to mCC even as its transcript is similarly abundant in iCCs. This regulation is crucial for allowing canonical Notch signaling to be active in the early population. Previous work in mammalian systems has suggested that the RNA-binding protein Musashi (Msi) is a translational repressor of *numb* that functions by binding to *numb* mRNA (Imai et al. 2001). In the lymph gland, *msi* mRNA is expressed widely in most cells, with the clear exception of the crystal cells where its transcript is seen at a decidedly low level (Figure 6D-E), and this is especially true for the mCCs. Furthermore, *msi* RNA shows a very strong negative correlation with *PPO2* in iCCs and mCCs (Figure 6F). Finally, similar to its RNA, a Msi fusion protein, is also expressed broadly except in the high Numb positive mCCs (Figure 6G-G’’; Figure 6-figure supplement 1A-B’). A knockdown of *msi* in all crystal cells (*lz-GAL4, UAS-msi^RNAi^*) causes a strong increase in Numb protein level in CCs relative to wild type (Figure 6H-J; Figure 6-figure supplement 1C-D’’). Similarly, when *msi*^RNAi^ is driven in *Hml* expressing cells (*Hml-GAL4, UAS-msi^RNAi^*) that include the iCCs but not the mCCs (Figure 6C-C’), an even more dramatic increase in the Numb protein is readily evident (Figure 6K-M; Figure 6-figure supplement 1E-F’). Combining the established nature of Msi as an RNA-binding translational repressor with the transcriptomic and genetic data presented here, we conclude that Msi controls *numb* translation and greatly attenuates its protein level in iCCs.

In summary, during CC development, Notch signaling turns on *numb* transcription in both iCC and mCC, but the presence of Msi represses *numb* translation in iCCs. Low Numb protein levels allow canonical ligand-dependent Notch signaling to occur in iCCs. In contrast, *msi* RNA and protein is virtually absent from mCCs, although the mechanism by which *msi* transcription is down-regulated is not yet fully clear. However, the transcriptional downregulation of *msi* is critical to CC maintenance since this mechanism facilitates the translation of *numb* RNA specifically in mCCs, where it functions to promote non-canonical Notch/Sima signaling and CC maturation, while repressing canonical ligand-dependent Notch-related signals.

## Discussion

Genome-wide transcriptomic analysis of the lymph gland highlights an unexpected level of heterogeneity, in spite of the fact that all analyzed cells belong to a single, small, hematopoietic tissue. Earlier genetic studies that identified zone specific “hallmark” genes define far fewer clusters than is revealed by single cell transcriptomic analysis. In all, using conservative criteria for cluster and subcluster separation, we find nine broadly defined groups of cells: One PSC, one mitosis and replication stress related (X), two CCs (iCC and mCC), two MZs (MZ1 and MZ2), two transitional populations (proPL and IZ), and one plasmatocyte (PL). The majority of clusters fit compactly together with closely apposed boundaries without large areas devoid of cells that map in between. This implies smooth developmental transitions between the clusters and subclusters even as, from a gene-enrichment point of view, they represent different cell types. The PSC, cluster X, and CCs are different enough from the rest to cluster separately from the main (core) body of cells. However, for the remaining populations, the transition from a pre-progenitor through progenitors, IPs/proPLs, and plasmatocytes is a continuous process. This indicates a gradual rather than a stepwise binary transition in gene expression between related groups of cells. Heterogeneous but related cell-populations making continuous rather than quantal decisions holds true as well, for mammalian hematopoiesis (Velten et al. 2017; Rodriguez-Fraticelli et al. 2018).

The developmental trajectory through these clusters is not a single linear path from the earliest to the latest cell in pseudotime. Rather, it is split, with multiple branch points and overlapping cluster-identity suggesting alternate paths of development. Based on this predicted trajectory, the cells within a cluster can be further distinguished for their maturity and developmental status and the 9 clusters defined above are subclassified into 22 zonal-states. The rather large number of the cluster-states arise due to multiple developmental paths rather than because of a lack of smooth transitions between excessively dissimilar cells.

In general, the primary distinction between the branches of developmental paths can be attributed to the presence of the major transitional populations, the IZ and the proPL cells, within the lymph gland. In a less defined way, cluster X is also transitional from MZ-like to CZ-like cells. In the model below (Figure 7), the primary distinction between the two major developmental paths to plasmatocyte formation is that one involves IZ and the other proPL. More speculative, and minor, paths to plasmatocytes might involve neither IZ nor proPL. There is no developmental path detected that includes both IZ and proPL. In a manner similar, but not identical to that seen for plasmatocytes, two independent, major paths are charted for the formation of CCs.

**Figure 7:**
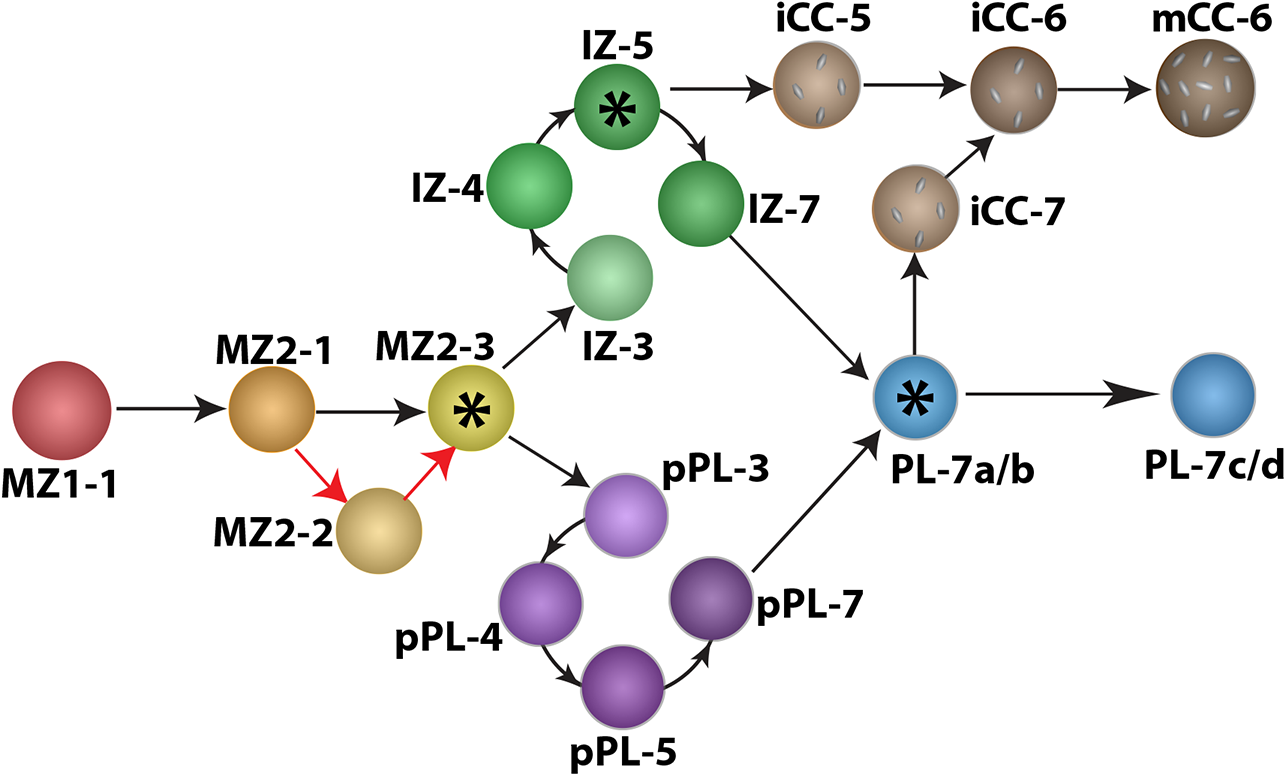
Model of developmental progression of lymph gland cells. A model for complexity and developmental progression of cell types within the *Drosophila* lymph gland (see Discussion section for details). Nodes at which cells make alternate fate choices (FC) are marked by asterisks. The PSC has a developmental origin separate from the lymph gland but shares enough similarities with MZ1 to suggest that some of the signaling functions of the PSC could be shared with MZ1. The most mature subcluster within the progenitors is MZ2-3, which is an FC point that leads to two alternate transitional zones IZ and proPL, each replete with its own set of subclusters. IZ-5 is a FC node that allows a split towards either a PL or a CC fate. The third FC node is at PL-7, a state where a choice to a CC fate is weighed against maturation to a terminally differentiated plasmatocyte. There is evidence for additional minor paths and inputs into the circuit discussed in the text. The most important feature of the model is the flexibility afforded by the identified transitional states and alternate routes towards the same or different fates. Also, this model represents a snapshot in developmental time and is likely to have evolved from a simpler earlier pattern.

IZ and proPL share common characteristics in expressing intermediate levels of CZ genes and these clusters overlap extensively on all three arms of the trajectory suggesting that their cells follow similar developmental dynamics. These two clusters are also both enriched in their expression of a significant number of shared gene products. Importantly, however, major functional differences between these two zones support the clustering and trajectory data to establish that they represent distinct cell types. Perhaps the most striking functional difference is in their signaling capacity. The equilibrium or backward signal from the CZ to the MZ that is important for progenitor maintenance initiates *via* Pvr and several unique downstream components specifically in proPLs and not in IZ cells (Figure 2). As a counter example, JNK signaling and MMP1 expression is a specific property of IZ cells (Figure 3), not seen for proPL. Cells of the two transitory states to maturity progress independently and in parallel to each other. Their overlap on the trajectory is a reflection of their independent but concurrent roles in development. This strategy allows for additional non-overlapping functions for IZ and proPL in addition to their transitory role. The equilibrium signal from proPL is important for progenitor maintenance while the IZ dependent JNK signal simultaneously promotes differentiation. Intermediate cell populations between marked progenitors have been described in T cell development in the thymus, but their signaling subtypes and paths of transition are not fully understood (Kaech and Cui 2012). We hope that in the future, *Drosophila* hematopoiesis might shed light on the functional relevance of transitional populations in mammalian hematopoiesis.

One of the distinguishing features of the analysis presented here is in its versatile approach of combining FACS sorted bulk RNA-Seq data with 10X genomics based single cell RNA-Seq results. These genomic data are then incorporated into representative examples of genetic manipulations, allowing for a detailed mechanistic interpretation of key developmental events. An independent and pioneering transcriptomic analysis of the lymph gland has been recently reported (Cho et al. 2020), which elaborates on several conclusions that are in common with ours and many others that differ. In general, the broader clusters map in a consistent manner between the two studies, but differences become apparent in the subclusters, which may partly be due to the higher sensitivity, precision and unbiased gene expression quantification afforded by the 10X platform over the DropSeq method (Macosko et al. 2015; Zhang et al. 2019). More importantly, however, our study uses unsupervised clustering without manual curation and regrouping based on known gene expression. This, combined with our use of marker-based sorting in bulk-seq to define new zone-specific genes, allows for finer discrimination between related clusters, the identification of novel cell types, and a more thorough dissection of hematopoiesis at this key developmental stage. On the other hand, the Cho et al. (2000) study has added value for its broader analysis of the lymph gland at different stages of development. The more extensive and multifaceted scheme utilized here is essential for sharper cell-type analysis and for obtaining mechanistic insights into paths followed and pathways employed during the complex hematopoietic process. None of the conclusions regarding the two transitional zones and the pathway analyses highlighted in the case studies would be possible without using a combination of the described methods. For example, the use of fluorescent markers to sort the IZ population allows us to definitively identify the distinct intermediate progenitor cell type and distinguish it from other transitional populations such as the proPL. If we attempt the subclustering process without a marker, based solely on intermediate levels of expression, different transitional cell types get combined into a single cluster. Finally, three other transcriptomic studies (Cattenoz et al. 2020; Fu et al. 2020; Tattikota et al. 2020) have examined circulating cells that have an entirely different developmental history than the cells of the lymph gland (Gold and Brückner 2015). We hope the lymph gland nevertheless provides a basis for comparison with different classes of fully mature cells. As for the lymph gland studies, for convenience, we provide supplementary material (Supplementary file 2) in which the identified gene sets for cell types defined by Cho et al. (2020) are superimposed on the clusters identified in this study. This exercise is entirely for the purpose of facilitating a direct comparison between the two sets of data for the purpose of generating a common framework and consistent nomenclature for the cluster-based analysis.

### Paths to plasmatocytes

In Figure 7 we present a model of lymph gland development from the earliest progenitor state to plasmatocytes and crystal cells based on adjacencies of expression-based clusters and their occupancy state in pseudotime. All paths initiate from MZ1, corresponding to a pre-progenitor population that expresses high levels of *Tep4* and *dome*. As development progresses through the next progenitor population, MZ2, they down-regulate *Tep4* but continue to express *dome*. After traversing through the large MZ2-1 population, the developmental status reaches MZ2-2/3, which is best described as “pre-transitional”. As development proceeds beyond MZ, the cells initiate *Hml* expression and the path to PL splits into the two separate branches. The first path to plasmatocytes is via IZ and the second is via the proPL transitional state. These split paths involve several subclusters of IZ and proPL, either of which can become a low *NimC1* version of PL (PL7-a/b) that matures to the high *NimC1* expressing differentiated plasmatocytes (PL7-c/d). While the data provide evidence for multiple developmental paths linking progenitors to plasmatocytes and crystal cells but none that we find includes an IZ-proPL segment (Figure 7).

Paths designated as “minor” involve very few cells and although consistent with our data, such paths are not yet firmly established. These are described here with the expectation that future genetic evidence will verify them. A PL cluster named PL-3 and the island cluster X seem to be involved in direct MZ to PL formation. As the name implies, PL-3s are the earliest plasmatocyte cell type to appear in state 3 of pseudotime. On the t-SNE, PL-3 is sandwiched between MZ2 and PL-7 with no intervening proPL or IZ cells. This suggests that the path could represent a direct MZ to PL transition. A similar, additional, such minor path likely includes Cluster X as an intermediary step as X subclusters into an MZ-like, a transitional, and a CZ-like subcluster suggesting a developmental transition step that does not involve either IZ or proPL. The expression pattern of *shg* (encoding E-Cadherin, E-Cad), which is especially high in the cells belonging to these minor paths, including PL-3 and X (Figures 1 and 2), provides supporting evidence, beyond the spatiotemporal arguments, for the existence of minor paths. The adjacencies of high E-Cad cells across unexpected cell types is compelling given the known biological function of E-Cad in patterning and its distinctive role in mediating morphological changes in many tissue types.

### Paths to crystal cells

Two paths, different from those described for plasmatocytes, track the formation of crystal cells and based on the number of cells involved, these two are likely to be of comparative strength. As mentioned earlier, all paths, whether they lead to CC or PL, initiate with MZ1 and MZ2, and then the first CC path follows a direct route from IZ to iCC. The other follows a transitional population to PL7a/b (*Hml* positive, low *NimC1*) and thence to iCC. A large majority of CCs are in the form of iCC-6 and they then mature to mCC-6. The events that mediate the transition from either IZ or PL to iCC were described in detail in the Results section (Figure 2).

In summary, the two major paths to plasmatocyte formation and the two that give rise to CCs, both involve one or the other of the transitory populations IZ/proPL. Minor paths that circumvent the transitional states might exist, and could in fact take on a more prominent role under mutant and stress conditions. Additionally, other minor paths, such as transdifferentiation of mature PLs to CCs (Leitão and Sucena 2015) are not incompatible with the data but are not detected in our study since under homeostatic conditions, such events are likely to be of low probability.

### Developmental metabolism and the transcriptome

The pleiotropy of metabolic genes makes it difficult to study their function by the usual loss and gain of function genetic means. This also makes it difficult to understand how development, in many instances, can be driven by metabolism, rather than the commonly held view that metabolic genes exclusively function as bystanders that support development. Based on a plethora of cancer studies as well as on our own analysis of mouse embryonic development, we have realized that co-regulation at a transcriptomic level of a set of metabolic genes that all belong to the same pathway is a good indication that they function in “developmental metabolism”, distinct from their non-specific requirements for cell survival. This process of transcriptional control is particularly apparent for metabolic pathways since their constituent genes are often co-regulated by a small set of factors such as Myc, Hif, and PGC-1alpha. This premise of cell-specific regulation of metabolism is quite dramatically seen in a few key cases shown here. A further mechanistic understanding of these data will require coupling them with metabolomic analysis that is beyond the scope of this study due to the very small tissue size of hand-dissected lymph glands. However, this should become possible in the near future. Currently such analyses in *Drosophila* are usually restricted to tissue from whole animals (reviewed in Cox et al. 2017). As for classical genetic analysis, the fatty acid oxidation pathway has been studied in some detail in the lymph gland (Tiwari et al. 2020).

The analysis presented in this paper demonstrates, for the first time in *Drosophila* hematopoiesis, that cells within individual zones are not only defined by their position within the organ and the markers that they express, but also by their metabolic status. The transcriptomic data can, in many instances, foreshadow how the metabolic profile of an individual cell promotes its progression through development. The PSC cells, as a group, for example, are well represented by most glycolysis related genes that are unlikely to play a bioenergetic role. Instead, the results point to a NADPH related effect linked to the requirement of low ROS for the PSC in the absence of immune challenge (Sinenko et al. 2012; Benmimoun et al. 2015; Louradour et al. 2017). Interestingly, the immediately adjacent MZ cells are quite different from the PSC in their metabolism and they maintain a physiologically relevant high ROS status Different cell types in the lymph gland have some measures of uniqueness in their marker-based as well as metabolic fingerprint over others. This includes the IZ, shown through direct visualization, isolation, FACS analysis, and transcriptomic clustering, to be a uniquely identified cell type. Surprisingly, they also show interesting metabolic signatures since the IPs are highly enriched for both synthesis and clearance of free ceramide from a cell. This is important given the role of ceramide in the activation of the JNK pathway and is indicative of ceramide-triggered transient activation of JNK and MMP1 in this narrow group of cells.

### The transcriptome in mechanistic interpretation of genetic data

Developmentally relevant genes usually function in a context-dependent manner, and the genetic tools in *Drosophila* are particularly well-suited to provide mechanistic insights when they are combined with transcriptomic data. For example, quantitation of the widely expressed *pointed* gene proved invaluable in dissecting its role in individual clusters and showed that it is required for entry into and exit from the IZ state, followed by a role in inhibiting CC and promoting plasmatocyte fate. Similarly, a comprehensive mechanistic analysis of crystal cell specification and maintenance would not have been possible without the identification of an alternative isoform of *sima*, and the transcriptional pattern of both *numb* and *musashi*. On the other hand, deciphering the functional relevance of these patterns required detailed classical genetic analysis.

In summary, genome-wide transcriptomic analysis not only provides a means to identify cell-type specific markers, but when combined with the powerful molecular genetic tools available in *Drosophila,* and increasingly so in mammalian systems, is useful for acquiring mechanistic and functional insights into developmental processes. Here we have combined these approaches to illustrate the continuing relevance of the use of *Drosophila* as the only invertebrate hematopoietic model that provides a logical framework within which to establish less-studied concepts such as the characterization of transitory populations, roles of developmental metabolism, non-canonical Notch signaling, and genetic dissection of pleiotropy.

## Supplementary Files

**Supplementary file 1:** Differentially expressed genes in scRNA-Seq data

**Supplementary file 2**: Comparison of clusters from this study with those identified as clusters by Cho et al., 2020

**Supplementary file 3:** Differentially expressed genes in bulk RNA-Seq data

**Table 1:** MZ_IZ_PL Bulk RNA-Seq differentially expressed genes

**Table 2:** PL_iCC_mCC Bulk RNA-Seq differentially expressed genes

**Supplementary file 3:** Gene lists used in this study for AUCell analysis

## Methods

### Drosophila strains

The *Drosophila* lines from our lab stock were used in this study as follows: *dome^MESO^>GFP Hml^Δ^-DsRed*, *lz-GAL4*, *dome^MESO^-GFP.nls Hml^Δ^-DsRed.nls*, *CHIZ-GAL4 UAS-mGFP* (IZ-specific GAL4, (Spratford et al. 2020)), *Tep4>GFP Hml^Δ^-DsRed*, *Tep4-QF2* (first used in this study), *Hml>2xEGFP*, and *UAS-Notch^ACT^*. The following lines were obtained from the Bloomington *Drosophila* Stock Center (BDSC): *lz-Gal4 UAS-mGFP* (#6314), *UAS-pnt^RNAi^* (#35038), *UAS-hep^ACT^* (#9306), *UAS-mihep* (#35210), *UAS-numb^RNAi^* (#35045), *LexAop2-6XmCherry* (#52271), *UAS-Notch^RNAi^* (#7077), *Msi-GFP* MiMIC protein trap (#61750), *10XQUAS-6XmCherry* (#52269). The following lines were kind gifts from other labs: *bnl-LexA* from Dr. Roy (Du et al. 2017), *UAS-msi^RNAi^ UAS-Dcr2* from Dr. Wappner, (Bertolin et al. 2016), *Hml^Δ^-DsRednls* from Dr. Brückner (Makhijani et al. 2011).

### Preparation of single cell suspension from larval lymph glands

Larvae were collected at the third instar stage and washed with DEPC water on a shaker to remove food traces before dissection. Pairs of lymph gland primary lobes were dissected from 11 larvae, including 6 females and 5 males, (for single cell RNA sequencing) and approximately 100 larvae (for bulk RNA sequencing) in 1X modified dissecting saline (MDS) buffer (9.9 mM HEPES-KOH, 137 mM NaCl, 5.4 mM KCl, 0.17 mM NaH_2_PO_4_, 0.22 mM KH_2_PO_4_, 3.3 mM Glucose, and 43.8 mM Sucrose, pH 7.4) and lymph glands were then placed into a glass dish containing Schneider’s medium (Gibco) kept on ice. Glass dishes were pretreated with 1% BSA in PBS and rinsed prior to use to prevent adherence of primary lobes. Three biological replicates were done in parallel. The lymph glands were dissociated as previously described with some modifications (Harzer et al. 2013; Khan et al. 2016). After being washed with MDS buffer twice, these tissues were transferred to 1.5 mL DNA LoBind tubes (Eppendorf) and incubated with 200 µL of dissociation solution containing 1 mg/mL of papain (Sigma, P4762) and 1 mg/mL of collagenase (Sigma, C2674) in Schneider’s medium. They were dissociated for 15 minutes in a shaking incubator at 25°C, 300 rpm. Next, 500 µL of cold Schneider’s medium was added and the suspension was gently pipetted up and down using a low-binding 1000 µL tip (Olympus Plastics) 20 times for mechanical dissociation. After centrifugation at 3000 rpm for 5 minutes, cells were resuspended and washed with 500 µL of 1X PBS (Corning, MT21040CV) containing 0.04% of UltraPure BSA (Invitrogen, AM2616) and then passed through a 35 micrometer cell strainer (Falcon 352235). For preparation of the single cell RNA-Seq sample, the cell suspension was concentrated by centrifuging and resuspending in a lower volume of PBS containing 0.04% BSA (30 µL). Cell concentration and viability were assessed using the Countess II automated cell counter (Applied Biosystems). The samples with final concentration of more than 650 cells/µL and viability of more than 85% were used for single cell RNA-Seq.

### Flow cytometry and cell sorting

For all RNA extractions from sorted populations, dissociated live lymph gland cells were sorted using the BD FACSARIA-H. Gates and compensation were based on single color controls. Cells were sorted into 300 µL of DNA/RNA Shield (Zymo) in DNA LoBind tubes and frozen at -80°C prior to RNA extraction.

For DNA content analysis, dissociated lymph gland cells were fixed in 1 mL of 1% formaldehyde solution in PBS after being dissociated using the above protocol. Cells were incubated in fixative in low binding tubes for 30 minutes at 4°C on a shaker, then were spun down and washed with PBS. Fixed cells were resuspended in a solution of PBS containing NucBlue™ Live ReadyProbes™ Reagent (Hoechst 33342) and incubated at room temp on a shaker for 30 minutes. Cells were transferred to 5 mL polystyrene tubes for flow cytometry analysis on the BD LSRII. Cells were gated to exclude doublets using the FSC-H vs FSC-W and SSC-H vs SSC-W comparisons.

### RNA extraction and qRT-PCR

For the bulk RNA-Seq and qRT-PCR, total RNA was extracted using the Quick-RNA Microprep Kit (Zymo). RNA quality control was performed using the Agilent 4150 TapeStation system. For qRT-PCR analysis, cDNA was generated using SuperScriptIV VILO Master Mix (Thermo Fisher). qRT-PCR was performed on cDNA using the PowerUp SYBR Green Master Mix (Thermo Fisher) and the StepONE Real Time PCR system (Applied Biosystems). Primers used were as follows: sima-RA/RB Forward 5’-GCAGAACTTCAAGGTGCAATAA-3’; sima-RA/RB Reverse 5’-CACCGTTCACCTCGATTAACT-3’; sima-RC Forward 5’-GAGGCGCACTAGTGACAAA-3’; sima-RC Reverse 5’-CGAGCGAGATAGCAACGG-3’.

### Bulk RNA-Sequencing and analysis

cDNA libraries were prepared using KAPA Stranded mRNA-seq kit (KAPA Biosystems) for the MZ/IZ/CZ bulk RNA-Seq experiment and the Universal Plus mRNA-Seq kit (Nugen) for the crystal cells bulk RNA-Seq. Libraries were sequenced on two lanes of a HiSeq3000 (Illumina) or NovaSeq 6000 SP (Illumina), respectively. RNA sample and cDNA library concentration and quality control was assessed using the Agilent 4200 TapeStation system. Sequencing data was analyzed using Partek Flow, a web-based software platform. Sequences were aligned to the *Drosophila melanogaster* reference genome r6.22 (Flybase) using STAR aligner with default parameters. Read counts were normalized by counts per million (CPM). Differential genes expression analysis was performed using ANOVA with a fold change cutoff of 2 and FDR<0.05 (see Supplementary file 3 Tables 1 and 2).

### Single cell RNA-Sequencing

Three samples were processed using 10X Single Cell 3’ GEX version 3 (10X Genomics) and sequenced on a NovaSeq 6000 S4 PE (Illumina) at UCLA Technology Center for Genomics & Bioinformatics. Approximately 8,400 cells per sample were put through the Chromium Controller Instrument (10X Genomics) to partition single cells into Gel bead-in-Emulsions (GEMs) and 10x barcoded libraries were then constructed. All cDNA libraries were sequenced on one lane of NovaSeq flow cell using a symmetric run (2×150bp).

### Single cell RNA-Seq data processing and analysis

Partek Flow was used to analyze single cell sequencing data. Before alignment, each paired read was trimmed according to 10x Genomics Chromium™ Single Cell 3’ v3 specifications. Trimmed reads were aligned as described in the bulk RNA-Seq above. After alignment, UMIs were deduplicated and barcodes were then filtered and quantified to generate a single cell count matrix. Genes with fewer than 30 total reads across three samples were excluded. Low-quality cells and potential doublets were filtered out by selecting out cells with high read counts (>85,000), an especially low (<1500) or high (>4500) number of genes detected, or a high percentage of mitochondrial reads (>6%). 21,157 cells passed these quality control filters across the three samples.

Read counts were normalized with the following order: (1) CPM; (2) Add 1; (3) Log2 transformation. Genes that were not expressed in any cells were excluded and a total of 9,458 genes were detected. Data was then corrected for batch effects between samples. All rRNA and three sex-related lncRNA (*lncRNA:roX1*, *lncRNA:roX2*, and *lncRNA:CR40469*) were filtered out. The data was imputed using the MAGIC algorithm (van Dijk et al. 2018) with the number of nearest neighbors =50.

For graph-based clustering, the top 2000 genes with the highest dispersion were used to perform principal components (PC) analysis (PCA). Based on the Scree plot, the first 13 PCs were selected to use as the input for clustering and data visualization tasks. Graph-based clustering was first performed with 50 nearest neighbors (NN) and resolution (res) of 0.1 - 0.5, giving 6-14 clusters, with 8 clusters (res=0.19) showing the most marked differences in gene expression between clusters. Each cluster was compared to the others using an ANOVA to identify differentially expressed genes which were enriched more than 1.5-fold (with a false discovery rate cutoff FDR<10xe-6) (see Supplementary file 1). Cluster identity was assigned based on the presence of known zone-specific markers in the differentially expressed genes. Gene set enrichment analysis (GO, KEGG) was performed on the differentially expressed genes for each cluster. The data was visualized using t-Distributed Stochastic Neighbor Embedding (t-SNE) with perplexity of 50 and PCA initialization.

To evaluate the developmental progression of the cells from a progenitor to a differentiated cell type, a subsequent trajectory and pseudotime analysis using Monocle 2 (Qiu et al. 2017) was performed. PSC and X clusters were both excluded from trajectory analysis. The trajectory was calculated using the top 1824 genes with the highest dispersion. For cluster X, a separate trajectory was constructed using the top 1500 highest dispersion genes.

To analyze the activity of a gene set in our data, we used the AUCell tool (Aibar et al. 2017), with the top 25% of genes. The minimum gene set size was generally five, but in a few cases where the number of genes in the pathway was between three and five, a smaller minimum was used. Gene lists used for AUCell analysis are found in Supplementary file 4.

We performed subclustering on isolated clusters (CC, X and PL) to determine subpopulations. The subclustering analysis follows the general procedure as the initial graph-based clustering and used 2000 genes with the highest dispersion within each of the isolated clusters. Two CC subclusters were identified using 5 PC, 60 NN, and res 0.1. Three X subclusters were generated using 4 PC, 30 NN, and res 0.5. PL showed four subclusters with 7 PC, 50 NN, res 0.2.

### Lymph gland dissection, immunostaining and imaging

Lymph glands were dissected and processed as previously described (Jung et al. 2005). Briefly, for MMP1 staining, lymph glands were dissected and immediately placed into 4% formaldehyde in PBS on ice and then fixed for 15 minutes at room temp. For all other stainings, lymph glands were dissected into cold PBS and then fixed in 4% formaldehyde in PBS at room temp for 20 minutes. After fixation, tissues were washed three times in PBS with 0.3% Triton X-100 (PBST) for 10 minutes each, blocked in 10% normal goat serum in PBST (blocking solution) for 30 minutes, followed by incubation with primary antibodies in blocking solution. Primary antibodies were incubated with tissues overnight at 4°C and then washed three times in PBST for 10 minutes each, followed by incubation with secondary antibodies for 3 hours at room temp. Samples were washed four times in PBST, with DAPI (1:1000, Invitrogen) added to the third wash to stain nuclei, and then placed into VectaShield mounting medium (Vector Laboratories) and mounted on glass slides. Notch internalization assays were performed as described (Mukherjee et al. 2011) using mouse anti-NECD (1:50, DSHB C458.2H). Lymph glands and dissociated cells were imaged using a Zeiss LSM 880 Confocal Microscope. Image files were prepared and analyzed using the Fiji and Imaris software.

The primary antibodies were used as follows: mouse anti-MMP1 (1:100 of a 1:1:1 mixture of 3B8D12, 3A6B4 and 5H7B1, DSHB) (Page-McCaw et al. 2003), rabbit anti-PPO2 (1:200, kind gift of Dr. Asano) (Asano and Takebuchi 2009), mouse anti-Hnt (1:200, DSHB 1G9-c) (Yip et al. 1997), guinea pig anti-Numb (1:200, kind gift of Jan lab) (Roegiers et al. 2001), preabsorbed rabbit anti-Numb (1:200, kind gift of Jan lab) (Rhyu et al. 1994), guinea pig anti-Sima (1:100) (Wang et al. 2016), mouse Hrs 8-2 antibody (DSHB) (Riedel et al. 2016). Primary antibodies were detected with secondary antibodies conjugated to Cy3 (1:100; Jackson ImmunoResearch Laboratories) or Alexa 633 (1:100), Alexa 555 (1:200), or Alexa 488 (1:200) (Invitrogen).

## Acknowledgements

We thank Fangtao Chi for engineering the *CHIZ-GAL4* construct used in this study, and past and present members of the laboratory for their help and advice. We gratefully acknowledge FlyBase; the Bloomington *Drosophila* Stock Center; the Vienna *Drosophila* Resource Center; the Kyoto Stock Center, and the fly community including Sougata Roy, Pablo Wappner, Dalmiro Blanco-Obregon, Katja Brückner, Tsunaki Asano, Yuh Nung Jan, Dirk Bohmann, and Andrea Page-McCaw for reagents. We acknowledge the help of the Broad Stem Cell Research Center (BSCRC) and Owen Witte for help and support, the MCDB/BSCRC Core Facility in Microscopy and the BSCRC core in Flow Cytometry, particularly Felicia Codrea, Jessica Scholes, and Jeffrey Calimlim for help with cell sorting. We thank the UCLA TCGB center, Xinmin Li, and Michael Mashock for help in sequencing. We acknowledge the Partek Flow technical support team and in particular Xiaowen Wang’s help, which was crucial for data analysis. We thank undergraduate research scholars Peiliang Zhou, Chloe Su, and Khoi Luc for their contributions. We thank David Eisenberg and Vy Phan Lai for their support of D.M.V through the Center for Global Mentoring. U.B. is supported by National Institutes of Health grants R01 HL-067395 and R01 CA-217608; J.R.G. by Ruth L. Kirschstein National Research Service Award number T32HL69766 and UPLIFT (UCLA Postdocs’ Longitudinal Investment in Faculty) Award number K12GM106996; L.M.G. by Ruth L. Kirschstein Institutional National Research Service Award number T32CA009056 and by National Heart, Lung, and Blood Institute of the National Institutes of Health under award number 3R01HL067395-16S1; D.M.V. by the Center for Global Mentoring at UCLA-DOE Institute for Genomics & Proteomics; and C.M.S. by Ruth L. Kirschstein National Research Service Award number T32HL863458.

---

**Figure 1—figure supplement 1:**
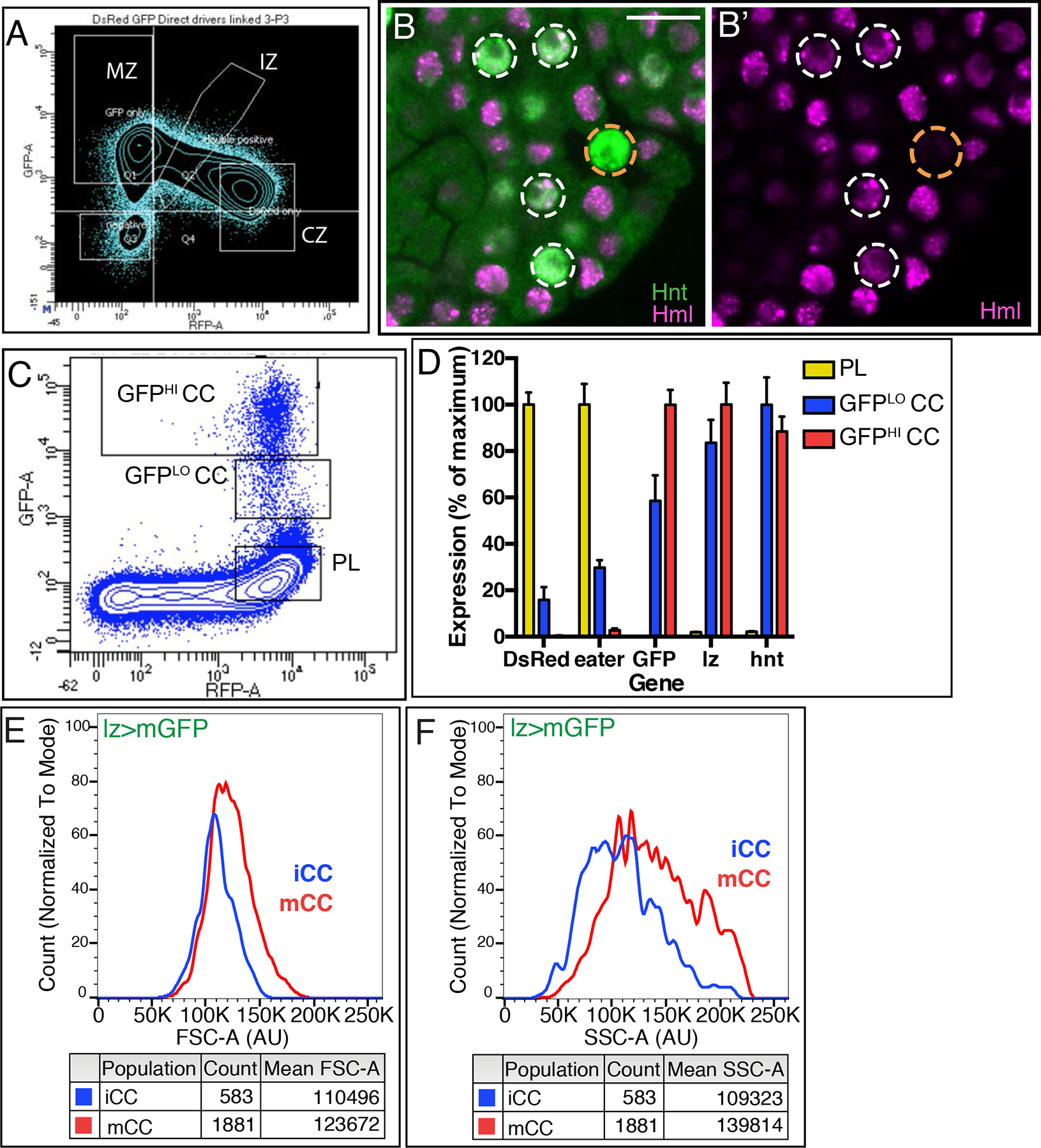
Resolving heterogeneity in sorted populations. **(A)** Flow cytometric analysis of dissociated lymph gland cells gated into four populations: MZ (high GFP, low DsRed), CZ (low GFP, high DsRed), IZ (intermediate levels of both GFP and DsRed), and double negatives (low GFP, low DsRed). **(B,B’)** Confocal image of *Hml^Δ^-DsRed.nls* (magenta) lymph gland stained with anti-Hnt antibody (green). As is more clearly evident in **B’**, several Hnt+ crystal cells express low levels of *Hml* (white circles). *Hml(DsRed)* is not seen in the highest Hnt expressing crystal cells (orange circle). Genotype in **C-F**: *lz-GAL4*, *UAS-mGFP*, *Hml^Δ^-DsRed.nls*. **(C)** Flow cytometry gates employed to distinguish between GFP^HI^ CC, GFP^LO^ CC, and PL populations used for data shown in Figure 1F-I. **(D)** Gene expression analysis from bulk RNA-Seq. *Hml* (*DsRed*) and *eater* are high, but *lz* and *hnt* are virtually missing from PL. All CCs are characterized by (*lz>*) *GFP*, also by *lz* and *hnt*. The difference between GFP^LO^ and GFP^HI^ CCs is more apparent from *GFP* (and *PPO1/2*, see main Figure) transcript level than the expression of *lz* or *hnt* transcripts, which are similar between the two populations. **(E)** Mean FSC-A, and therefore average cell size, is larger for mCCs than for iCCs. **(F)** Mean SSC-A, a measure of intracellular complexity, is smaller for iCCs than for mCCs.

**Figure 2—figure supplement 1:**
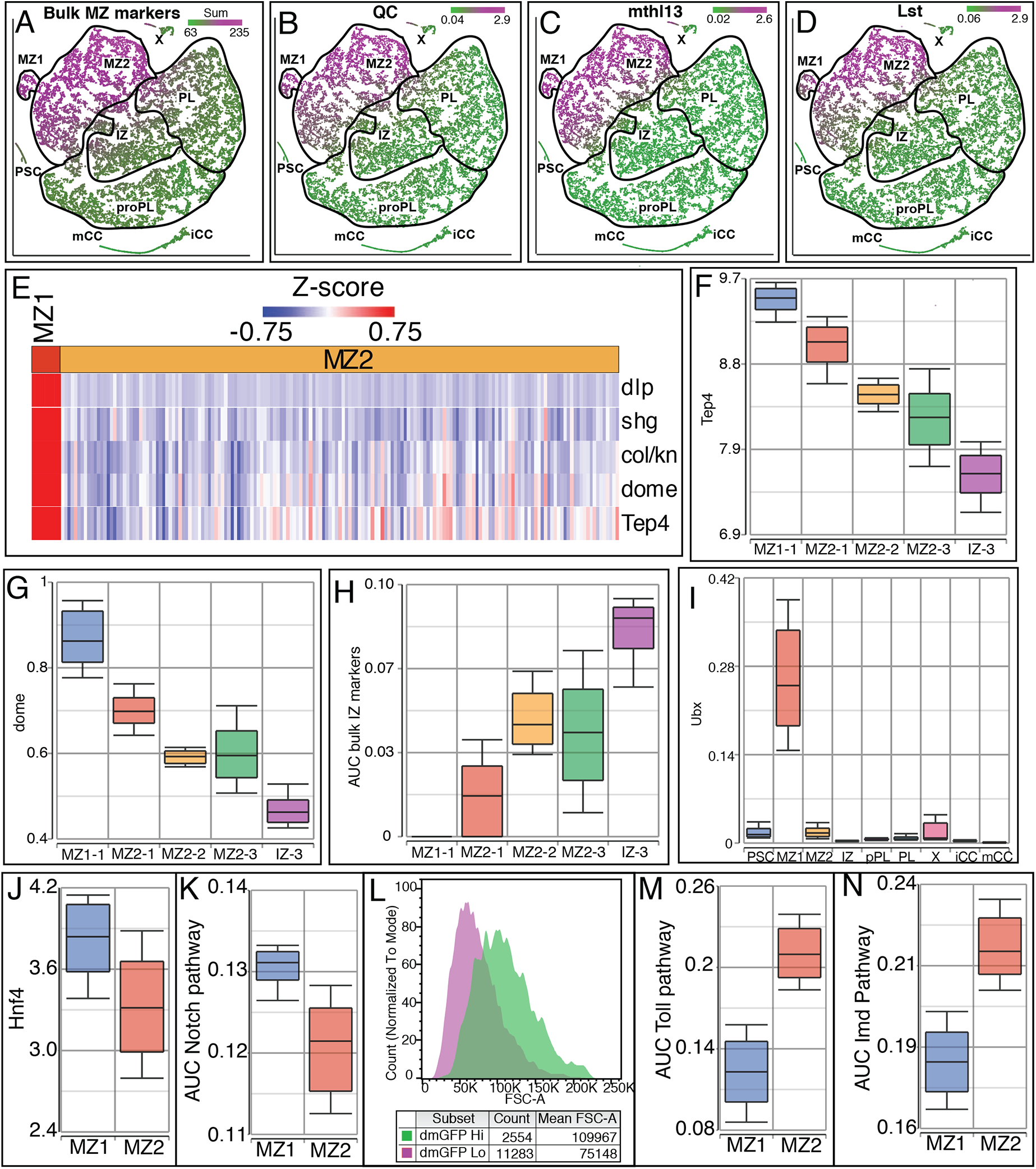
Comparison of MZ1 and MZ2 cluster characteristics. **(A)** Gene list identified as MZ enriched in bulk RNA-Seq data is used here to obtain individual expression values in single cell RNA-Seq experiments. A sum of these expression values mapped onto the t-SNE plot highlights MZ1 and MZ2. These data point to the overall equivalency of results between the bulk RNA-Seq and single cell RNA-seq data. **(B-D)** Expression of newly identified MZ-specific genes *QC* **(B),** *mthl13* **(C)** and *Lst* **(D)** are high in MZ1 and MZ2 but low elsewhere. **(E)** MZ1-specific genes with low MZ2 expression. **(F, G)** Progressive decline of *Tep4* **(F)** and *dome* **(G)** expression from MZ1-1 onwards through IZ-3. **(H)** AUCell analysis of the IZ-enriched markers derived from bulk RNA-Seq shows no expression in MZ1, progressively increasing through IZ-3. **(I)** *Ubx* is most highly expressed in MZ1 compared to all other clusters. **(J-K)** Enrichment of MZ1 expression compared to MZ2 is seen for *Hnf4* **(J)** and Notch pathway components (AUCell) **(K)**. **(L)** When *dome^MESO^>EGFP* positive MZ cells are split into high-GFP (dmGFP Hi) and low-GFP (dmGFP Lo) expressing populations, flow cytometric analysis shows that lower maturity (high-GFP) cells show larger cell size (FSC-A) than higher maturity (low-GFP) cells. **(M-N)** AUCell analysis of MZ1 and MZ2 clusters shows that the Toll pathway **(M)** and Imd pathway **(N)** are more enriched in MZ2 than in MZ1.

**Figure 2—figure supplement 2:**
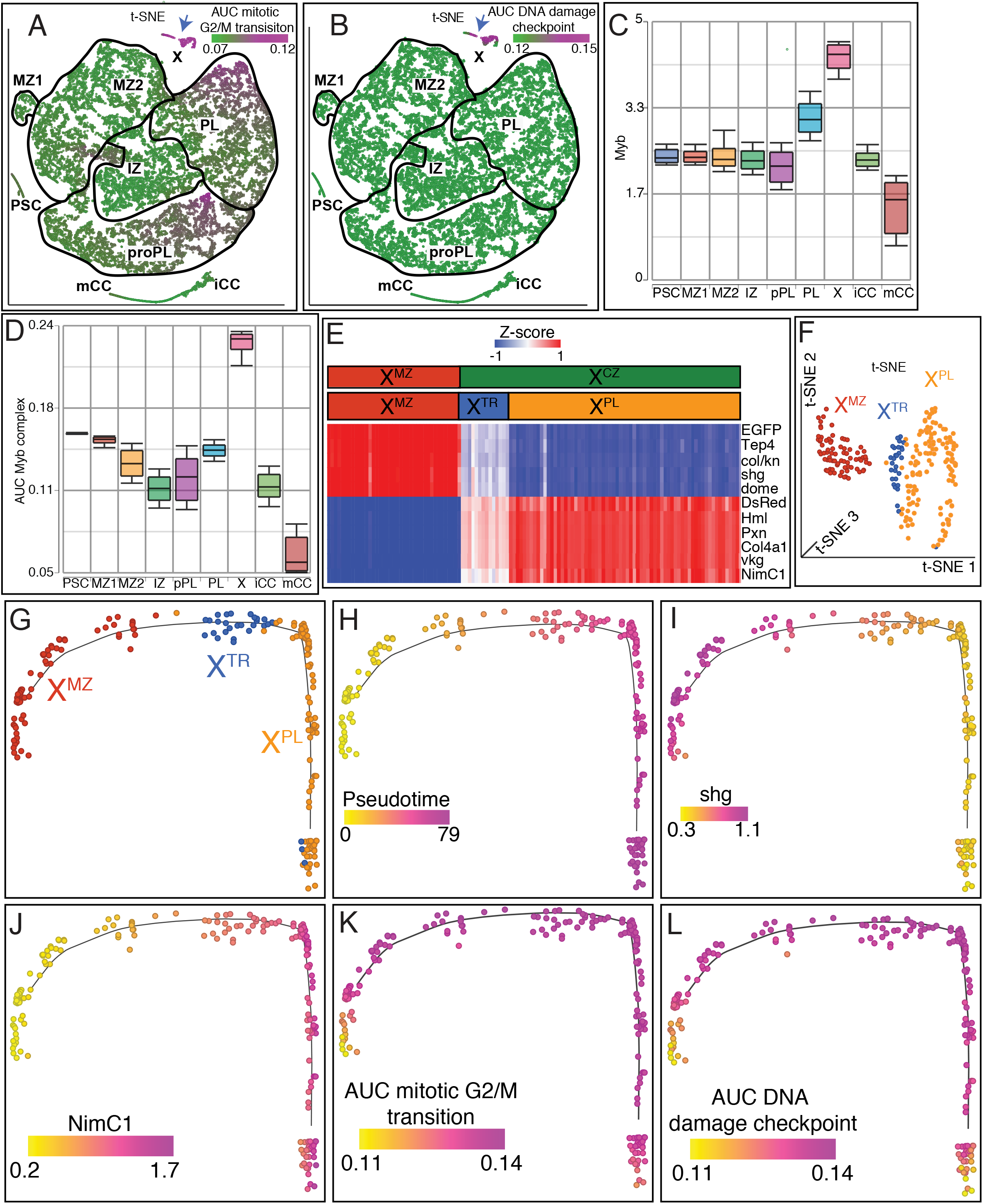
Characteristics of Cluster X. Cluster X cells (blue arrow) show the highest enrichment of mitotic G2/M transition genes **(A),** intra-S DNA damage checkpoint genes **(B)**, *Myb* **(C),** and genes encoding components of the Myb complex **(D)**. **(E)** The two islands of X (X^MZ^ and X^CZ^), which split into a total of three subclusters (X^MZ^, X^TR^, and X^PL^) show distinct expression patterns of hallmark zone-specific genes. X^MZ^ cells express high levels of MZ and low levels of PL genes. X^PL^ cells display high expression of PL and low expression of MZ genes. X^TR^ cells exhibit moderate expression of both MZ and PL genes. **(F)** Graph-based clustering of X shown on a 3-dimensional t-SNE plot. X is broken up into 3 clusters: X^MZ^, X^TR^, and X^PL^. **(G-L)** Trajectory analysis of isolated cluster X cells. Superimposed graph-based clustering **(G)** and pseudotime analysis **(H)** shows that the trajectory of X follows the sequence X^MZ^, X^TR^, X^PL^. **(I)** The MZ marker *shg* (*E-Cad*) is high at the beginning of the X trajectory, moderate in the middle and low at the end. **(J)** The CZ marker *NimC1* is low at the beginning of the X trajectory, moderate in the middle and high at the end. **(K-L)** In contrast to the MZ and CZ related genes, mitotic G2/M transition **(K)** and intra-S DNA damage checkpoint **(L)** related genes are distributed fairly uniformly over the entire trajectory.

**Figure 2—figure supplement 3:**
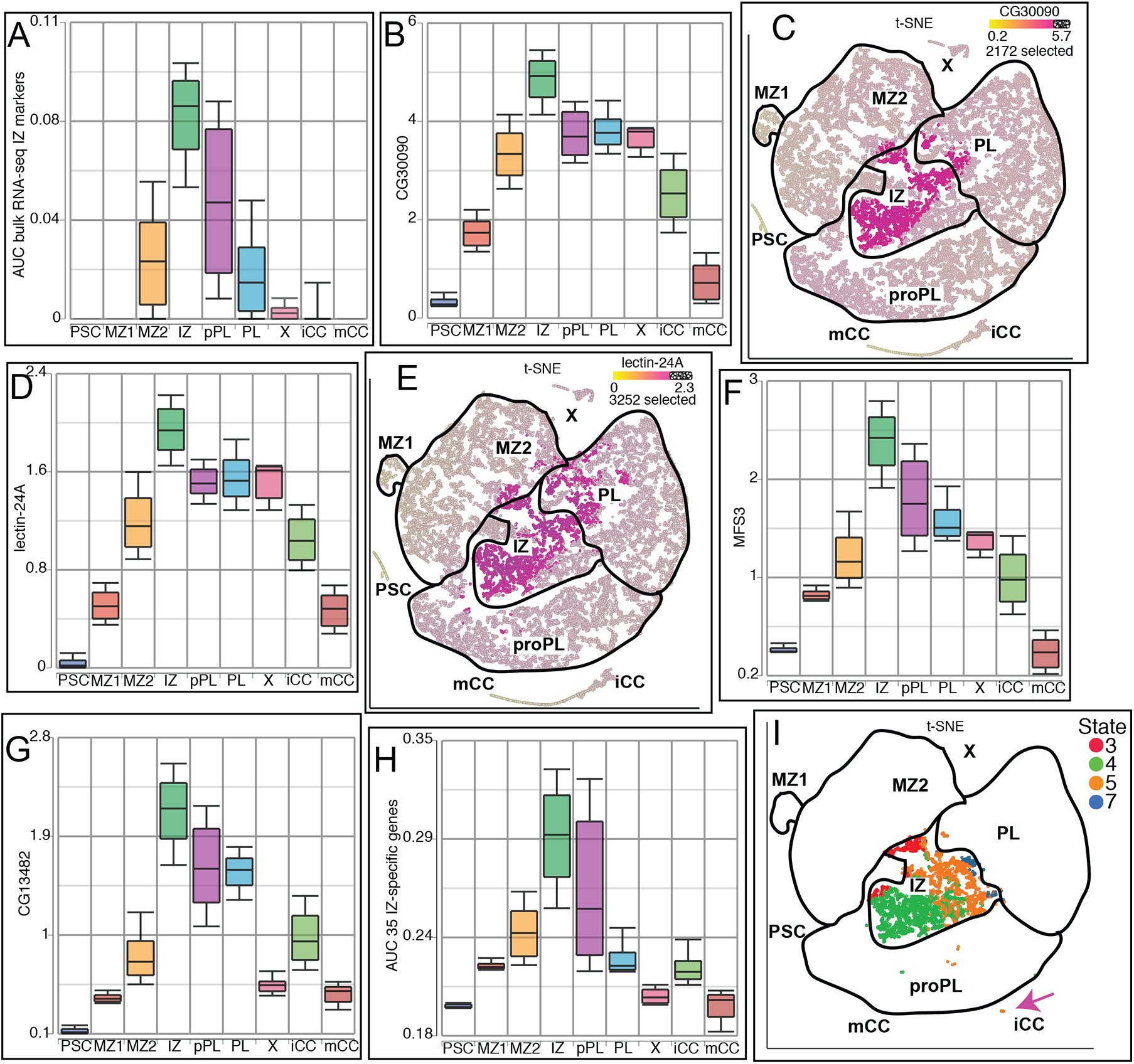
IZ-enriched markers identified in bulk RNA-Seq analysis mapped for their expression in single cell experiments. **(A)** As a gene set (AUCell), these genes show highest enrichment in IZ, with lower levels in proPL, MZ2, and PL. All others have near 0 AUC level. **(B-G)** Expression of individual IZ-enriched genes. **(B-C)** *CG30090* is highest in IZ on **(B)** an expression plot and **(C)** when visualized on a t-SNE that has been thresholded for high *CG30090* expression. **(D-E)** The same is true for *lectin-24A*. **(F-G)** Expression of *MFS3* **(F)** and *CG13482* **(G)** is highest in the IZ, but these genes also overlap somewhat with proPL. **(H)** AUCell analysis of the 35 IZ-specific differentially expressed genes identified in scRNA-Seq are highest in IZ compared to the other clusters, but some proPL cells also show high expression. **(I)** t-SNE plot showing the different trajectory states that IZ cells belong to. IZ-3 cells are found at the MZ border, while IZ-4 cells are located further down in the IZ cluster and border both IZ-3 and IZ-5 cells. IZ-5 cells are central within the IZ and border IZ-4 and IZ-7. A small number of IZ-5 cells are located on the crystal cell island (pink arrow). IZ-7 cells border the PL cluster.

**Figure 2—figure supplement 4:**
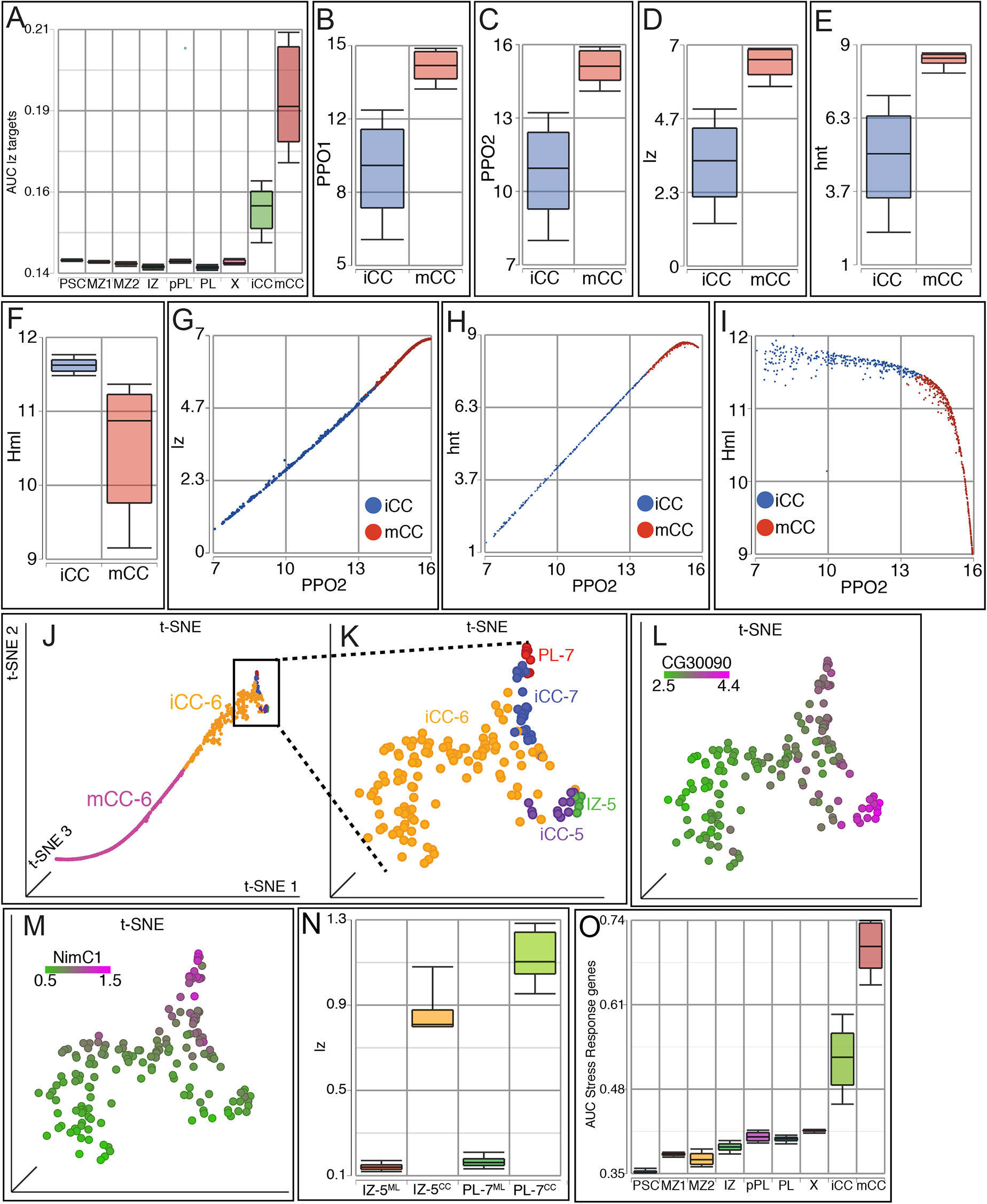
Crystal cell genes. CCs have high expression, and within CCs, mCCs have higher expression than iCCs, for Lz target genes **(A)**, *PPO1* **(B)**, *PPO2* **(C)**, *lz* **(D)** and *hnt* **(E)**. **(F)** In contrast, *Hml* shows lower expression in mCC than in iCC. **(G-H)** *PPO2* levels are highly and positively correlated with *lz* **(G**; r=1) and *hnt* **(H**; r=0.99) in both iCCs and mCCs. **(I)** In contrast, *Hml* is negatively correlated with *PPO2* levels, especially in mCCs. **(J-K)** 3D t-SNE visualization of crystal cell island showing subclusters and their cluster-state classifications. **(J)** The entire CC island showing the relative locations of iCC-6 and mCC-6 cells that make up the majority of CCs, with the base of the CC island showing small populations of cells from other clusters. **(K)** A magnified view of the CC island (boxed part of **J**). The small non-overlapping iCC groups, iCC-5 and iCC-7, border iCC-6 on opposite arms. IZ-5 cells border iCC-5 cells, while PL-7 cells border iCC-7 cells. **(L-M)** Zone-specific gene expression in cells shown in **(K)**. Compared to other nearby cells, **(L)** the IZ marker *CG30090* is expressed highest in the cells in the lower right corner, which include IZ-5 and iCC-5. **(M)** The PL marker *NimC1* is expressed at higher levels in the cells in the upper right corner which include PL-7 and iCC-7. **(N)** IZ-5 and PL-7 cells on the CC island (IZ-5^CC^ and PL-7^CC^, respectively) have higher levels of *lz* expression compared to the corresponding populations from the “mainland” area of the t-SNE (i.e. IZ-5^ML^ and PL-7^ML^) **(O)** AUCell analysis shows that stress response genes are highly enriched in CCs compared to other clusters, with mCC higher still than iCC.

**Figure 2—figure supplement 5:**
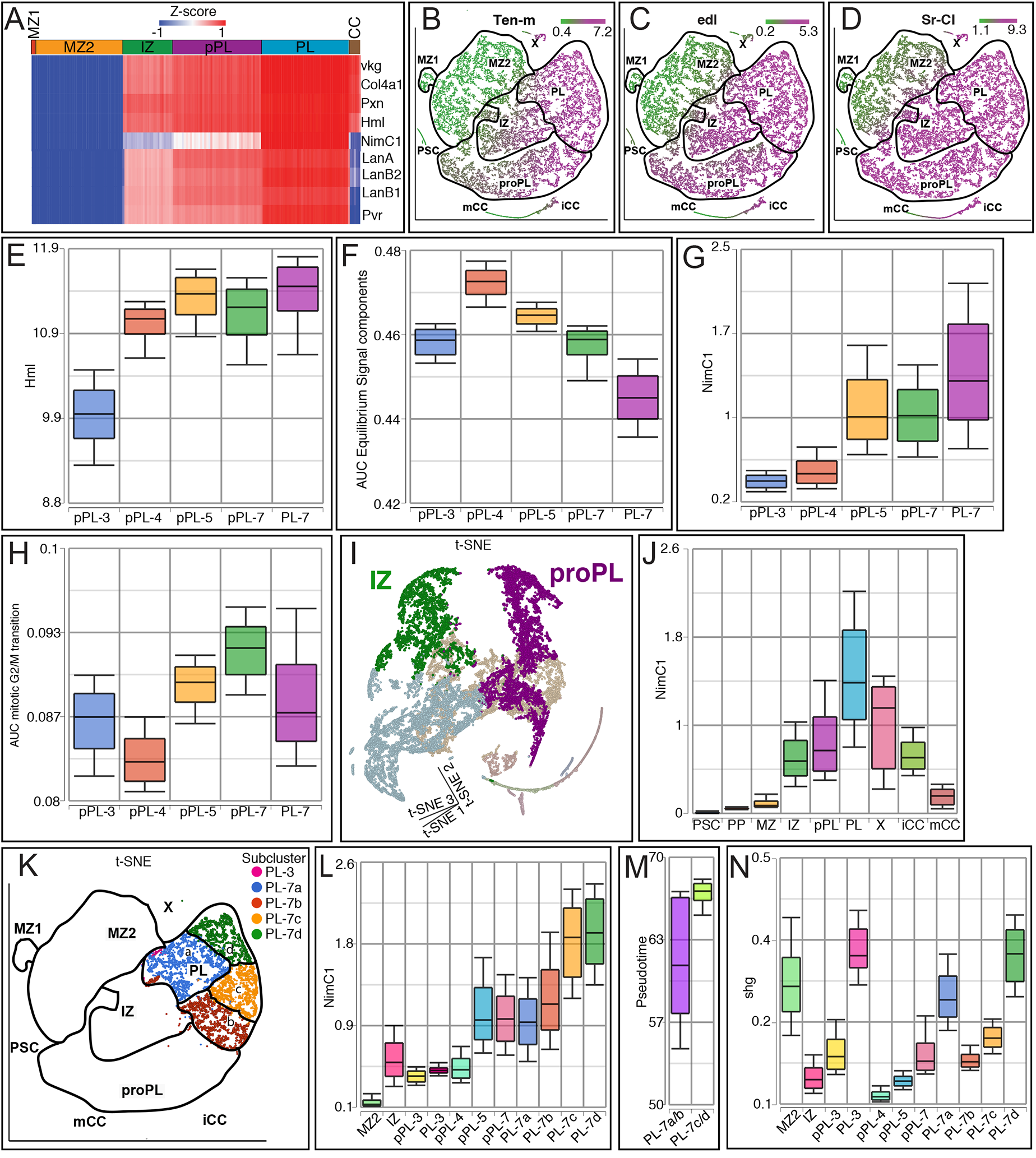
Expression profiling proPL and PL. **(A)** CZ related markers are most highly expressed in PL and have decreased expression in proPL and IZ. *NimC1* is the most significant of these, and is lower in proPL than in PL and not expressed in IZ cells. *NimC1*, *LanA*, *LanB2*, *LanB1*, and *Pvr* are plasmatocyte-specific genes as they are not expressed in crystal cells (CC). **(B-D)** Newly identified CZ marker genes, *Ten-m* **(B)** *edl* **(C)** and *Sr-Cl* **(D)** are highly expressed in IZ, proPL, PL, iCC, and a subset of Cluster X. **(E-H)** Expression of individual genes and enrichment of specific pathways distinguish individual subclusters of proPL and PL. **(E)** *Hml* is expressed at low levels in proPL-3, but is higher in the other proPL and PL subclusters. **(F)** AUCell representing the equilibrium signal gene set is uniquely enriched in the subcluster proPL-4. **(G)** *NimC1* expression is highest in PL-7, lower in proPL-7 and proPL-5, and it is not expressed in proPL-4 or proPL-3. **(H)** mitotic G2/M transition genes are represented slightly higher in proPL-7 compared to the other subclusters. **(I)** 3D t-SNE emphasizes that the IZ and proPL clusters are separated in space and are not connected to each other. Although there are rare instances of connecting cells, the data strongly suggest that IZ and proPL are two independent and parallel means to connect MZ with PL. **(J)** *NimC1* expression is highest in PL compared to all other clusters. **(K-N)** Subclusters of PL. When PL cells are further split by graph-based clustering, four subclusters are found: a, b, c and d. **(K)** t-SNE visualization of the subclusters PL-3, PL-7a, PL7-b, PL-7c, and PL-7d. The PL-7 cluster-state includes cells that are PL (by graph-based clustering) and are in state 7 on the trajectory (late appearing and on a terminal branch). **(L)** PL-7c and PL-7d have the highest expression of *NimC1* than all other subclusters. *NimC1* expression in PL-7a and PL-7b is comparable to that of proPL-5 and proPL-7. **(M)** Graphical representation of pseudotime comparing PL-7a/b and PL-7c/d based on the trajectory in Figure 2. PL-7a/b cells are earlier in pseudotime than PL-7c/d. **(N)** Expression of the MZ marker *shg* (*E-Cad*) differs in the PL populations. PL-3 and the PL-7 subclusters closest to MZ2 (e.g. PL-7a and PL-7d) exhibit high levels of *shg* similar to MZ2 and higher than any other IZ, proPL or PL subcluster.

**Figure 2—figure supplement 6.**
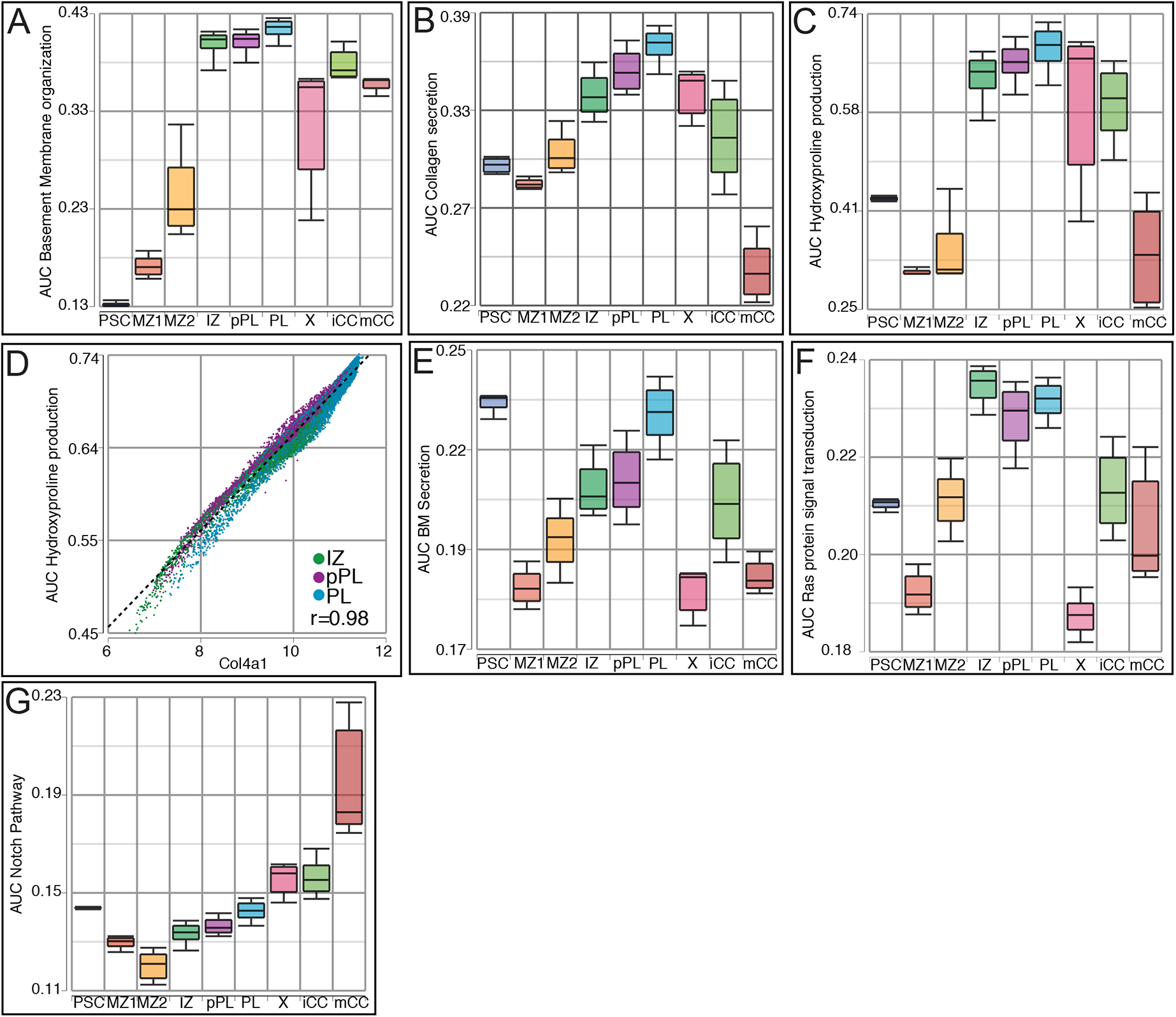
AUCell analysis of ECM and Ras related pathways. Gene sets representing basement membrane organization **(A)**, collagen secretion **(B)**, and hydroxyproline production **(C)** are most highly enriched in IZ, proPL, and PL, with lower representation in X, iCC, and mCC, and lowest in MZ2, MZ1, and PSC. **(D)** *Col4a1* is highly correlated with hydroxyproline production gene expression (AUC) in IZ, proPL, and PL (r=0.98). **(E)** Basement membrane secretion related genes are highly enriched in the PSC and PL. **(F)** Ras pathway related signal transduction genes are highly enriched in IZ, proPL, and PL. **(G)** Notch pathway genes are uniquely enriched in mCC based on AUCell analysis.

**Figure 3—figure supplement 1:**
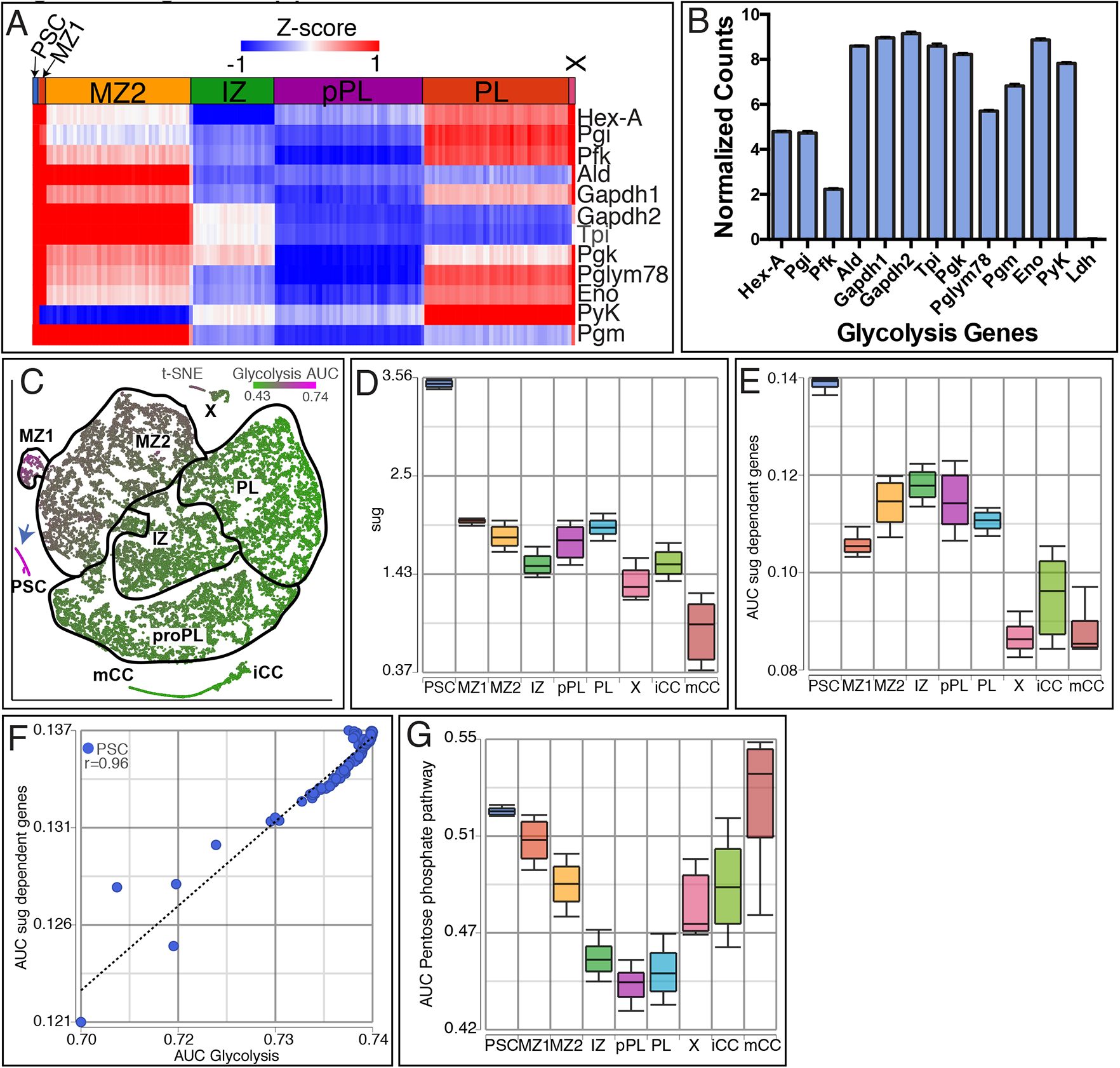
Unique metabolic signatures of the PSC. Data based on single cell RNA-Seq analysis. **(A)** All glycolytic genes (except *Ldh*; see text), are highly expressed in the PSC, only subsets are enriched in the other clusters. MZ1 is the most similar to PSC. **(B)** Average normalized counts within the PSC cluster highlights the lack of *Ldh* expression when compared with other genes in the glycolytic pathway. **(C)** AUC scores for glycolysis genes visualized on a t-SNE highlights the strong enrichment in the PSC (blue arrow). **(D)** *sugarbabe* (*sug*) is highly expressed in the PSC. **(E)** Sug transcription factor target genes are also enriched in the PSC **(F)** Sugarbabe target genes show a strong positive correlation with AUC activity scores for glycolysis in PSC cells (r =0.96). **(G)** The Pentose phosphate pathway shows high expression activity in the PSC relative to other clusters (except mCCs).

**Figure 3—figure supplement 2:**
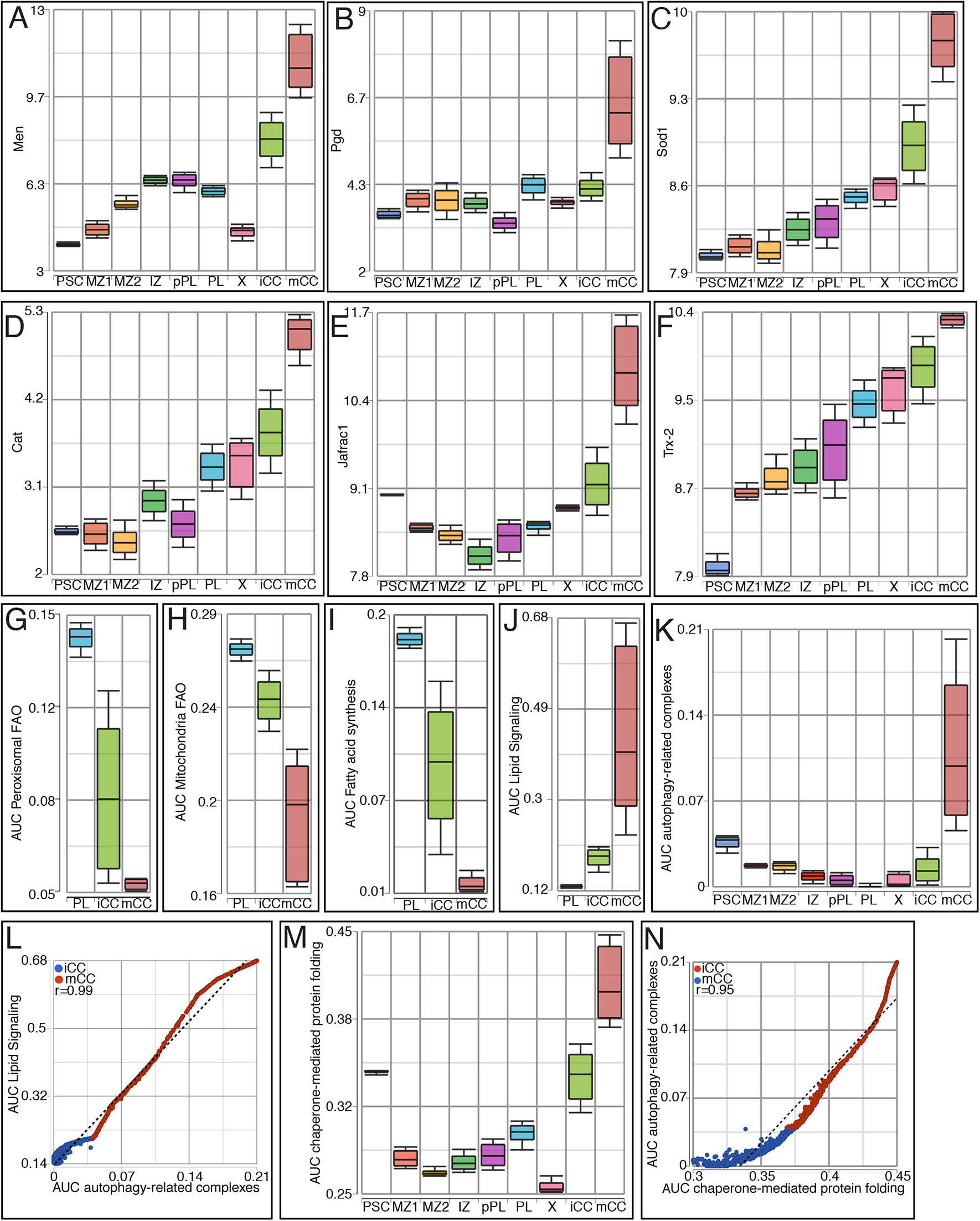
Metabolic changes during CC maturation. **(A-B)** Dehydrogenase enzyme encoding genes involved in NADPH production, *Men* **(A)** and *Pgd* **(B)**, are very highly expressed in mCCs. The iCCs express high *Men* but not *Pgd*. **(C-F)** The ROS scavengers *Sod1* **(C)**, *Catalase* (*Cat*) **(D)**, *Jafrac1* **(E)**, and *Trx-2* **(F)** are also highly expressed in iCC, increasing significantly in mCC. **(G-I)** The activity (AUCell) of fatty acid pathways are low in iCC and lower still in mCC, compared to PL (which is very similar in level to IZ and proPL). This trend holds true for peroxisomal fatty acid beta oxidation pathway **(G)**, mitochondrial fatty acid beta oxidation pathway **(H)**, and the fatty acid synthesis pathway **(I)**. **(J)** Pathways involved in synthesis and regeneration of intracellular lipid signaling components follow the opposite trend. **(K)** Autophagy-related genes show high enrichment in mCCs. **(L)** The gene sets utilized in **(J)** and **(K)** show a strong positive correlation between each other in the CCs (r=0.99). **(M)** Chaperone-mediated protein folding genes are highly enriched in mCCs and **(N)** correlate very well with autophagy-related components (r=0.95).

**Figure 3—figure supplement 3:**
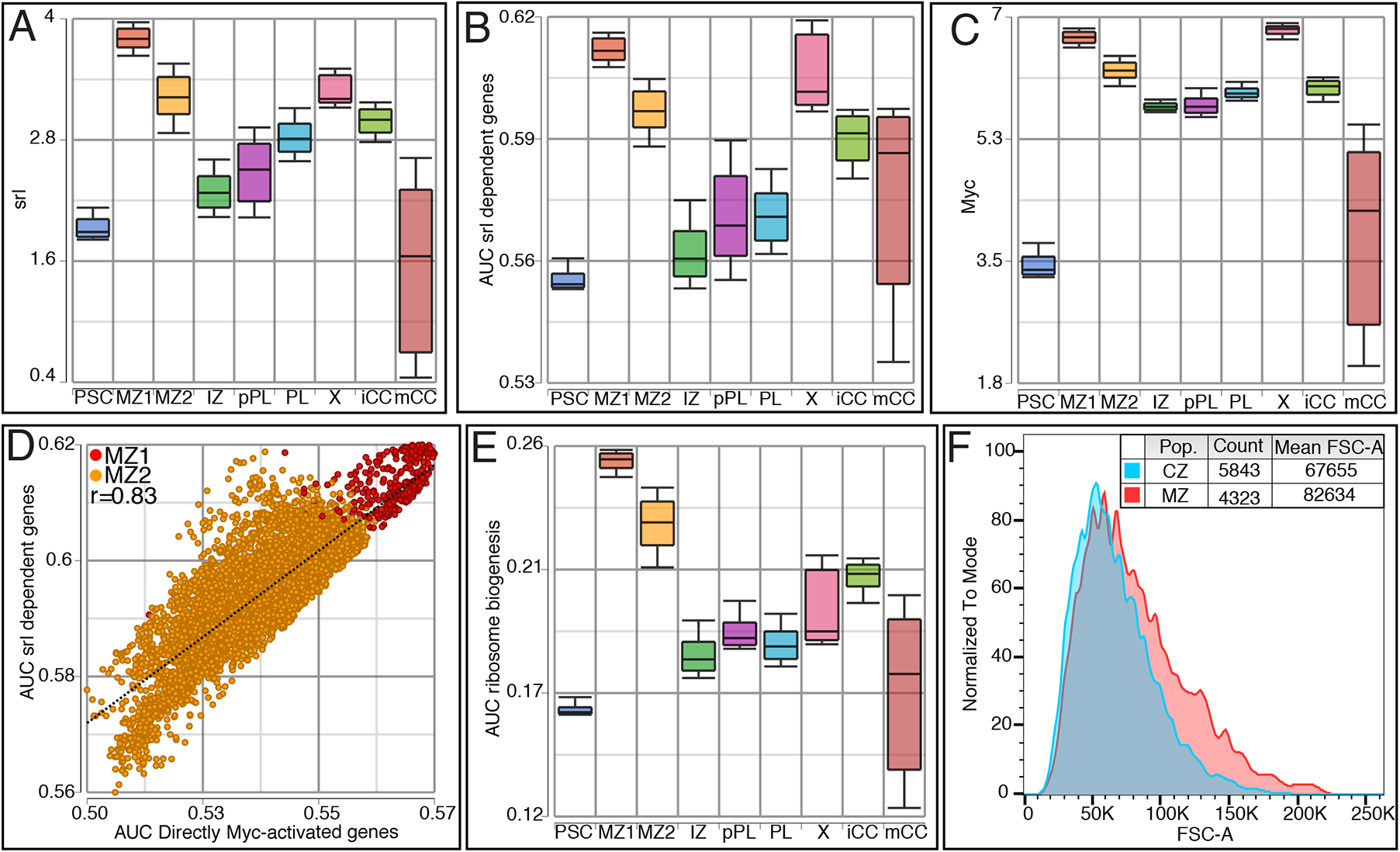
Metabolism of the MZ. **(A-E)** Data based on single cell RNA-Seq analysis. **(A)** *Spargel* (*srl*; *PGC1-alpha*) transcript is highly expressed in MZ1 and MZ2 with higher levels in MZ1. **(B)** AUCell analysis of Srl transcription factor target genes shows enrichment in both MZ1 and MZ2 progenitors. **(C)** Expression of *Myc* is highest in the MZ1 and X clusters. MZ2 also shows relatively high *Myc* expression compared to the IZ, proPL, and PL clusters. **(D)** AUC scores for Srl target genes and Myc-activated genes are positively correlated in MZ1 and MZ2 cells (r=0.83). **(E)** Ribosome biogenesis genes (AUC) are highly enriched in MZ1 and MZ2. **(F)** Flow cytometric analysis of *dome^MESO^-GAL4, UAS-EGFP, Hml^Δ^-DsRed* lymph glands shows that on average, MZ cells show a higher forward scatter (FSC-A; a measure of cell size) than the cells of the CZ.

**Figure 3—figure supplement 4:**
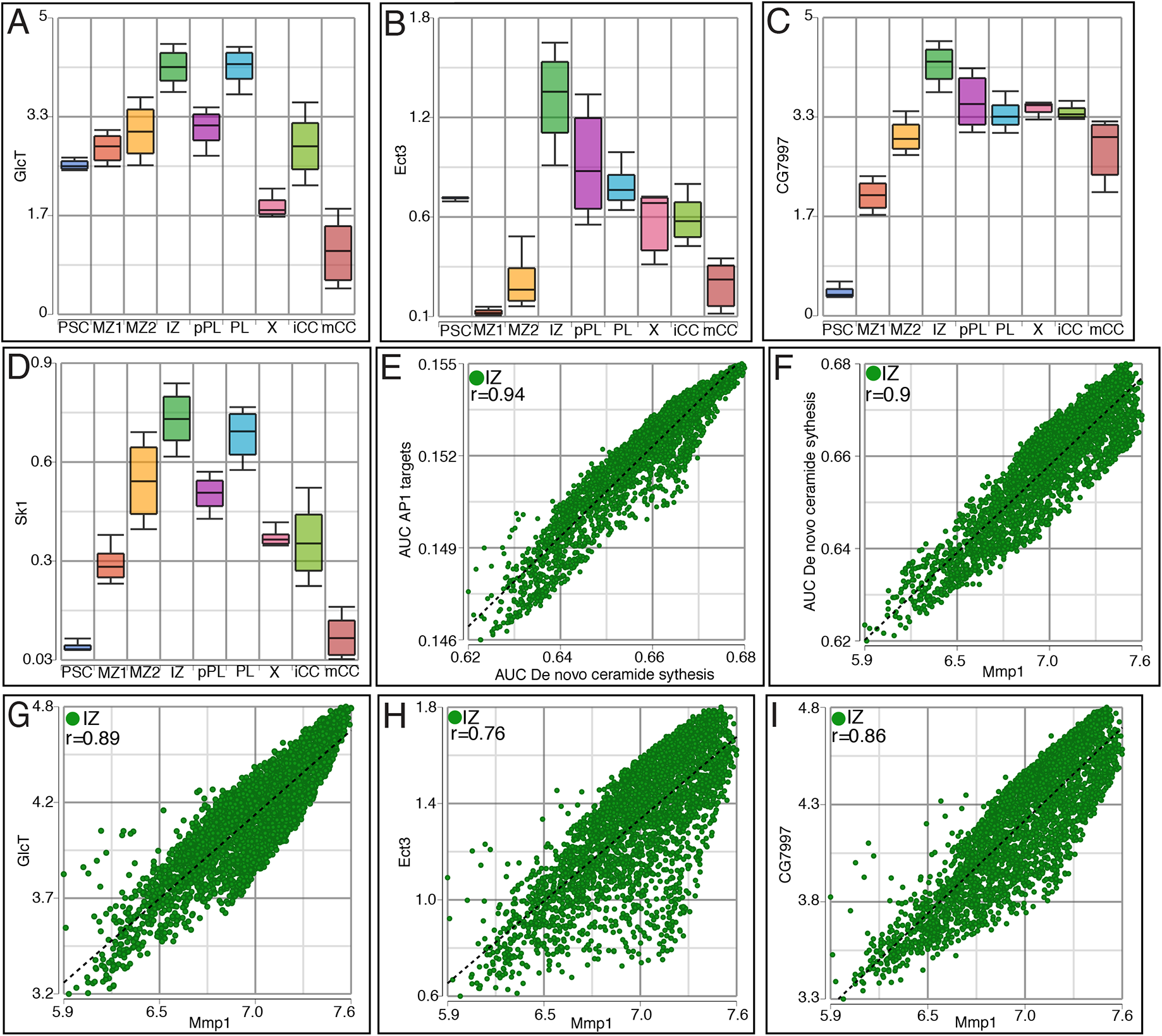
Sphingolipid metabolism pathways are enriched in the IZ. All data are based on single cell RNA-Seq analysis. **(A-D)** Genes involved in sphingolipid metabolism that are highly represented in the IZ include *GlcT* **(A),** *Ect3*/*Beta-gal* **(B)**, *CG7997*/*alpha-Gal* **(C)**, and *Sk1* **(D)**. **(E-H)** Positive correlation within the IZ is seen in comparisons of AP-1 transcriptional targets with *de novo* ceramide synthesis pathway activity **(E**; r=0.94), *de novo* ceramide synthesis pathway activity with the expression of the JNK transcriptional target *Mmp1* **(F**; r=0.9), and *MMP1* expression with that of three glycosphingolipid related genes: *GlcT* **(G**; r=0.89), *Ect3* **(H**; r=0.76) and *CG7997* **(I**; r=0.86)

**Figure 5—figure supplement 1:**
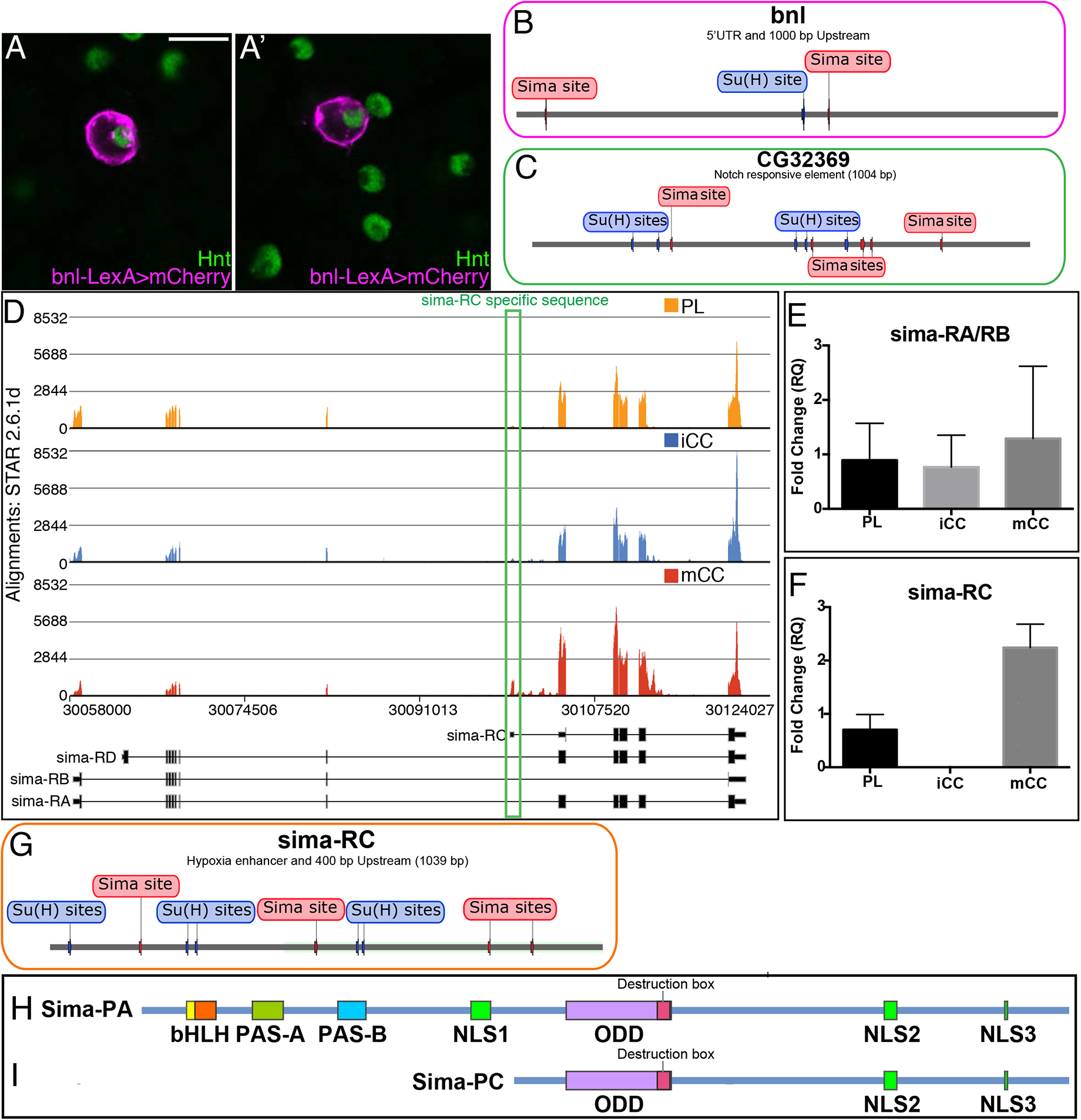
Notch/Sima related gene expression in mature crystal cells. **(A-A’)** *bnl-LexA* driving mCherry (magenta) marks a subset of Hnt+ (green) CCs. Scale bar, 10 µm. **(B)** Sequence analysis of the *bnl* promoter region (5’ UTR and 1000 bp upstream of the start site) shows potential Sima sites near potential Su(H) sites. **(C)** Sequence analysis of the Notch response element region of *CG32369* shows multiple potential Sima and Su(H) binding sites. **(D)** *Sima* transcript is alternatively spliced (*sima* gene model and predicted splice variants from flybase (r6.22) are indicated). *sima-RC* is specifically expressed in mCCs (lower track) with higher alignment counts than in PL (upper track) and iCCs (middle track) while *sima-RA* splice variant is highest in PL. **(E-F)** qRT-PCR analysis shows **(E)** no difference in *sima-RA*/*RB* transcript levels between PL and CC populations, while **(F)** *sima-RC* transcript levels are significantly higher in mCC than in iCC and PL. **(G)** The enhancer region of *sima-RC* contains multiple putative Sima and nearby Su(H) binding sites. **(H-I)** Schematic representation of predicted protein domains in Sima-PA **(H)** and Sima-PC **(I).** Sima-PC lacks the N-terminal bHLH and PAS regions required for DNA binding and dimerization with Tango but retains the ODD destruction box region that regulates protein stability under hypoxic conditions.

**Figure 5—figure supplement 2:**
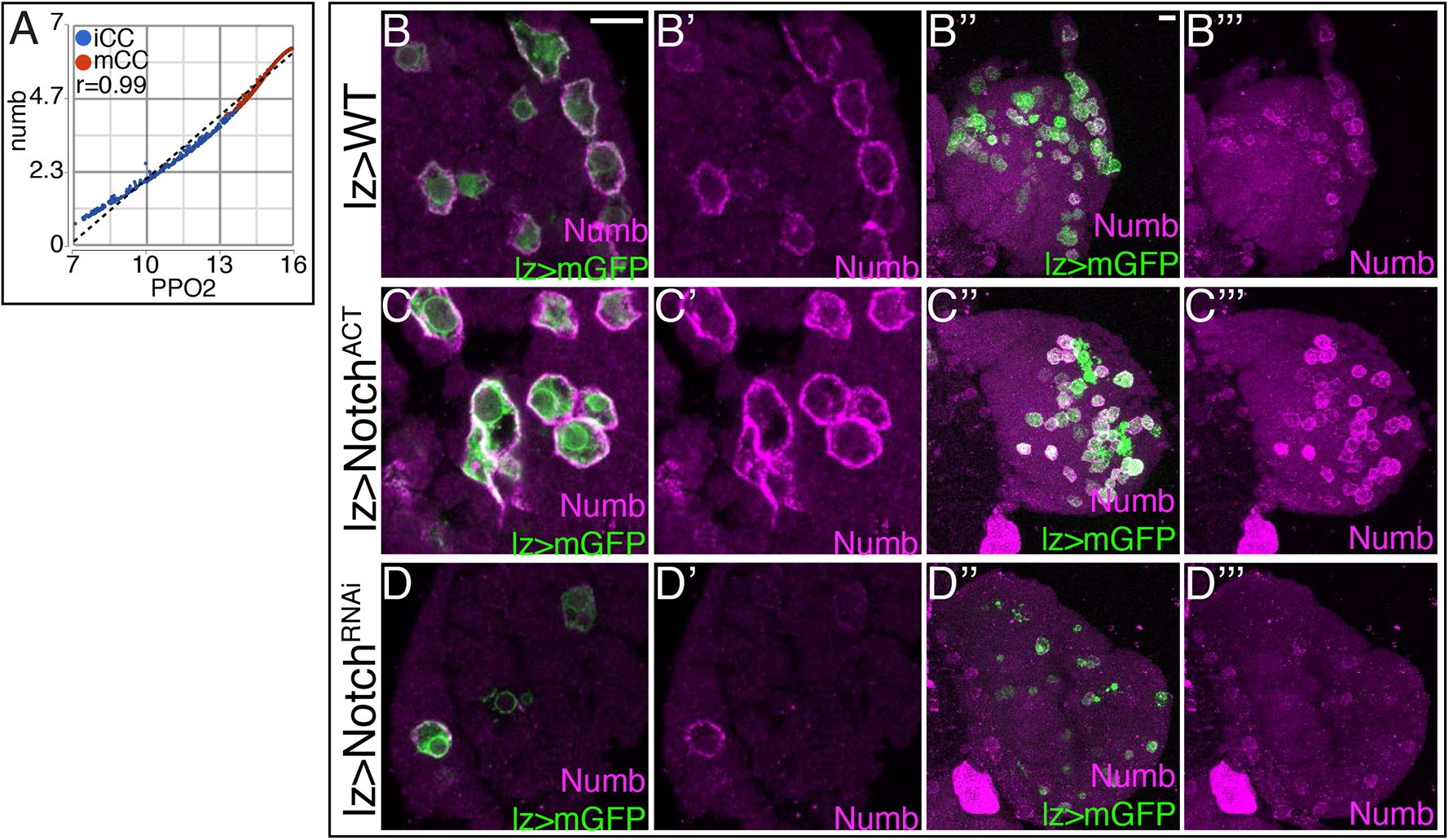
Numb levels are controlled by Notch signaling. **(A)** n*umb* transcript levels increase with maturity in both iCCs and mCCs and its expression strongly correlates with that of *PPO2* (r=0.99). Genotype in **B-D’’’**: *lz-GAL4*, *UAS-mGFP.* **(B-B’’’)** Wild-type lymph gland showing moderate Numb levels (magenta) in most *lz-GAL4, UAS-mGFP* expressing CCs (green). **(C-C’’’)** Overactivation of Notch signaling with *lz-GAL4*, *UAS-Notch^ACT^* greatly increases Numb levels (magenta) in CCs (green). **(D-D’’’)** Loss of Notch signaling with *UAS-Notch^RNAi^* both decreases the number of CCs (green) and decreases Numb levels (magenta) in the remaining CCs. Scale bars, 10 µm.

**Figure 5—figure supplement 3:**
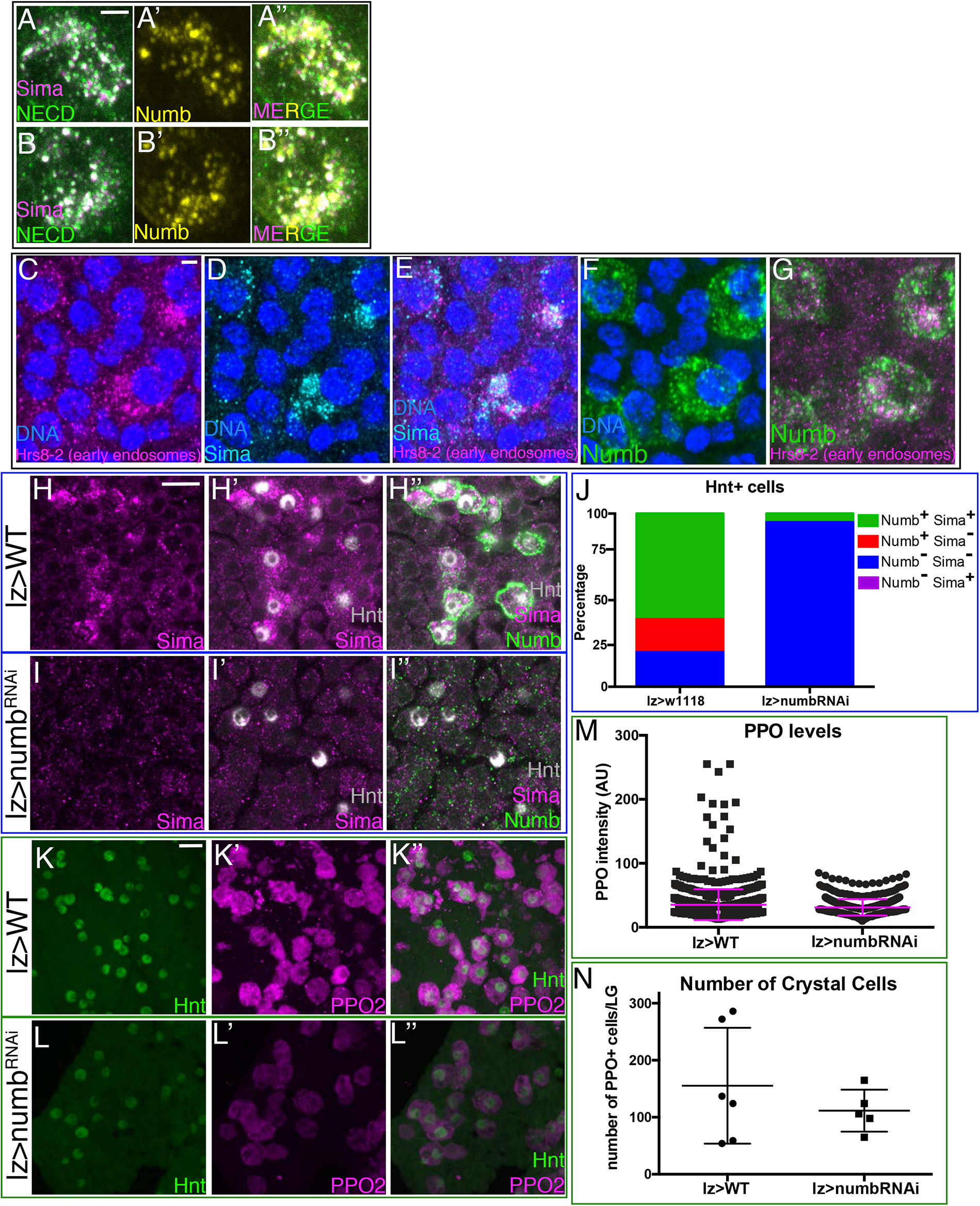
Numb promotes Notch/Sima signaling and CC maturity. **(A-B”)** Two example cells showing partial co-localization of Numb punctae (yellow; **(A’-A’’)** and **(B’-B’’)**) with Sima (magenta) and N^ECD^ (green). Punctae appear white in **(A)** and **(B)** due to colocalization of magenta and green. **(C-G)** Large Sima punctae (cyan) colocalize with Hrs8-2 positive early endosomes (magenta) in Numb+ crystal cells (green). DNA (blue). **(H-N)** Genotype, *lz-GAL4*. **(H-H”)** Large Sima punctae (magenta) in Numb+ (green) Hnt+ (grey) crystal cells are seen in wild-type lymph glands. **(I-I’’)** Sima punctae (magenta) are virtually eliminated when *numb* (green) is depleted from crystal cells (marked by Hnt, grey) with *lz-GAL4* driving *UAS-numb^RNAi^*. **(J)** Quantification of the percentage of Hnt+ crystal cells that are positive or negative for Sima and Numb in wild type and upon knockdown of *numb*. No Numb-Sima+ cells are evident in either genotype. Depletion of *numb* causes loss of nearly all Sima+ CCs. **(K-K’’)** PPO2 levels (magenta) are high in most Hnt+ (green) crystal cells in wild-type lymph glands. **(L-L’’)** PPO2 levels (magenta) decrease in Hnt+ (green) crystal cells upon depletion of *numb* with *numb^RNAi^.* **(M)** Quantification of PPO2 levels in individual cells show loss of virtually all high PPO2 expressing crystal cells upon loss of *numb* using *lz-GAL4* driving *UAS-numb^RNAi^*. **(N)** The total number of PPO+ crystal cells does not change with *numb^RNAi^*. **(A-G)** Scale bars, 2 µm. **(H-I, K-L)** Scale bars, 10 µm.

**Figure 5—figure supplement 4:**
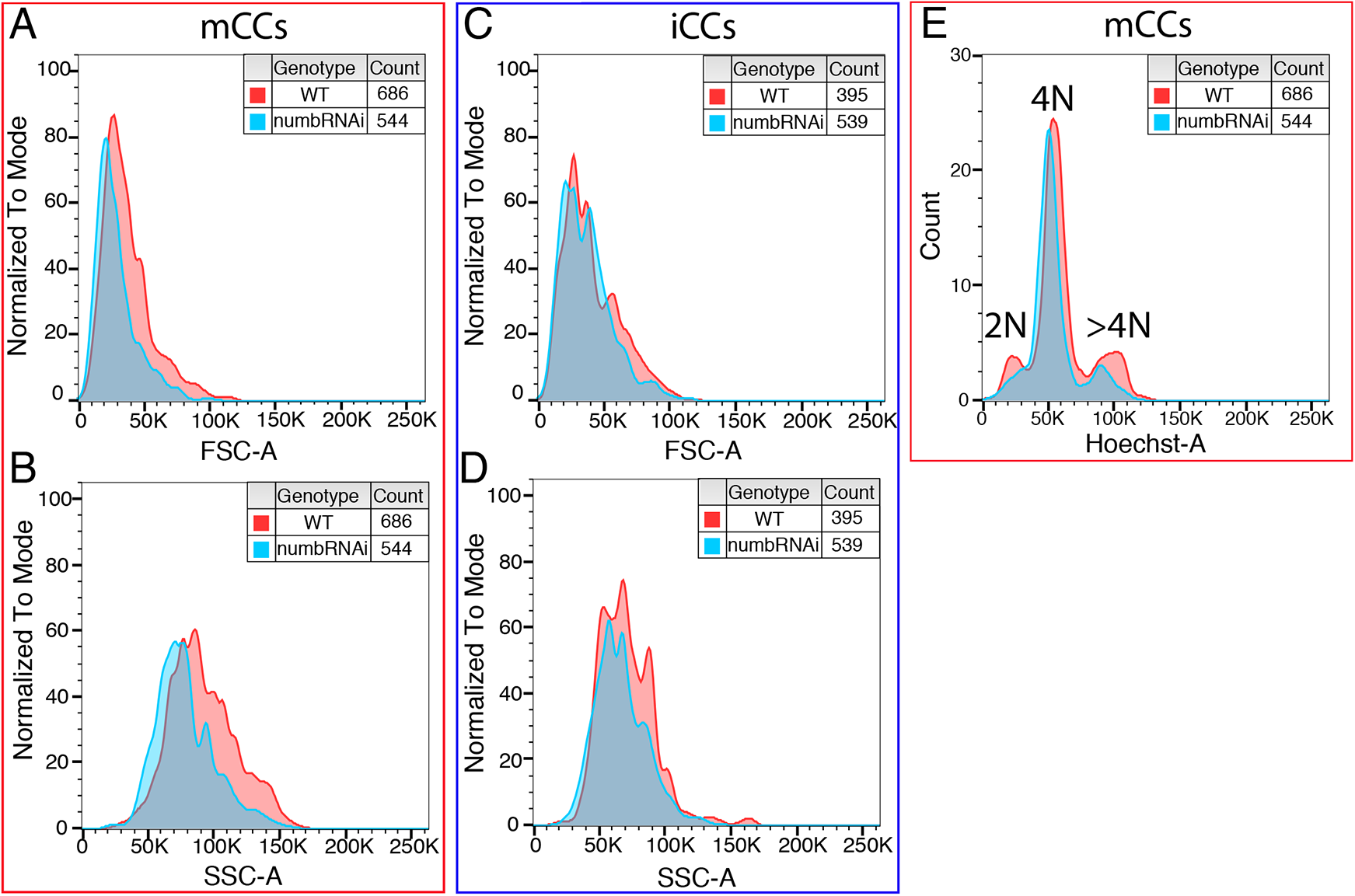
Flow cytometric analysis of CC subclasses from *lz-GAL4* wild-type and *lz-GAL4, UAS-numb^RNAi^* lymph glands. **(A-B)** Depletion of *numb* decreases cell size (Forward Scatter; FSC-A) and cellular complexity (Side Scatter; SSC-A) of mCCs. **(C-D)** Loss of *numb* does not change cell size or cellular complexity of iCCs. **(E)** The number of mCCs with >4N DNA content is reduced upon *numb^RNAi^* expression.

**Figure 6—figure supplement 1:**
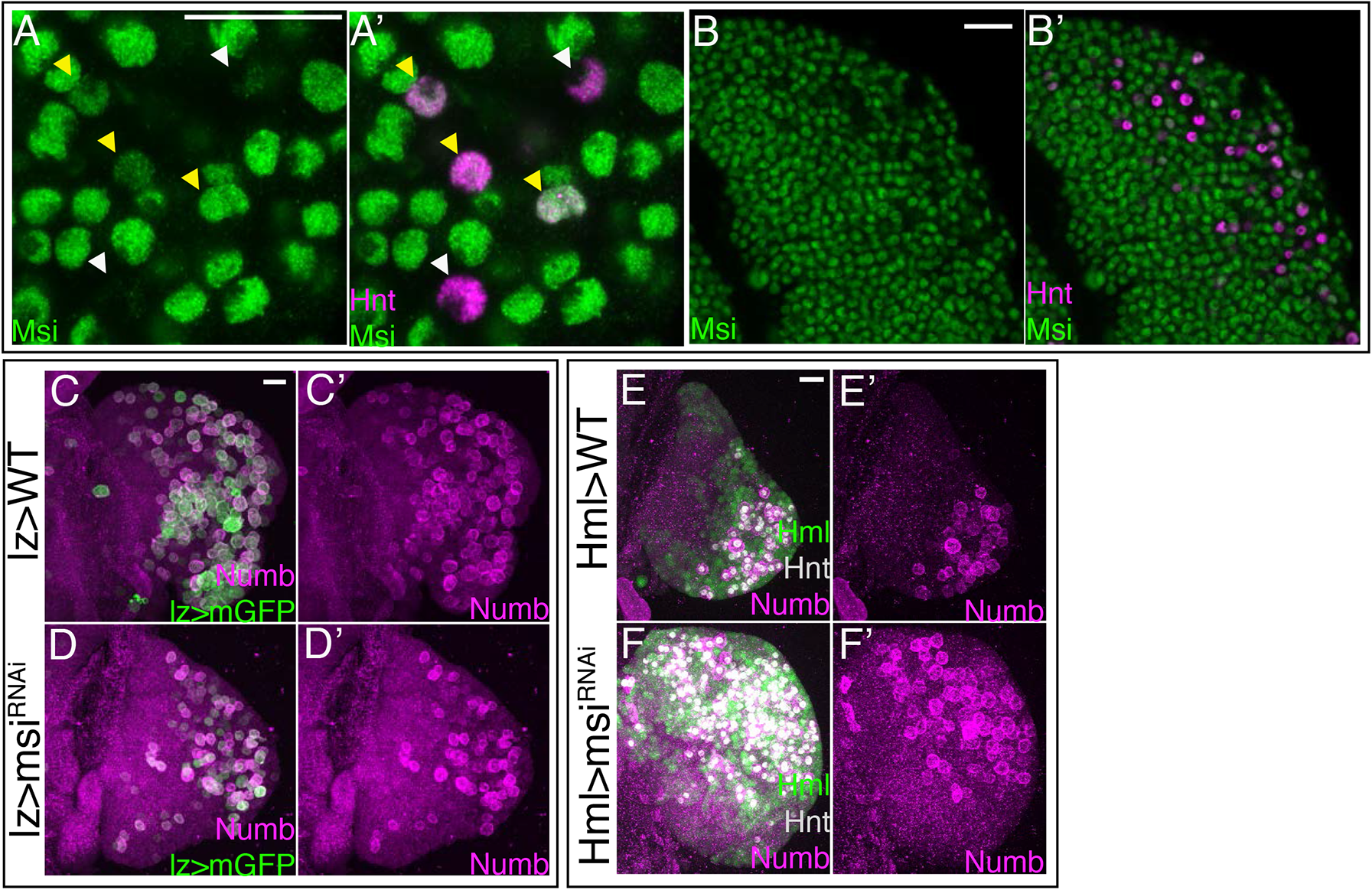
Numb protein level is controlled by Musashi. **(A-B’)** Wild-type lymph gland expressing GFP-tagged Msi (green) and stained for Hnt (magenta) shows that Msi is high in most cells, except in a subset of CCs with the highest Hnt levels (white arrowheads). Yellow arrowheads denote Hnt+ CCs with high Msi-GFP. Shown in high **(A,A’)** and low **(B,B’)** magnification. **(C-F’)** Lower magnification view showing middle third maximum intensity projections of the entire lymph gland lobes from Figure 6H (C-C’); Figure 6I (D-D’); Figure 6K (E-E’); and Figure 6L (F-F’). Scale bars, 20 µm.

