## Supplementary file 2 for "Paths and Pathways that Generate Cell-Type Heterogeneity and Developmental Progression in Hematopoiesis"

**Page 1. Figure legend.**
**Supplementary File 2**: Comparison of clusters from this study with those identified as clusters by Cho et al., 2020.

**Page 2: (A-F)** To create a uniform platform for comparison of clusters identified in this study with that by Cho et al., 2020, we used AUCell to map enrichment of the gene lists from each of the Cho et al., 2020 clusters onto the cells within our t-SNE visualized clusters. We find that genes enriched in:
**(A)** PSC$^{\text{Cho et al.}}$ shows high expression in our PSC cells,
**(B)** PH$^{\text{Cho et al.}}$ shows high expression in MZ1 and MZ2 cells,
**(C)** CC$^{\text{Cho et al.}}$ shows high expression in iCC and mCC cells,
**(D)** PM$^{\text{Cho et al.}}$ shows high expression in IZ, proPL, and PL cells,
**(E)** Adipohemocyte$^{\text{Cho et al.}}$ shows high expression in IZ, a subset of proPL, and mCC cells,
**(F)** GST-rich$^{\text{Cho et al.}}$ shows high expression in most of our clusters.
**(G-H)** Cho et al. (2020) use intermediate levels of expression of the four genes, *Tep4*, *Ance*, *Hml*, and *Pxn* (the first two are MZ markers, the last two are CZ markers) as a reasonable measure of cells that belong to an intermediary state. Following the same method for identifying intermediate expression of the above four genes from our study, we find that
**(G)** Cells expressing these levels of the genes map at the boundary between cluster MZ2 and the proPL, IZ, and PL clusters.
**(H)** Remarkably, this combination of cells are largely confined to State 3 of our trajectory suggesting that this process of selection identifies transitory states rather than zones.
**Abbreviations:** For clusters identified in this study, the abbreviations are as defined in the main text. To avoid confusion over nomenclature, the superscript "Cho et al." is added to each of the clusters assigned in the corresponding paper. For example, PSC$^{\text{Cho et al.}}$ (Posterior Signaling Center), PH$^{\text{Cho et al.}}$ (Prohemocytes), CC$^{\text{Cho et al.}}$ (Crystal Cells), PM$^{\text{Cho et al.}}$ (Plasmatocytes).

**Page 3:** Map of subclusters identified by Cho et al. to our t-SNE visualization plot as on Page 2.
**(A-L)** AUCell of gene lists from Cho et al. subclusters usually map to more than one cluster from this study.
**(A-F)** PH$^{\text{Cho et al.}}$ subclusters 1-6.
**(G-J)** PM$^{\text{Cho et al.}}$ subclusters 1-4.
**(K-L)** CC$^{\text{Cho et al.}}$ subclusters 1-2.

Cluster gene lists from Cho et al. mapped onto scRNA-Seq data from this study

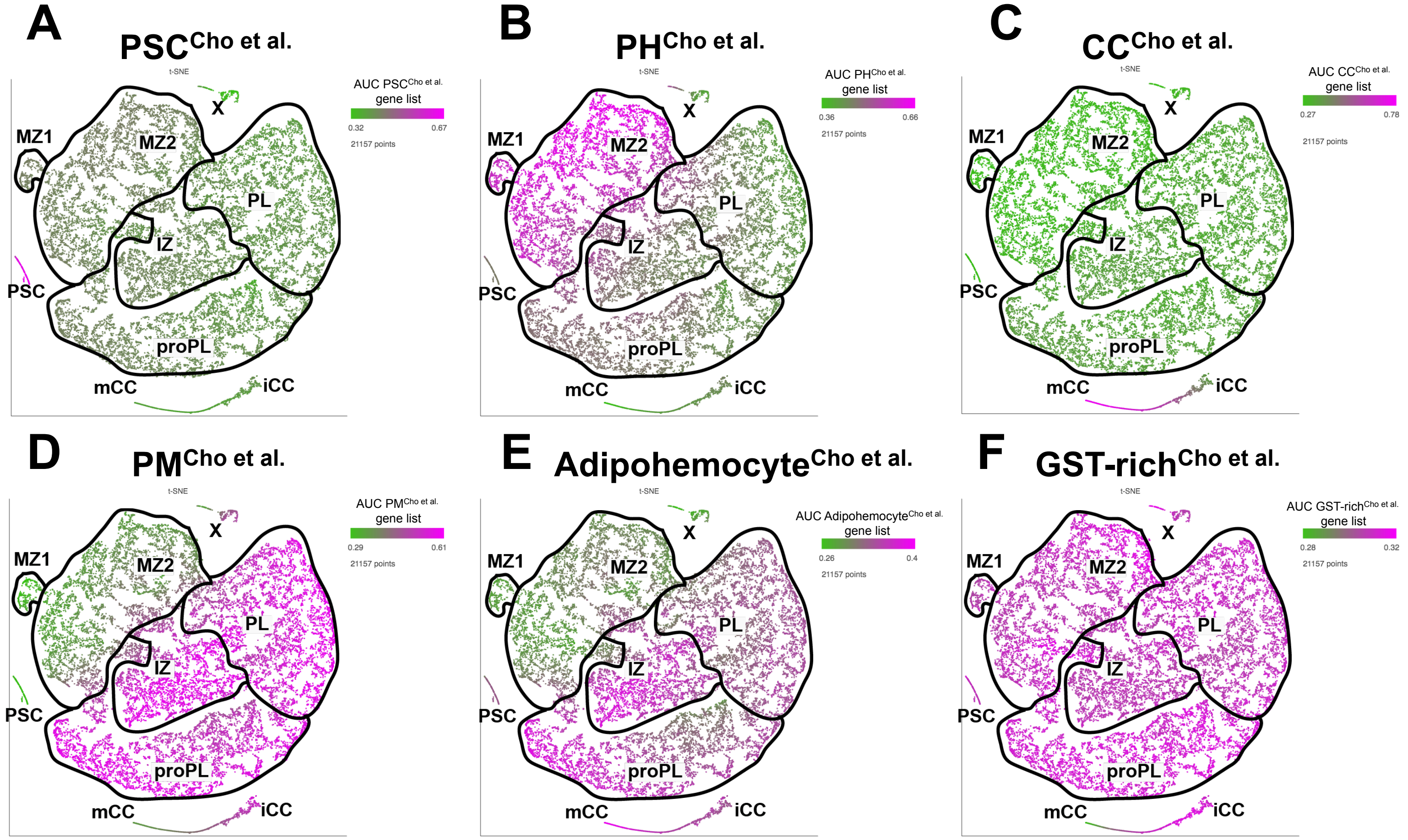


Zone of intermediate *Tep4*, *Ance*, *Hml*, and *Pxn* expression

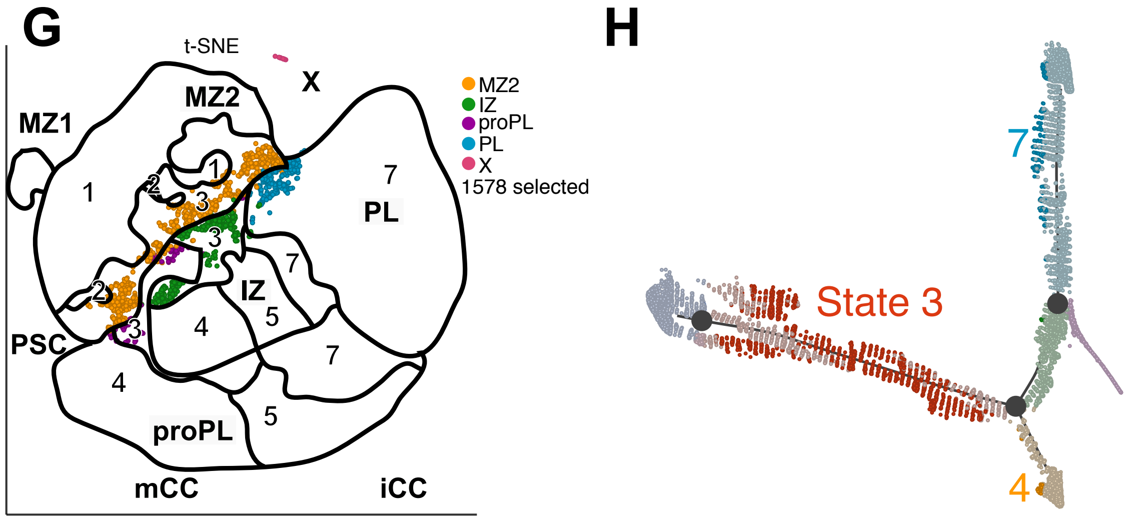

Subcluster gene lists from Cho et al. mapped onto scRNA-Seq data from this study

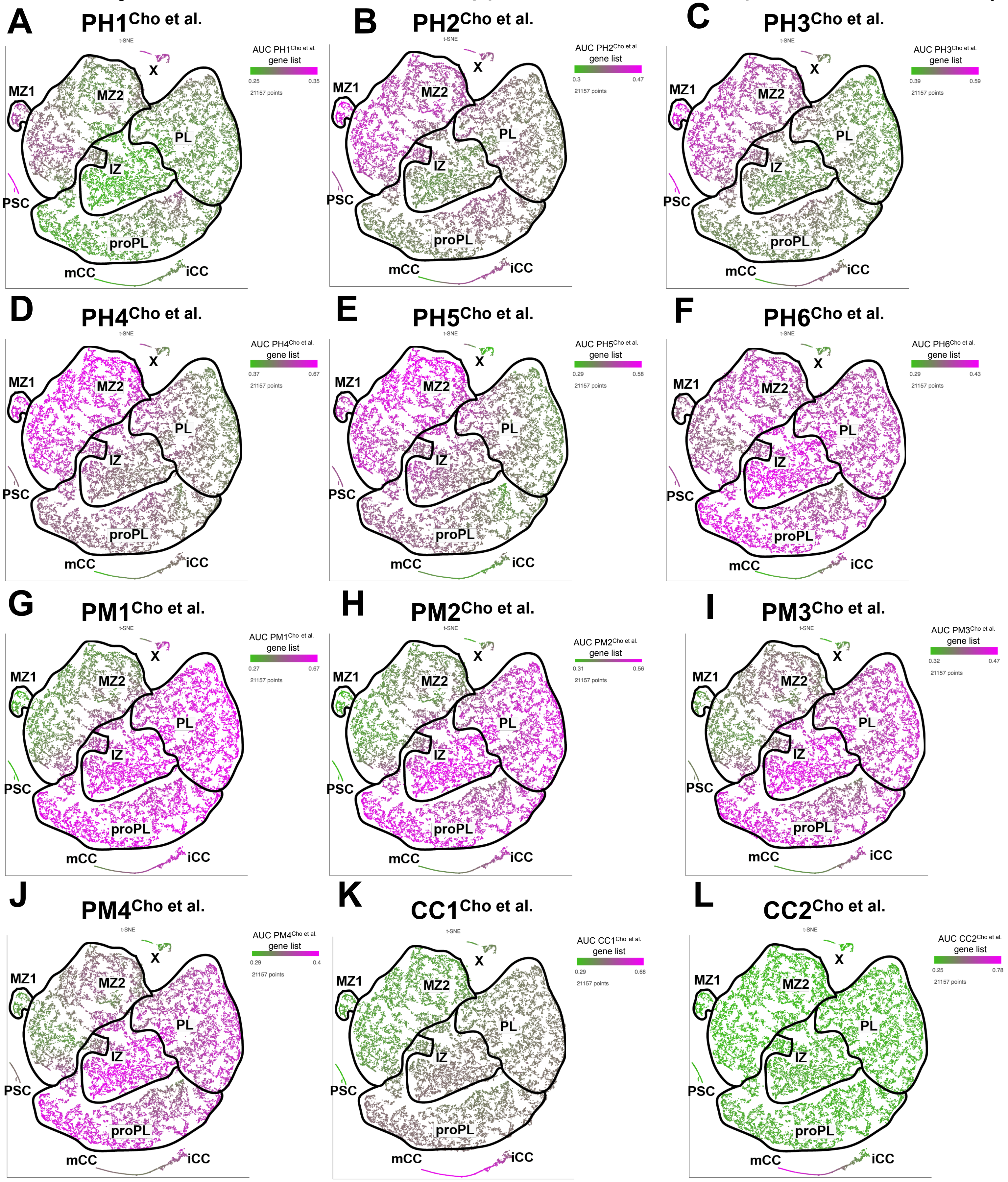
